# *De novo* designed 3-helix bundle peptides and proteins with controlled topology and stability

**DOI:** 10.1101/2025.06.30.662276

**Authors:** Xiyue Leng, Katherine I. Albanese, Lia R. Golub, Arthur A. Norman, Jonathan Clayden, Derek N. Woolfson

## Abstract

Computational protein design is advancing rapidly. However, approaches and methods are needed to increase success rates and to elaborate designs. Here we describe the combination of rational and computational design to deliver three-helix bundle (3HB) peptide assemblies and single-chain proteins with control over topology and thermal stability. First, we garner sequence-to-structure relationships from antiparallel 3HBs in the Protein Data Bank. This gives core-packing rules, including layers of hydrogen-bonded polar residues, which are combined with surface-charge patterning to design complementary sequences for acidic (A), basic (B), and neutral (N) helices. By altering the design of the N helix, two sets of synthetic peptides are generated for clockwise and anticlockwise arrangements of the three-helix assemblies. Solution-phase characterization shows that both ABN peptide mixtures form stable, heterotrimeric assemblies consistent with the targeted ‘up-down-up’ topologies. Next, AlphaFold2 models for both designs are used to seed computational designs of single-chain proteins where the helices are connected by loop building. Synthetic genes for these express in *E. coli* to yield soluble, monomeric, and thermally stable proteins. By systematically introducing additional polar layers within the core, the thermal stability of these proteins is varied without compromising the specificity of the helix-helix interactions. Chemical and thermal denaturation reveals comparable thermodynamic parameters to those of highly stable natural proteins. Four X-ray crystal structures confirm that the design models and AlphaFold2 predictions match to sub-Å accuracy.

## 1. INTRODUCTION

Protein design is advancing rapidly, increasingly using computational and artificial intelligence (AI) methods.^1–3^ Despite recent advances, a deeper understanding of sequence-to-structure/function relationships remains essential to deliver designs that are more predictable, yield higher experimental success rates, and are generally more fit for purpose.

To reach this point, *de novo* protein design has undergone several phases, which we can learn from.^4–7^ Historically, minimal protein design employed fundamental chemical principles—such as sequence patterns of hydrophobic and polar residues—to render mimics of simple, natural protein folds. Aided by developments in bioinformatics, rational design emerged, integrating analyses of natural protein sequence and structural databases, such as the Research Collaboratory for Structural Bioinformatics Protein Data Bank (RCSB PDB)^8, 9^, into *de novo* design. This has led to improved sequence-to-structure relationships to inform better designs.^4^ As the field has matured, computational methods have increasingly been used in design pipelines, including parametric design to sketch out backbones, and physical forcefields to assess sequences that best fit these.^10, 11^ Now, AI-based models built from large sequence and structural datasets that capture relationships *en masse* are being used to generate backbones and sequences either separately or simultaneously.^1, 3, 6, 12, 13^

However, while AI-based methods can exploit both known and unknown relationships, they often lack interpretability, making it difficult to uncover how these relationships govern design outcomes. Despite recent success in delivering complex *de novo* protein scaffolds and functions,^14–17^ the explainability and success rates of AI-driven methods remain low. Therefore, to continue advancing the field, these gaps in understanding and applicability need to be filled. This does not necessarily mean replacing ‘black-box’ methods. Rather, they need to be augmented to add reasoning and insight to enhance their predictive power, efficiency, and robustness. In short, we would like to achieve *fully programmable protein design* based on understanding sequence-to-structure/functional relationships in proteins. *En route* to this, we advocate combining rational and computation protein design for targets when possible leading to more explainable AI for *de novo* protein design.

α-Helical coiled coils (CCs) have been particularly fruitful targets for *de novo* protein design due to many well-established principles that link primary sequences to a variety tertiary and quaternary structures.^4, 18, 19^ As such, they are well suited to the challenge of combining rational, computational, and AI-based approaches in protein design. Generally, CC structures have two or more right-handed α helices supercoiled around one another to form usually left-handed helical bundles. These are encoded by 7-residue (heptad) sequence repeats denoted (*abcdefg*)_n_. The *a* and *d* sites are predominately occupied by hydrophobic residues, giving the pattern (*hpphppp*)_n_. When configured into an α helix, this produces a hydrophobic seam that drives helix association, and sets a framework for specifying helix orientation (parallel or antiparallel), oligomeric states (dimer and above), partner preferences (homo- or heteromeric), and overall topology and handedness of the tertiary or quaternary structure as described below.^18, 20^ In addition, the introduction of electrostatic or polar interactions flanking the hydrophobic core—usually at the *e* and *g* sites for dimers to tetramers, or *b* and *c* sites for pentamers and above—can be used to define CC assemblies further.^21, 22^ Combined, established combinations of amino acids at the *g-a-d-e* positions can be used to generate toolkits of *de novo* CC assemblies.^7, 23, 24^ In turn, these peptide assemblies can be repurposed to design functional peptides for catalysis^25, 26^, materials assembly^27–30^, and in cell and synthetic biology^31, 32^. Most recently, some of the peptide sequences and experimental 3D structures have been used as “seeds” to deliver more-complex, single-chain proteins through computational protein design leveraging the new AI-based methods.^33, 34^

Here, we focus on CC assemblies of three α helices, termed 3-helix bundle (3HB) CCs. These are appealing due to their relative simplicity and small size, making them ideal for studying structure, folding, and stability, and, therefore, for developing further fundamental design rules and principles^35–37^ and new applications^38–40^. The homotrimeric peptide Coil-Ser provides an early example of an ‘up-down-up’ 3HB architecture—*i.e.*, with one helix aligned antiparallel to the other two—discovered serendipitously in an attempt to design a parallel dimeric CC.^41^ Notable progress has also been made towards desymmetrizing 3HBs using attractive ion pairs at the interfacial positions, effectively guiding the specific formation of heterotrimeric assembly over alternative oligomeric states or homo- or mixed-trimers.^42–44^ DeGrado and colleagues have advanced this by developing an iterative design transitioning Coil-Ser-derived sequences to native-like globular proteins.^35^ Recent innovations include using buried hydrogen bonds and shape-complementary packing to create highly specific heterotrimers that serve as biological scaffolds.^45^

We build on this foundation here to describe the design of a series of 3HB CC peptide assemblies and single-chain proteins with up-down-up arrangements of three helices but with different and complementary sequences; that is, an ABN system composed of acidic (A), basic (B), and neutral (N) strands. By strategically positioning interhelical charges, we can dictate the overall conformational handedness of the 3HBs, achieving either clockwise (CW) or anticlockwise (ACW) topologies in solution. Models for each of these peptide assemblies are then used to seed the computational design of single-chain 3HB proteins, which are expressed from synthetic genes in *E. coli*, and confirmed by X-ray structures with the intended topologies and handedness of the tertiary structures. By introducing polar residues into the hydrophobic core, we fine-tune thermal stability without compromising folding specificity. This study demonstrates the feasibility of designing 3HB CCs with precise control of topology and thermal stability by combining rational, computational, and AI-based *de novo* design, and shows how design success rates can be improved through such pipelines.

## 2. RESULTS AND DISCUSSION

### 2.1 Rational design of up-down-up heterotrimeric α-helical peptide assemblies

To garner sequence-to-structure relationships for the target, we searched the CC+ database of structurally validated CC proteins for examples of up-down-up 3HBs (Tables S1 and S2).^46^ This returned 169 examples of antiparallel heterotrimeric peptide assemblies and 3-helix protein bundles with up-down-up connectivity. This was from a total of 285 3-helix CCs in the database. Manual assignment of the subset of 169 structures revealed a strong natural bias: 121 had the anticlockwise topology, ans 48 had the clockwise arrangement. Notably, in single-chain proteins, the predominant anticlockwise topology has right-handed helical connectivity consistent with the left-handed supercoiling of CCs. Given the underrepresentation of clockwise topologies in nature and the associated design challenge, we initially prioritized this configuration. The helical sequences of the 169 subset were used to compile a 20 × 7 amino-acid profile from their component *a* – *g* heptad repeats (Figures 1A and B). The SWISS-PROT normalized data revealed preferences for certain amino acids at the different heptad positions, which were used as rules to design peptide sequences.

**Figure 1.**
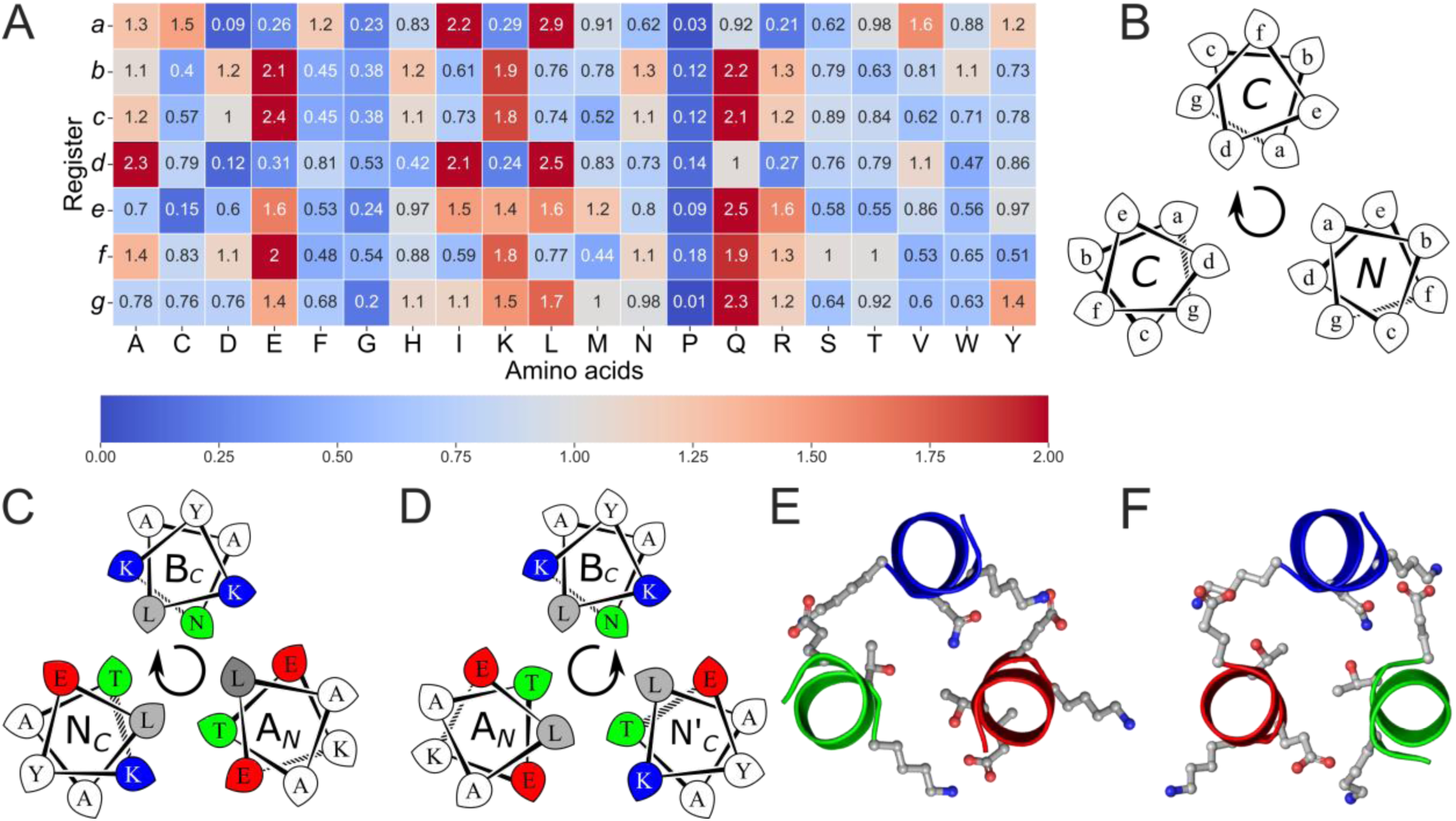
The rational design of new sequences to form antiparallel CC heterotrimers. (A) Amino-acid propensities for each position of the heptad repeats for 3HB CCs pulled from the CC+ database.^46^ Raw counts (Table S1) were normalized using the amino-acid frequencies in SWISS-PROT to provide the propensity scale shown as a heat map (high to low: red to blue). (B) Helical-wheel diagram for a clockwise antiparallel trimeric assembly, showing heptad assignments and with the helical termini closest to the viewer labelled. (C&D) Helical-wheel representations of apCCTri-BĀN (clockwise) and apCCTri-BĀN’ (anticlockwise), respectively. Sequences have canonical heptad repeats, *abcdefg*. The residues at key positions of the designs, *g, a, d,* and *e*, are highlighted: gray for hydrophobic, Leu; green for polar, Thr and Asn; red for acidic, Glu; and blue for basic, Lys. (E&F) Slices through the third heptad of the AF2 models for apCCTri-BĀN and apCCTri-BĀN’, respectively, with the key side chains shown as sticks. For B – D, arrows indicate the overall handedness of the quaternary structure. This is defined as follows: with the first helix at 12 o’clock and coming out towards the viewer, the assemblies have either a clockwise (B-Ā-N) or an anticlockwise (B-Ā-N’) arrangement.

The core-defining *a* and *d* positions were made Leu, as this was the most preferred residue at both sites. The core-flanking *e* and *g* positions were made combinations of Glu and Lys, as they featured highly at these sites and offered possibilities for directing helix-helix partnering through electrostatic interactions.^18, 47^ Building on previous work^42, 43^ to design away from homomeric assembly and to favor heteromeric association, we designed one sequence to be acidic (A) with *e* = *g* = Glu, and a second basic (B) peptide with *e* = *g* = Lys. Typically, such AB systems are designed for even oligomers as the alternating charges can be satisfied by C2, C4 or D2 symmetry.^31, 48–50^ Therefore, to target an ABX-type heterotrimer, the third helix was made neutral (N) with *e* = Glu plus *g* = Lys to give complementary interactions to both flanking acidic and basic helices (Figure 1C). As noted previously by DeGrado and co-workers,^35^ helical wheels suggests an alternative design for the assembly with the opposite (anticlockwise) cyclic order of the three helices, which can be achieved by switching the polarity *e* = Lys and *g* = Glu giving an alternative neutral peptide (N’) (Figure 1D). To help specify the antiparallel arrangement further, specifically with the A helix antiparallel (denoted Ā) to the B and N helices, we incorporated a layer of polar residues at *d-a-a* sites of the Ā-B-N/N’ combinations in the otherwise hydrophobic cores (Figures 1E and F).^22, 51, 52^ This was chosen by inspecting the CC+ dataset derived above for buried, three-residue constellations of polar side chains. This revealed STAT5a (1y1u), which has an ordered, hydrogen-bonded network between Thr-155 (*a*), Thr-236 (*d*) and Asn-289 (*a*).^53^ This polar layer was compatible with both the clockwise (CW, BĀN) and anticlockwise (ACW, BĀN’) target assemblies. The designs were completed as 4-heptad sequences with *b* = *c* = Ala to provide specificity for the assembly, and combinations of polar and aromatic residues at the *f* sites to enhance solubility and to introduce chromophores (Tables 1 and S3).

**Table 1.**
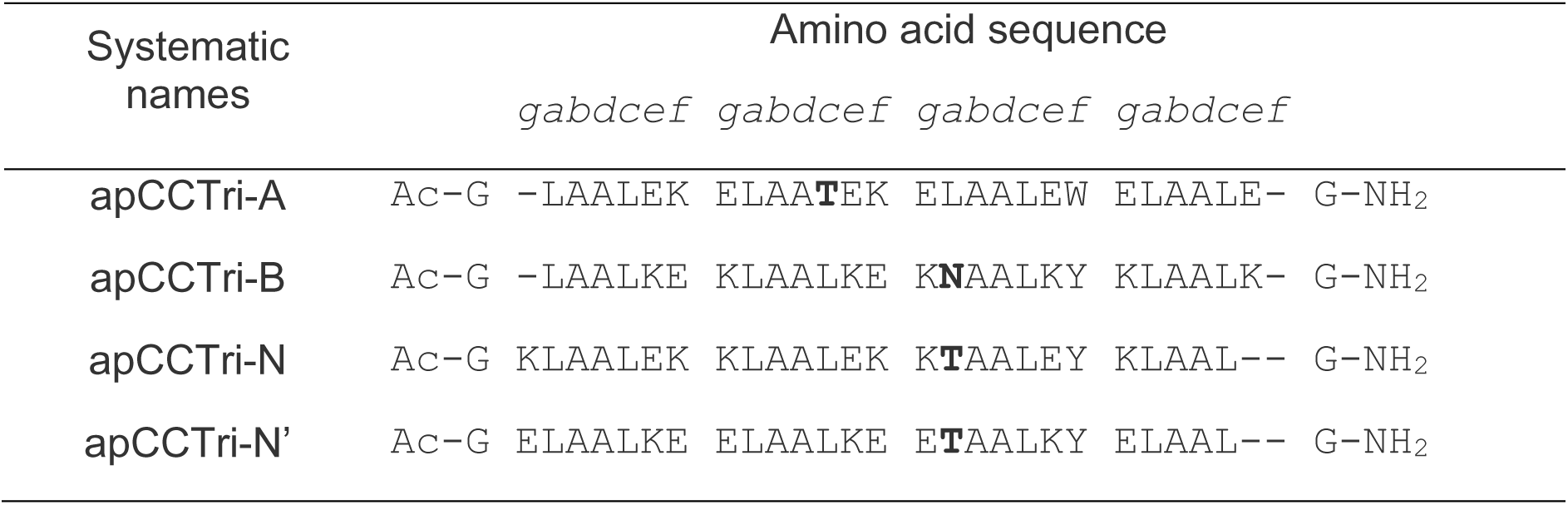
Designed sequences of apCCTri-A, B, N and N’ peptides.

Ahead of experimental work, all combinations of the A, B, and N and A, B, and N’ sequences were modelled using AlphaFold2-multimer (AF2, Table S4). The predicted models were consistent with the target heterotrimeric assemblies with the designed helical topologies, *i.e.*, mixed parallel and antiparallel helices and clockwise or anticlockwise peptide assembly. The target and alternate state models (AAA, AAB, ABB, *etc*) were assessed by predicted template modelling (pTM) and local distance difference test (pLDDT) scores. However, the average pLDDT scores were all above 95%, so, effectively, AF2 did not discriminate between the models (Table S4).

### 2.2. Biophysical characterization of synthetic peptides confirms the design targets

All four peptides A, B, N, and N’ were synthesized by Fmoc solid-phase peptide synthesis, purified by reverse-phase HPLC, and verified by MALDI-TOF mass spectrometry (Table S3, Figures S1). The peptides were characterized in solution individually and as 1:1:1 mixtures as described below.

First, circular dichroism (CD) spectroscopy was used to probe the folding and thermal stability of the peptides at 100 µM concentration in phosphate buffered saline at pH 7.4 (PBS; Figure 2A). In contrast to AF2 predictions, peptide A was completely unfolded under these conditions. Peptide B was partially folded at 5 °C but unfolded readily with increased temperature (midpoint of thermal denaturation, T_M_ < 15 °C). These behaviors are consistent with the design hypothesis, as highly charged peptides are not expected to self-associate appreciably. By contrast, the neutral, peptides N and N’ were highly helical and thermally stable (T_M_ ≈ 78 and 54 °C, respectively). The difference in the T_M_ values can be explained by the order of Glu and Lys residues in the sequence.^21^ However, both mixtures BĀN and BĀN’ were also highly helical and highly thermally stable (T_M_ ≈ 73 and 67 °C, respectively, Figure S2). For completeness, data for the pairwise combinations were collected and compared with the respective theoretical averages, showing reduced helicity and cooperative folding (Figure S2, Table S6). The solution-phase oligomeric states for the stable complexes, N_3_, N’_3_, BĀN and BĀN’, were determined by sedimentation-velocity (SV) and sedimentation-equilibrium (SE) experiments in analytical ultracentrifugation (AUC) at 20 °C (Figures 2C and D, Table S6, Figures S3 and S4). These revealed monodisperse trimeric species as designed.

**Figure 2.**
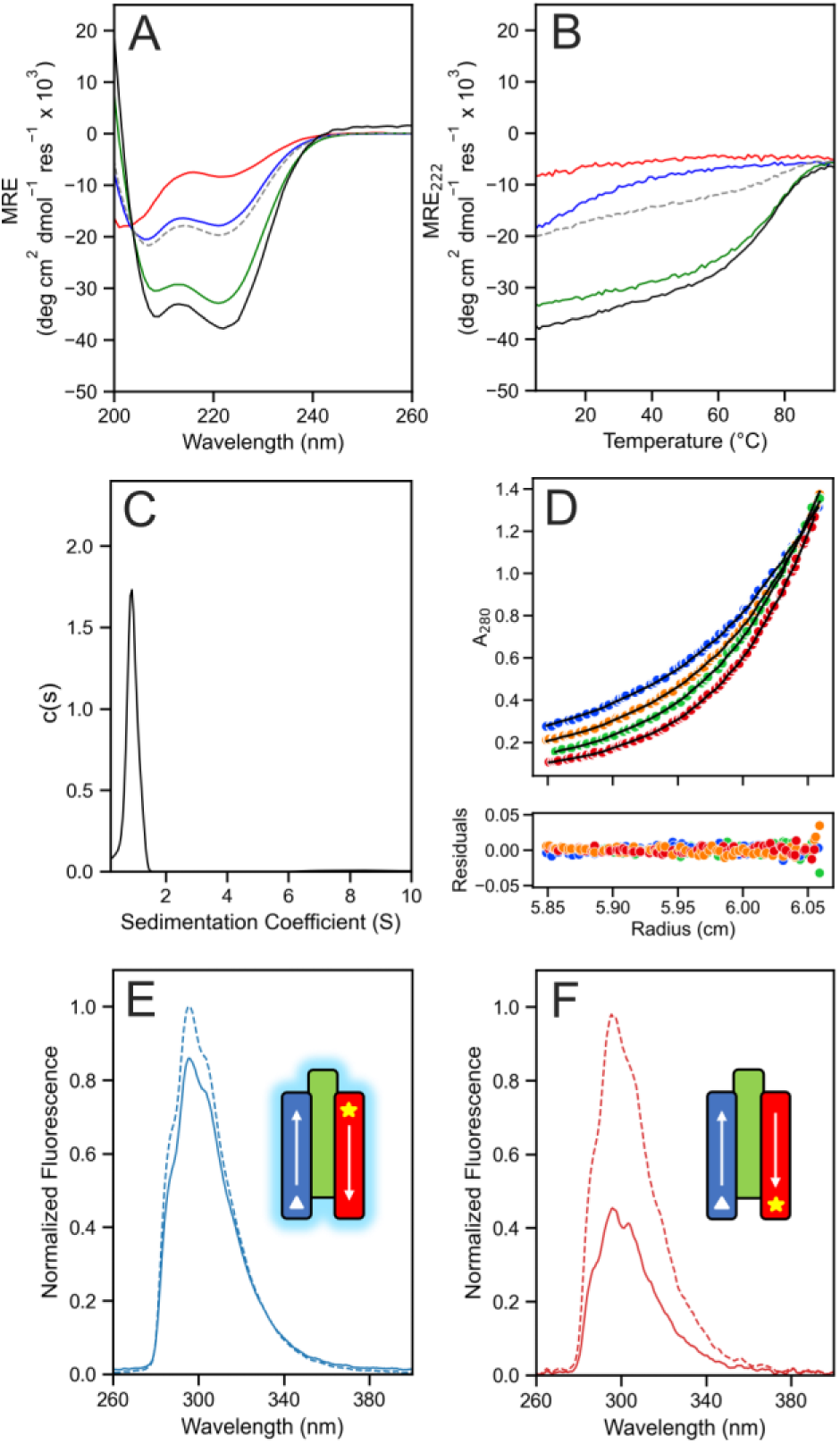
Biophysical characterization of the apCCTri A, B and N peptides. (A) CD spectra at 5 °C, and (B) thermal responses of the CD signals at 222 nm (ramping from 5 to 95 °C at 1 °C min^−1^) of the individual peptides (A, red; B, blue; and N, green), the equimolar 1:1:1 mixture (black), and the calculated theoretical averages (dashed). Conditions: 100 µM in 1 × PBS, pH 7.4. (C) AUC-SV and (D) AUC-SE profiles of apCCTri-BĀN 1:1:1 mixture. Fits returned molecular weights consistent with a trimeric assembly (1.1 and 0.9 × the summed masses, respectively). Conditions: 250 µM in 1 × PBS, pH 7.4; SV run at 60 krpm, SE run at 40 – 52 krpm. Key: 40 krpm, blue; 44 krpm, orange; 48 krpm, green; and 52 krpm, red. (E&F) Fluorescence-quenching assays for mixtures of apCCTri-B-nMSE and apCCTri-N plus apCCTri-A-n4CF (E) or apCCTri-A-c4CF (F). Dotted lines are for the fluorescently-labelled A peptides alone, and solid lines are for the 1:1:1 mixture. In the cartoons, arrows indicate the peptide direction from *N* to *C* termini; 4CF is represented by the yellow star and MSE by the white triangle. Conditions: 33 µM of each peptide in phosphate buffer (8.2 mM potassium phosphate dibasic, 1.8 mM potassium phosphate monobasic), pH 7.4.

Despite multiple attempts, we could not crystallize any of the species formed. Therefore, we used a fluorescence-quenching assay to probe helix orientation in the BĀN and BĀN’ systems in solution.^54^ First, guided by the AF2 models, we placed spatially proximal fluorophore (4-cyanophenylalanine, 4CF) and quencher (L-selenomethionine, MSE) pairs on two different peptides. For instance, the B peptide was remade with an *N*-terminal MSE, apCCTri-B-nMSE; and two variants of the A peptide were made with 4CF at the *N*-terminal *g* site or the *C*-terminal *e* site, apCCTri-A-n4CF and apCCTri-A-c4CF, respectively. As expected for an antiparallel arrangement of A and B helices, no quenching of 4CF fluorescence was seen when apCCTri-B-nMSE and apCCTri-A-n4CF were mixed with the unlabeled N peptide (apCCTri-N), Figure 2E. However, quenching did occur with the other mixture with apCCTri-A-c4CF, Figure 2F. This demonstrated that the A helix was oriented antiparallel to B in solution as designed. To probe the orientation of the N helix, we carried out the analogous experiments using the two 4CF-labelled A peptides, unlabelled B, and with the N peptide *N*-terminally labelled with MSE, apCCTri-N-nMSE. This gave similar results to those shown in Figures 2E and 2F indicating that the N helix also aligns antiparallel to A and, therefore, parallel to B (Figure S5).

Following this solution-phase characterization, and consistent with our systematic naming of *de novo* CC peptide,^18^ we named the two heterotrimeric peptide assemblies apCCTri-BĀN and apCCTri-BĀN’.

### 2.3 Peptide assemblies can be used as seeds for single-chain protein designs

Recently, we introduced the concept of rationally seeded design of single-chain proteins from characterized peptide assemblies.^33, 34^ This takes computational or experimental structural models of the peptide assemblies and uses computational design to link the *C* and *N* termini of neighboring helices. To do this here, we used MASTER to search the RCSB PDB for loops to connect helices in the AF2 models for the clockwise, apCCTri-BĀN, and anticlockwise, apCCTri-BĀN’, topologies.^55, 56^ Models for these were built using AF2 in single-sequence mode, and assessed by pLDDT and root mean square deviation (RMSD) from the peptide models (Tables S4 and S5). Next, ProteinMPNN was used to optimize the loop regions of the single-chain models while keeping the helical seeds completely intact. Based on pLDDT and RMSD, two BĀN and one BĀN’ single-chain designs were chosen to test experimentally.

Synthetic genes for the sequences were expressed in *E. coli* (Table S3). As the parent peptide assemblies were thermally stable, the cell lysate was heat-shocked at 65 °C for 10 min. Immobilized metal affinity column chromatography (IMAC) and size exclusion chromatography (SEC) were used to yield purified proteins (Figures S6). CD spectroscopy of all proteins showed that they were highly α helical and hyperthermally stable, and AUC experiments confirmed they were monomeric in solution (Figures 3A – E).

**Figure 3.**
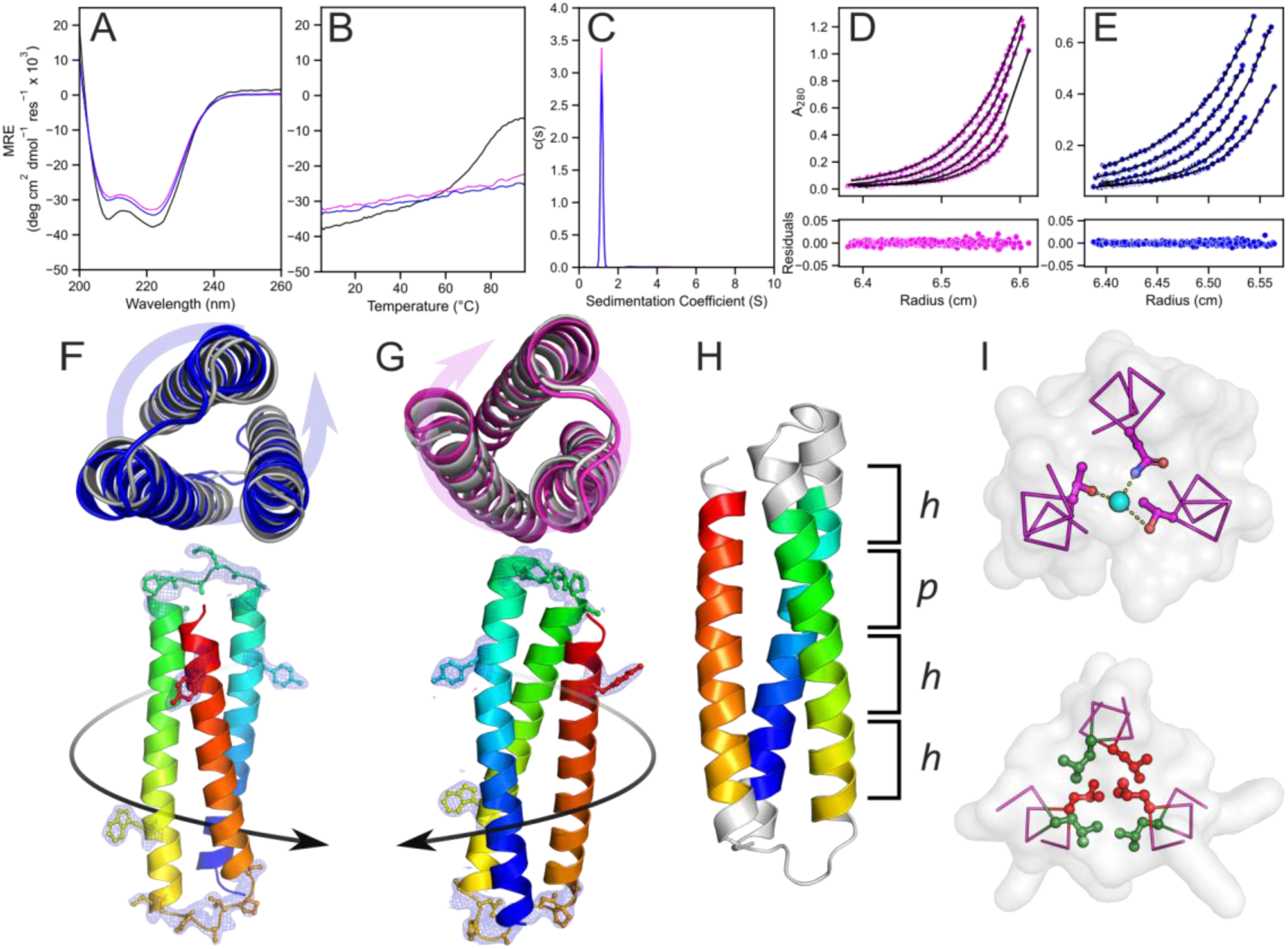
Rationally seeded computational designs of the single-chain 3HB proteins based on models for the apCCTri-BĀN peptide assemblies. Two different designs were made and characterized, sc-apCC3-CW (data and images in magenta) and sc-apCC3-ACW (blue). (A) CD spectra at 5 °C and (B) thermal responses of the CD signals at 222 nm (ramping from 5 to 95 °C at 1 °C min^−1^). Conditions: 10 µM protein, PBS, pH 7.4. Data for the apCCTri-BĀN peptide assembly are included for comparison (black). (C) AUC-SV and (D&E) AUC-SE data. Fits returned molecular weights of 0.91 and 0.84 × monomer mass for sc-apCC3-CW, and 0.87 and 0.77 × monomer mass for sc-apCC3-ACW, respectively. Conditions: 50 µM protein, PBS, pH 7.4; SV run at 60 krpm, and SE run at 44 – 60 krpm. (F&G top) Top views of the X-ray crystal structures of sc-apCC3-CW1 (G; 2.10 Å, 9rgv), and sc-apCC3-ACW (F; 2.20 Å, 9rgx) aligned with AF2 model (rank 1, gray) to Cα-RMSD of 1.079 and 1.742 Å, respectively. Arrows indicate the overall handedness of the tertiary structure. This is defined as follows: with the N-terminal helix at 12 o’clock and coming out towards the viewer, the chain trace follows either a clockwise or anticlockwise path. (F&G bottom) Side views of the same structures with each helix identified by electron densities of the loop region and the Trp and Tyr side chains, which are contour level (*σ*) = 1, 2mFo-DFc maps displayed to within 1.3 Å to the region of interest. (H) X-ray crystal structure of sc-apCC3-CW2 (2.15 Å, 9rgw) with CC regions identified by Socket2^57^ colored as chainbows, KIH packing cutoff is 7 Å. (I) Slices through the structure at the third heptad repeat with the water-mediated polar layer (top), and the second heptad showing KIH packing of the consolidated Leu core, with *a* knobs in red and *d* in green (bottom).

We solved X-ray crystal structures at 2.10 Å, 2.15 Å and 2.20 Å resolution for the two BĀN- and one BĀN’-based designs, respectively (Figure 3F – I, Tables S7 and S8). This was done by molecular replacement using the AF2 models of the peptide assemblies as the search models. All confirmed the three-helix bundles with up-down-up topologies for the B-Ā-N/N’ helices. All had consolidated hydrophobic cores with knobs-into-holes packing between the core Leu residues confirmed by Socket2 (Figure 3H – I).^57^ In addition, water-mediated hydrogen-bonded layer of Thr (A) – Asn – Thr (N/N’) was present in a clockwise design (Figure 3I). Moreover, the conformational handedness of the two structures was different and as designed. Therefore, we named these proteins sc-apCC3-CW (for the clockwise BĀN combination, 9rgv and 9rgw) and sc-apCC3-ACW (for the anticlockwise BĀN’, 9rgx) to form part of the growing toolbox of *de novo* CC peptide assemblies and single-chain proteins.^18, 33, 34^

### 2.4 The relative importance of helix-helix interactions and connecting loops

Next, we tested the importance of loop design in our two parent designs. To do this, we swapped the loops between the two topologies: the clockwise bundle was furnished with the computationally designed loops for the anticlockwise design, and *vice versa*, to give sc-apCC3-CW-mismatch and sc-apCC3-ACW-mismatch, respectively. In addition, we substituted the designed loops for flexible glycine (Gly, G) plus serine (Ser, S) linkers in both topologies, using six- and eight-residue GS linkers to bridge the gaps rendering sc-apCC3-CW-flex and sc-apCC3-ACW-flex, respectively. The sequences for these redesigns are given in Table S3.

All four modified constructs expressed well in *E. coli*, were monomeric, α helical, and thermally stable (Figures S6 – 9). However, the CD spectra revealed a progressive decrease in helicity from the parent design (Figures S7 and S10), with a ≈10% reduction for the constructs with flexible linkers and a further ≈15% drop for those with the mismatched loops compared (Table S9, Figure S10). This suggests that even with optimized core designs—which are sufficient to drive the correct assembly of the ABN-peptide assemblies—loop optimization is important for single-chain protein design. Moreover, at least for our design target, flexible linkers are better than using mismatched loops.

Despite extensive trials, crystals could not be obtained for any of the flexible or mismatched constructs. Therefore, AF2 was used to predict models for all variants. Consistently, this gave high-confidence models (Table S4). Moreover, secondary structure analysis predicted high helical contents of ≈ 73 – 78%, ^58, 59^ which is higher than the experimentally observed values (Figures S7 and S10, Table S9). This indicates that, while AF2 confidently predicts local secondary and overall structure extremely well, it does not capture subtleties in sequence-to-structure relationships. That said, inspection of the predicted models revealed that some lower-ranked model for sc-apCC3-CW-mismatch had the anticlockwise topology, hinting that loop mismatching may permit access to alternative folds and that AF2 may well capture this (Figure S11).

### 2.5 Tuning the thermal stability of sc-apCC3 through polar layers in the core

As we have demonstrated previously^33, 34^ and above, the application of robust and well-understood design rules and the rationally seeded design pipeline yields exceptionally stable CC-based proteins with control over handedness of protein topology. Hyper-stable proteins have advantages for applications requiring resilience to harsh conditions or as structural scaffolds. However, extreme stability may not always be desirable in biological systems—particularly in processes that rely on dynamics, such as ligand binding, conformational switching, and enzymatic catalysis. For example, natural CC domains found in heat shock proteins provide a balance between stability and responsiveness, exhibiting structural flexibility and enabling conformational changes to assist protein folding under conditions of stress.^60^

To alter thermal stability in our designs without compromising the 3HB fold and topology, we sought to destabilize sc-apCC3-CW1 systematically by modifying its consolidated, largely Leu-based hydrophobic core. For this section, this parent design with a single polar layer of Asn@*a*, Thr@*d*, and Thr@*d* (N-T-T) in the B, A, and N helices (Figure 1) is referred to as sc-apCC3-CW-1NTT. As hydrophobic core packing is a key stabilizing force in water-soluble globular proteins^61^, we investigated whether introducing additional polar layers could attenuate core stability without compromising the overall 3HB structure.

Additional polar layers were introduced sequentially at the second, first, and fourth heptads to give sc-apCC3-CW-2NTT, sc-apCC3-CW-3NTT, and sc-apCC3-CW-4NTT, respectively (Figure 4A). This progression ensures that the central regions of the protein are destabilized first and remodelled with polar interactions, while maintaining fully hydrophobic and stabilizing heptad repeats at the termini. As a control, we included a variant without a polar layer, namely sc-apCC3-CW-0NTT. All sequences were evaluated using AF2, which consistently predicted clockwise 3HB topologies with high confidence (Table S4). The variants with 0 and 2 N-T-T layers crystallized, and the X-ray crystal structures aligned with the parent design, with Cα RMSDs of 1.015 Å and 0.605 Å, respectively (Tables S7 and S8; pdb ids 9rgy, 9rgz). The variants with 3 and 4 N-T-T layers could not be crystallized.

**Figure 4.**
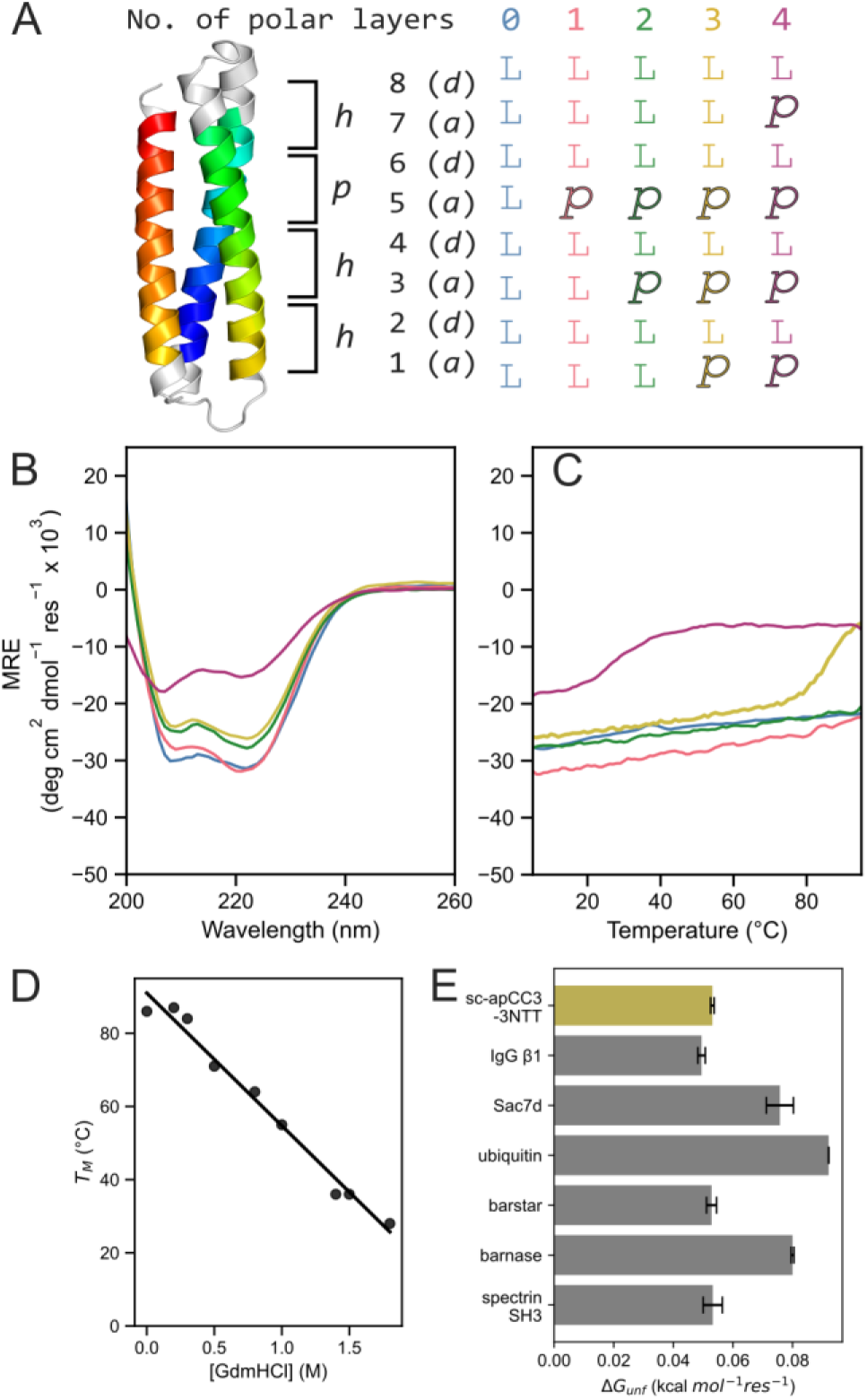
Systematic destabilization of sc-apCC3-CW with multiple polar layers in the core yields proteins with accessible unfolding transitions. (A) Design schematic for the designs with 0 – 4 polar layers, sc-apCC3-CW-“X”NTT. Key: X = 0, blue; 1, pink; 2, green; 3, yellow; and 4, purple. (B) CD spectra recorded at 5 °C. (C) Thermal responses of the CD signals at 222 nm (ramping from 5 to 95 °C at 1 °C min^−1^). Conditions: 10 µM protein in 1 × PBS, pH 7.4. (D) T_M_ as a function of GdmHCl concentration. A linear regression y = −36.25x + 90.99 is fitted to the data (R² = 0.98). (E) Comparison of per-residue Gibbs free energy of unfolding (Δ*G*_unf_) with natural proteins of similar size.

These observations were supported by the CD spectroscopy. CD spectra showed a progressive decrease in helicity with the addition of N-T-T layers (Figures 4C and S7, Table S6). The most extreme case was sc-apCC3-CW-4NTT, which retained only ∼50% helicity relative to the 1NTT and 0NTT variant. Variable-temperature CD measurements revealed that sc-apCC3-CW with 0, 1, and 2 N-T-T layers were hyperthermally stable and did not unfold upon heating (Figure 4D). In contrast, sc-apCC3-CW-3NTT and -4NTT had measurable T_M_ values of 86.5 ± 0.8 and 25.3 ± 0.7 °C, respectively (Figures 4D and S7). Whilst sc-apCC3-CW-3NTT showed good reversibility from the pre- and post-melt CD spectra, sc-apCC3-CW-4NTT remained unfolded after heating and was disregarded from further analysis (Figure S7).

Thus, sc-apCC3-CW-3NTT is an example of a *de novo* designed single-chain protein with an accessible, reversible, and cooperative thermal-unfolding transition. This is unusual in contemporary designed single-chain proteins, which are often hyper-stable. Therefore, we characterized the thermodynamics of its unfolding in more detail.

The CD spectra and T_M_ values for sc-apCC3-CW-3NTT did not change with protein concentration (Figure S12), confirming that the protein folds as a non-associating monomer as designed. Given its appreciable thermal stability, we used guanidinium hydrochloride (GdmHCl) as a chemical denaturant to access melting transitions at lower T_M_ values for a full thermodynamic analysis (Figures S13 – 15). This gave well-defined, sigmoidal, thermal-unfolding curves, which were fitted to a two-state model to estimate T_M_ over a range of GdmHCl concentrations (Figures S13 – 15, Table S10).^62^

The resulting T_M_ values were linearly related to the GdmHCl concentration (Figure 4D). Encouraged by this, we sought to estimate the thermodynamics of unfolding at 0 M GdmHCl using the linear extrapolation method (LEM). Traditional van’t Hoff analysis assumes Δ*C*_p_ = 0. However, this neglects the substantial structural reorganization and exposure to solvent associated with protein unfolding, which results in a positive Δ*C*_p,unf_.^5, 63^ Therefore, we used a global nonlinear Gibbs-Helmholtz fitting procedure to model the free energy of unfolding (ΔG_unf_) as a function of temperature and GdmHCl concentration under the LEM (Table S10).^64, 65^ This gave Δ*G*_unf_ of 6.16 ± 0.07 kcal mol^−1^ at 25 °C in the absence of denaturant (Figure S15). This is consistent with the observed stability of the protein under those conditions. Moreover, when expressed per residue, it is comparable to the most stable natural globular proteins under similar conditions (Figures 5E and S16),^66–68^ most of which are not all-α-helical proteins. The modest change in enthalpy (Δ*H*_unf_ = 24.08 ± 0.53 kcal mol^−1^) and entropy (Δ*S*_unf_ = 0.06 ± 0.00 kcal mol^−1^K^−1^) reflect the small size of the 3HB. The small heat capacity change (Δ*C*_p_ = 0.09 ± 0.01 kcal mol⁻¹ K⁻¹) indicates opposing hydration effects upon burying polar (negative Δ*C*_p_) and nonpolar side chains (positive Δ*C*_p_), which partly cancel.^69–71^ Nonetheless the net value, along with its dependence on the denaturant (ΔΔ*C*_p,[GdmHCl]_ = 0.46 ± 0.01 kcal mol⁻¹ K⁻¹ M⁻¹), are consistent with a compact native state and the progressive solvation of the largely hydrophobic core upon unfolding.

## 3. Conclusion

In summary, we have developed sequence-to-structure relationships for the rational design of simple up-down-up, heterotrimeric, coiled-coil peptide assemblies. These relationships are informed by modelling and confirmed in solution using a variety of biophysical measurements. Two distinct topologies of the design target are generated by exploiting electrostatic interactions to specify the helix-helix interfaces. Next, AlphaFold2 models for the two assemblies are used to seed the computational design of single-chain three-helix-bundle proteins with clockwise and anticlockwise arrangements of the component helices. Synthetic genes for these designs express well in *E. coli* to give hyper-stable, water-soluble, monomeric proteins. High-resolution X-ray crystal structures for the two designs match the computational models with atomic accuracy, confirming the design rules and pipeline used. The structures reveal that rational design can precisely specify conformational handedness, reliably accessing the more-common anticlockwise configuration seen in natural helical bundles, but also the less-prevalent clockwise form. This ability to control of helix-helix interfaces firmly establishes another level of programmability in CC design.^35^

Miniprotein designs of the type delivered here provide relatively straightforward model systems to test relationships between designed sequence and structure, stability and function.^72–75^ This is essential for the field to move towards quantitative and fully programmable protein design. As part of this quest, here we show that in addition to sequence-to-structure relationships for the well-defined secondary and tertiary elements of the targeted fold, loop design is important. Specifically, for connecting adjacent elements of secondary structure, design-specific loops appear to be better than flexible linkers, which are better than mismatched loops. In addition, we show that the hyperstability often observed with modern *de novo* designed scaffolds can be attenuated through robust rational. In our case, the core packing of the parent design consists of eight layers, with three residues in each layer. All but one of these layers comprise solely leucine residues. The remaining layer is a hydrogen-bonded constellation of one asparagine and two threonine side chains. By introducing two more of these layers, such that 3/8 of the layers and 9/24 of the core residues are polar, the protein unfolds reversibly below 100 °C in aqueous buffer. Detailed analysis of experimental unfolding data reveals that the modified design has thermodynamic parameters comparable to natural proteins of similar size. This targeted approach adds to and complements the emerging quantitative understanding of stability determinants from high-throughput studies of other *de novo* miniproteins.^75^

We posit that further systematic analyses of this and similar *de novo* proteins will help to uncover sequence-to-structure/stability relationships and advance fully predictive and quantitative *de novo* protein design.^74–76^ In addition, as we^25, 77^ and others^40, 78^ have demonstrated, small *de novo* peptide assemblies and single-chain proteins of the type delivered here provide robust scaffolds to generate functional *de novo* proteins by grafting or otherwise introducing functional residues such as binding and catalytic sites. Regardless of how functionalization is introduced—*i.e.*, rationally, computationally, or generatively with AI—an advantage of using predesigned and well-characterized scaffolds is that the roles and positions of most residues are known and understood at the atomistic level, providing a robust foundation for designing, introducing, and controlling functional modifications.

## Associated content

### Supporting Information

Experimental materials and methods, CC+ search results, peptide and protein sequences, purification data, biophysical and structural characterizations, and thermodynamic data and fits (PDF).

The raw data and code used in this publication have been deposited at Zenodo (10.5281/zenodo.15620429)

## Author contributions

X.L. and D.N.W. conceived the study and designed the peptide sequences and experiments. X.L., K.I.A. and L.R.G. synthesized and characterized the peptides. X.L. designed and characterized the proteins. A.A.N. contributed to the design concept for distinguishing bundle handedness. X.L. collected the X-ray data and determined the protein X-ray crystal structures. J.C. and D.N.W. provided supervision and sourced funding. X.L. and D.N.W. wrote the paper. All authors have read and contributed to the preparation of the manuscript.

## Notes

The authors declare no competing financial interest.

## Acknowledgements

X.L. is supported by an Engineering and Physical Sciences Research Council (EPSRC)-funded PhD studentship through the Centre for Doctoral Training in Technology Enhanced Chemical Synthesis (EP/S024107/1). K.I.A. was supported by a Biotechnology and Biological Sciences Research Council (BBSRC)–National Science Foundation grant (BB/V004220/1 and 2019598). We are also grateful to the Max Planck-Bristol Centre for Minimal Biology, which supports K.I.A. and D.N.W.. J.C. is supported by ERC through Advanced Grant DOGMATRON (883786). We thank the Mass Spectrometry Facility for access to the EPSRC-funded Bruker UltraFlex MALDI-TOF instrument (EP/K03927X/1) at School of Chemistry, University of Bristol. We thank Diamond Light Source for access to beamlines I03, I04, I24 (proposals mx 37593 and mx31440). We further thank N.J. Savery for scientific discussions and co-supervision of A.A.N.. Finally, we thank R. Petrenas for assistance with determining the X-ray crystal structures and members of the Woolfson and Clayden laboratories for many helpful discussions.

## Table of Content graphic

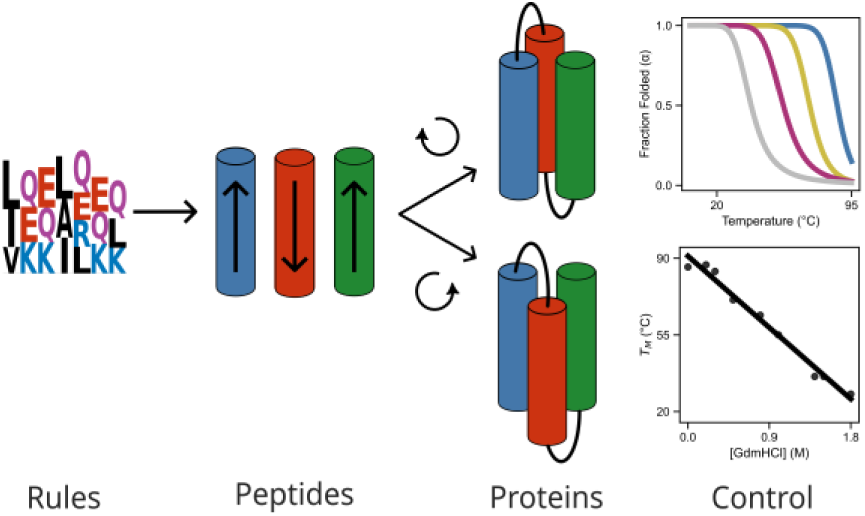

## Notes

https://10.5281/zenodo.15620429

### Competing Interest Statement

The authors have declared no competing interest.

