## supplementary information for "*De novo* designed 3-helix bundle peptides and proteins with controlled topology and stability"

### Methods

#### 1.1 General

All solvents, chemicals, and reagents were purchased from commercial sources and used without further purification. Fluorenylmethoxycarbonyl(Fmoc)-α-L-amino acids, Rink amide MBHA resin for solid-phase peptide synthesis (SPPS) and N,N-dimethylformamide (DMF) were purchased from Sigma-Aldrich and Cambridge Reagents. Coupling reagents Oxyma Pure and diisopropylcarbodiimide (DIC) were purchased from Fluorochem and Sigma-Aldrich, respectively. Morpholine and trifluoroacetic acid (TFA) were purchased from Sigma-Aldrich; pyridine was from Thermo Fisher; triisopropylsilane (TIPS) was from Acros Organics. Luria-Broth (LB), antibiotics, IPTG, and L-rhamnose were purchased from Thermo Fisher. All other chemicals were reagent grade and purchased from Sigma-Aldrich.

All reference made to clockwise (CW) and anticlockwise (ACW) topology / overall assembly handedness is defined as follows: with the N-terminal helix at 12 o'clock and coming out towards the viewer, the chain trace follows either a CW or ACW path.

### 1.2 Computational tools

#### CC+ search

Antiparallel coiled-coil trimers and bundles were analyzed through the online CC+ database.[1] Three-helix coiled coils (CCs) with an antiparallel orientation, hetero-oligomeric partnering, canonical sequence repeats, and consisting of many chains of more than 11 residues were searched with a sequence redundancy cut-off of ≤ 50%. Based on the database that was updated on 1st April 2023, 169 CCs were assigned from 285 α-helical sequences. 121 and 48 CCs were assigned anticlockwise and clockwise, respectively, upon visual inspection. Amino-acid profiles were compiled for each position of the heptad repeat, *abcdefg*. The profile was normalized using the expected amino-acid frequencies from SWISS-PROT to give propensities for each residue.

#### AlphaFold2 and ESMFold predictions

AlphaFold2-multimer predictions (AF2) were performed using the ColabFold (version 1.5.2) with seed 000, single sequence mode on, and three recycle steps to return the top 5 rank models.[2, 3] Antiparallel CC trimeric peptides were tested in order to assess the predicted models for each possible assembly. The same single sequence mode was used to assess the sc-apCC3 models. Predicted local distance difference test (pLDDT) scores per residue, predicted template modelling score (pTM), and the predicted alignment error (pAE) matrices were collected to evaluate the quality and confidence of each prediction. To further ensure consistency in topology, particularly important when designing clockwise (CW) and anticlockwise (ACW) variants of largely similar related sequences, ESMFold was used as another structure prediction method. This served to verify that all models retained the expected fold and did not diverge significantly from AF2 predictions in overall topology.

#### MASTER

Method of Accelerated Search for Tertiary Ensemble Representatives (MASTER)[4, 5] was used to build fragments (loops) between adjacent helices in the antiparallel trimeric peptides to connect the N-to-C terminus into a single chain protein (Table S5).

The built-in MASTER database was used and the following parameters were set to find matches to the antiparallel queries (the first and last heptad of each chain): rmsdcut=0.5 and wgap=‘3-10’.

ProteinMPNN

After MASTER, ProteinMPNN[6] (v_48_020; T = 0.2, Cys, Met, Arg, Asn and Gly omitted, 10 sequences, training noise = 0.2 Å, seed 42) was used to optimize the sequences of the loop region. The peptide backbones were fixed throughout the design process. The 10 designs were then run through AF2 and ESMFold. The sequences with the highest pLDDT (> 85) from AF2, and ESMFold models in agreement with the intended topology were selected for experimental characterizations upon visual inspection.

Dictionary of Secondary Structure of Proteins DSSP

Secondary structure standardization assignments were generated using the DSSP[7, 8] algorithm based on AF2 predicted models. DSSP was used to annotate secondary structure elements and assess the extent of predicted helicity across designed sequences. These assignments were used to verify compatibility with the intended three-helix bundle topology.

### 1.3 Peptide synthesis and purification

#### Automated microwave Fmoc solid-phase peptide synthesis (SPPS)

Automated microwave SPPS was performed on a Liberty Blue (CEM) synthesizer with inline UV monitoring. The synthesis was performed on a 0.1 mmol scale with side-chain protection of the amino acids as follows: Gln(Trt), Glu(O*t*Bu), Lys(Boc), Tyr(*t*Bu), Trp(Boc). Rink amide (MBHA, 0.65 mmol/g loading, 100-200 mesh) was used for a synthesis on a 0.1 mmol scale. The coupling reactions were performed by adding amino acids dissolved in DMF (2.5 mL, 0.2 M), the coupling reagent DIC in DMF (1.0 mL, 1 M) and Oxyma Pure in DMF (1 mL, 0.5 M) to the respective resin. Standard couplings were performed at 90 °C for 4.5 min (100 W for 20 s, 60 W for 10 s, 35 W for 240 s). Standard deprotections were performed using 20% (v/v) morpholine in DMF at 90 °C for 1.5 min (125 W 30 s, 32 W 60 s). All peptides were manually acetyl capped through addition of pyridine (0.5 mL) and acetic anhydride (0.25 mL) in DMF (9.25 mL), shaking at room temperature (rt) for 20 minutes. The resin was washed 3 times with DMF followed by 6 times with DCM before cleavage. Peptides were cleaved from the resin with addition of 10 mL of a mixture 95:2.5:2.5 *v/v* TFA / $H_2O$ / TIPS, shaking at rt for 2 hours. The TFA solution was then filtered to remove the resin beads and was reduced in volume to ≈ 5 mL or lower using a flow of $N_2$. Cleaved peptide was precipitated with cold diethyl ether (≈ 45 mL), isolated *via* centrifugation and dissolved in a 1:1 mixture MeCN/$H_2O$. Crude peptides were lyophilized to yield a white or off-white powder. Crude peptides were kept in -20 °C for short-term storage, and -80 °C for long-term storage.

#### Semi-preparative scale high-performance liquid chromatography (HPLC)

All peptides were purified by reverse phase HPLC (JASCO) using a Luna C18 (Phenomenex) column (150 x 10 mm$^2$, 5 μM particle size, 100 Å pore size). Crude peptide was dissolved at 7 mg/mL in 20% *v/v* MeCN in $H_2O$ with 0.1% TFA, injected onto the column and eluted with a 3 mL/min linear gradient (20-100%) of MeCN in $H_2O$ with 0.1% TFA each over 40 min. Elution of the peptide was detected with inline UV monitoring at 220 and 280 nm wavelengths simultaneously. A column oven (50 °C) was employed to improve separation. Pure fractions were identified by analytical

HPLC and matrix-assisted laser desorption/ionization–time of flight (MALDI-TOF) mass spectrometry, then pooled, and lyophilized.

### Analytical HPLC

Analytical HPLC traces were obtained using a Jasco 2000 series HPLC system and a Phenomenex Kinetex C18 (100 x 4.6 $mm^2$, 5 µm particle size, 100 Å pore size) column. Chromatograms were monitored at 220 and 280 nm wavelengths. The linear gradient was 20–100% *v/v* MeCN in water (each containing 0.1% TFA) over 25 min at a flow rate of 1 mL/min. When required, a column oven (50 °C) was employed to assist peptide elution.

### Mass spectrometry

Mass spectra were collected on a Bruker UltraFlex MALDI-TOF mass spectrometer operating in positive-ion reflector mode. Peptides were spotted on a ground steel target plate using α-cyano-4-hydroxycinnamic acid dissolved in 1:1 *v/v* MeCN/$H_2O$ as the matrix. Masses quoted are for the monoisotopic mass as the singly protonated species.

### 1.4 Protein expression and purification

All genes were transformed then recombinantly expressed in *E. coli* Lemo21-DE3 (NEB) cells. Flasks containing 1 L of LB-kanamycin/chloramphenicol and 0.5 mM L-rhamnose were inoculated with 5 mL overnight cultures and incubated to an $OD_{600}$ of ~0.6 at 37 ˚C with 200 rpm shaking. Expression was induced with 0.5 mM IPTG, and cultures were incubated at 37 ˚C overnight with 200 rpm shaking.

Following expression, cultures were pelleted at 4 krpm for 10 minutes. Cell pellets were resuspended in 20 mL lysis buffer (50 mM Tris, pH 7.4, 500 mM NaCl, 30 mM imidazole, 1 mg/mL lysozyme) for 30 min at 37 ˚C. Resuspended pellets were sonicated using a Biologics Model 3000 Ultrasonic homogenizer with settings to 50% power, 90% pulser (1 pulse/second) for 7 minutes, then clarified at 13 krpm for 30 min. The clarified lysate was heat shocked at 65 ˚C for 10 minutes then cooled on ice for 10 minutes before re-clarifying at 13 krpm for 10 minutes.

The expressed proteins were first purified with Ni-affinity chromatography. Filtered lysate was loaded onto an ÄKTAprime plus (GE Healthcare) equipped with a HisTrap-5 mL HP column (Cytiva). His-tagged proteins were eluted at 55% Buffer B (Buffer A: 50 mM Tris, 500 mM NaCl, 30 mM imidazole, pH 7.4; Buffer B: 50 mM Tris, 500 mM NaCl, 300 mM imidazole, pH 7.4) using a step gradient. Fractions were combined and further purified by size exclusion chromatography using a HiLoad 16/600 Superdex 75 pg size exclusion column (GE Healthcare) equilibrated in buffer containing 50 mM sodium phosphate, pH 7.4, 150 mM NaCl. Eluted fractions were pooled, concentrated, and run on SDS-PAGE to confirm identity. Pure fractions were buffer exchanged into crystallization buffer(20 mM Tris, 50 mM NaCl, pH 8.0).

### 1.5 Solution-phase biophysical characterization

#### Peptide and protein concentration determination

Peptide and protein concentration was determined at 280 nm using a Nanodrop 2000 (ThermoScientific) spectrometer ($\varepsilon_{280}$(Trp) = 5690 cm$^{-1}$; $\varepsilon_{275}$(Tyr) = 1405 cm$^{-1}$, Table S4).

#### Circular dichroism (CD) spectroscopy

CD data were collected on a JASCO J-810 or J-815 spectropolarimeter fitted with a Peltier temperature controller in the far-UV region. Samples were made up as 100 μM overall assembly or individual peptide solution in phosphate buffered saline (PBS; 8.2 mM sodium phosphate dibasic, 1.8 mM potassium phosphate monobasic, 137 mM sodium chloride, 2.4 mM potassium chloride), pH 7.4 at 5 °C. For the sc-apCC-3 proteins, CD spectra were acquired at 10 μM protein concentration in PBS at 5 °C. Data were collected in a 1 mm quartz cuvette between 190 and 260 nm and the instrument was set as follows: band width 1 nm, data pitch 1 nm, scanning speed 100 nm/min, 1 s response time. Every CD curve was obtained by averaging 8 scans and subtracting the background signal of buffer and cuvette.

For thermal denaturation experiments, peptide unfolding and refolding were monitored at a wavelength of 222 nm. A temperature range of 5–95 °C and a temperature ramp rate of 60 °C per hour were used with the same settings and peptide or protein concentration as above. Chemical denaturation experiments for sc-apCC3-CW-3NTT were performed across a series of fixed guanidinium chloride (GdmHCl) concentrations prepared in PBS. The sc-apCC3-CW-3NTT samples were subjected to temperature ramping in 2 °C steps (heating rate of 2 °C/min) and the CD signal at 222 nm was monitored. The resulting unfolding curves were analyzed by fitting to a two-state model (folded ↔ unfolded), allowing the signal to vary linearly with denaturant concentration ([GdmHCl]).

The spectra were converted from ellipticities (mdeg) to mean residue ellipticities (MRE, deg·cm$^2$·dmol$^{-1}$·res$^{-1}$) by normalizing for concentration of peptide bonds and the cell path length using the equation:

$$MRE = \frac{\theta \times 10^6}{c \times l \times n}$$

where the variable $\theta$ is the measured difference in absorbed circularly polarized light in millidegrees, $c$ is the micromolar concentration of the compound, $l$ is the path length of the cuvette in mm, and $n$ is the number of amide bonds in the polypeptide.

Analytical ultracentrifugation (AUC)

Analytical ultracentrifugation (AUC) was performed on a Beckman Optima X-LI analytical ultracentrifuge with an An-60-Ti rotor (Beckman-Coulter). Buffer densities, viscosities and peptide partial specific volumes ($\bar{v}$) were calculated using SEDNTERP (http://rasmb.org/sednterp/).

For sedimentation velocity (SV), peptide sample solutions of 310 or 410 μL were prepared in PBS at sample concentration (250 μM for assembly or individual peptide, 50 μM for protein) and placed in a sedimentation velocity cell with an epon or aluminium, 2-channel centerpiece and sapphire windows. The reference channel was loaded with 320 μL (or 420 μL, respectively) of PBS buffer. The samples were centrifuged at 60 krpm at 20 °C, with absorbance scans taken across a radial range of 5.8–7.3 cm at 5 min intervals to a total of 120 scans.

Data from a single run were fitted to a continuous c(*s*) distribution model using SEDFIT, at 95% confidence level. Residuals for sedimentation velocity experiments are shown as a bitmap in which the grayscale shade indicates the difference between the fit and raw data (residuals < -0.05 black, > 0.05 white). Good fits are uniformly grey without major dark or light streaks.

Sedimentation equilibrium (SE) experiments were performed in triplicate in a six-channel centerpiece (110 μL per channel) channels at 20 °C. The samples were centrifuged at speeds in the range of 44-60 krpm and scans at each recorded speed were duplicated after equilibration for 8 hours. Data were fitted using SEDPHAT to a single species model. Monte Carlo analysis was performed to give 95% confidence limits.

### 1.6 Structural characterization

#### Crystal growth

Diffraction-quality peptide and protein crystals were grown using a sitting-drop vapor-diffusion method. Mixtures of acidic, basic and neutral peptides and sc-apCC3 proteins were annealed beforehand. Purified protein in 20 mM Tris pH 8.0, 50 mM NaCl was concentrated to approximately 20 mg/mL. Protein samples were annealed to 90 °C in a heat block (Grant), incubated at 90 °C for 10 min and slowly cooled down to room temperature over 90 min.

Commercially available sparse matrix screens were used (Morpheus®, JCSG-plus™, Structure Screen 1 and 2, Pact Premier™, ProPlex™; Molecular Dimensions), and the drops were dispensed using a robot (Oryx8; Douglas Instruments). For each well of an MRC 2-drop plate, 0.3 µL of peptide or protein solution and 0.3 µL of reservoir solution in parallel with 0.4 µL of the peptide or protein solution and 0.2 µL of reservoir solution were mixed and the plate was incubated at 20 °C. Crystals generally formed within a month, and after looping were soaked in reservoir solution containing 25% glycerol as a cryoprotectant.

Crystals of sc-apCC3 and its variants were obtained by optimization by seeding. For seeding experiments, 16-20 mg/mL of protein solution were used and for each well of an MRC 2 drop plate, 0.3 µL protein solution, 0.1 µL of seed, and 0.2 µL of reservoir solution in parallel with 0.4 µL of the peptide or protein solution, 0.1 µL of seed, and 0.1 µL of reservoir solution were mixed and the plate was incubated at 20 °C.

Final crystallization conditions for proteins are provided in Table S7.

#### X-ray crystal structure determination

Diffraction data for the crystals were obtained at the Diamond Light Source on beamlines I03, I04, and I24.

Data were processed using the automated pipelines: Xia2 pipelines, which ports data through DIALS or MOSFLM to POINTLESS and AIMLESS as implemented in the CCP4 suite[9], or XDS to XSCALE. Final structures were obtained after iterative rounds of model building with COOT and refinement with PHENIX Refine[10]. Solvent-exposed atoms lacking map density were either deleted or left at full occupancy. Data collection and refinement statistics are provided in Table S8.

**Supplementary data**

1.7 Supplementary tables

Table S1. Raw amino-acid counts for antiparallel three-helix coiled coils found by the CC+ database[1]

| - | a | b | c | d | e | f | g | Sum |
|---|---|---|---|---|---|---|---|---|
| A | 161 | 110 | 128 | 295 | 84 | 156 | 93 | 1027 |
| C | 31 | 7 | 10 | 17 | 3 | 16 | 15 | 99 |
| D | 8 | 86 | 71 | 10 | 48 | 86 | 60 | 369 |
| E | 27 | 179 | 204 | 32 | 159 | 192 | 137 | 930 |
| F | 72 | 22 | 22 | 49 | 30 | 26 | 38 | 259 |
| G | 25 | 34 | 34 | 58 | 25 | 54 | 20 | 250 |
| H | 29 | 36 | 31 | 15 | 32 | 28 | 36 | 207 |
| I | 205 | 46 | 55 | 196 | 127 | 49 | 95 | 773 |
| K | 26 | 138 | 136 | 22 | 120 | 143 | 128 | 713 |
| L | 431 | 94 | 91 | 382 | 222 | 104 | 241 | 1565 |
| M | 34 | 24 | 16 | 31 | 43 | 15 | 35 | 198 |
| N | 39 | 66 | 55 | 46 | 47 | 64 | 57 | 374 |
| P | 2 | 7 | 7 | 10 | 6 | 12 | 1 | 45 |
| Q | 56 | 109 | 104 | 63 | 144 | 106 | 128 | 710 |
| R | 18 | 93 | 85 | 23 | 129 | 98 | 92 | 538 |
| S | 64 | 67 | 75 | 79 | 56 | 96 | 61 | 498 |
| T | 81 | 43 | 57 | 66 | 43 | 76 | 71 | 437 |
| V | 166 | 71 | 54 | 118 | 86 | 51 | 59 | 605 |
| W | 15 | 15 | 10 | 8 | 9 | 10 | 10 | 77 |
| Y | 53 | 27 | 29 | 39 | 41 | 21 | 61 | 271 |
| Sum | 1543 | 1274 | 1274 | 1559 | 1454 | 1403 | 1438 | 9945 |

Table S2. Helical regions identified for antiparallel three-helix coiled coils found by the CC+ database

| PDB ID | Helix Range (Start-End:Chain) | Sequence (Registry) |
|---|---|---|
| 1bg1 | 151-179:A 199-228:A 270-284:A | VRKRVQDLEQKMKVVENLQDDFDFNYKTL(defgabcdefgabcdefgabcdefgabcd) KMQQLEQMLTALDQMRRSIVSELAGLLSAM(gabcdefgabcdefgabcdefgabcdefga) LAESQLQTRQQIKKL(abcdefgabcdefga) |
| 1e12_ba1 | 139-150:A 62-75:C 111-125:C | AADIGMCVTGLA(defgabcdefga) IWGATLMIPLVSIS(efgabcdefgabcd) TWALSTPMILLALGL(abcdefgabcdefga) |
| 1env_ba1 | 6-76:A 124-149:A 2-72:C | IEEILSKIYHIENEIARIKKLIGEARQLLSGIVQQQNNLLRAIEAQQHLLQLTVWGIKQLQARILAVERYL(defgabcdefgabcdefgabcdefgabcdefgabcdefgabcdefgabcdefgabcdefgabcd) INNYTSLIHSLIEESQNQQEKNEQEL(defgabcdefgabcdefgabcdefga) IEDKIEEILSKIYHIENEIARIKKLIGEARQLLSGIVQQQNNLLRAIEAQQHLLQLTVWGIKQLQARILAV(abcdefgabcdefgabcdefgabcdefgabcdefgabcdefgabcdefgabcdefgabcdefga) |
| 1ez3 | 37-61:A 71-103:A 113-130:A | VEEIRGFIDKIAENVEEVKRKHSAI(abcdefgabcdefgabcdefgabcd) TKEELEELMSDIKKTANKVRSKLKSIEQSIEQE(gabcdefgabcdefgabcdefgabcdefgabcd) LRIRKTQHSTLSRKFVEV(abcdefgabcdefgabcd) |
| 1fio | 40-64:A 76-101:A 115-129:A | ISQINRDLDKYDHTINQVDSLHKRL(abcdefgabcdefgabcdefgabcd) LRHSLDNFVAQATDLQFKLKNEIKSA(gabcdefgabcdefgabcdefgabcd) AENSRQRFLKLIQDY(abcdefgabcdefga) |
| 1fs7 | 355-369:A 379-393:A 417-431:A | AFDNIGKAHLETGKA(defgabcdefgabcd) LKEIRTHIRHAQWRA(abcdefgabcdefga) GNEEAQKARIKLVKV(defgabcdefgabcd) |
| 1gu6 | 340-357:A 367-384:A 402-419:A | KIKVEDQLVHAHFEAKAA(abcdefgabcdefgabcd) MKPIQDDIRHAQWRWDLA(abcdefgabcdefgabcd) LGTAMDKAADARTKLARL(abcdefgabcdefgabcd) |
| 1gu6 | 336-357:E 367-384:E 399-419:E | INDLKIKVEDQLVHAHFEAKAA(defgabcdefgabcdefgabcd) MKPIQDDIRHAQWRWDLA(abcdefgabcdefgabcd) LRMLGTAMDKAADARTKLARL(efgabcdefgabcdefgabcd) |
| 1hx1 | 159-179:B 207-218:B 233-254:B | LKKLKHLEKSVEKIADQLEEL(efgabcdefgabcdefgabcd) VKATIEQFMKIL(abcdefgabcde) SRLKRKGLVKKVQAFLAECDTV(defgabcdefgabcdefgabcd) |

| | | |
|---|---|---|
| 1hz4 | 51-62:A 70-81:A 89-100:A | IVATSVLGEVLH(defgabcdefga)<br>SLALMQQTEQMA(defgabcdefga)<br>YALWSLIQQSEI(defgabcdefga) |
| 1o5h | 25-40:B 66-80:B 108-119:B | GAVGSVVGAXACALAE(defgabcdefgabcde)<br>XEEARLKLFDLAKKD(abcdefgabcdefga)<br>VPXDVIRVXKDL(defgabcdefga) |
| 1ocr | 19-30:C 42-56:C 31-42:J | TGALSALLMTSG(defgabcdefga)<br>LLMIGLTTNMLTMYQ(defgabcdefgabcd)<br>LYRVTMTLCLGG(gabcdefgabcd) |
| 1qbz | 33-75:A 119-144:A 32-71:C | LAGIVQQQQQLLDVVKRQQELLRLTVWGTKNLQTRVTAIEKYL(defgabcdefgabcdefgabcdefgabcdefgabcdefgabcd)<br>VDFLEENITALLEEAQIQQEKNMYEL(defgabcdefgabcdefgabcdefga)<br>LLAGIVQQQQQLLDVVKRQQELLRLTVWGTKNLQTRVTAI(defgabcdefgabcdefgabcdefgabcdefgabcdefga) |
| 1r8i | 41-59:A 150-167:A 176-190:A | LEQMAQQLEQLKSQLETQK(abcdefgabcdefgabcde)<br>YNNQMQELSDMQALTEQI(abcdefgabcdefgabcd)<br>IADLQARIQTSQGAI(abcdefgabcdefga) |
| 1u7l | 61-86:A 127-155:A 281-317:A | LIVESEELSKVDNQIGASIGKIIEIL(abcdefgabcdefgabcdefgabcde)<br>IKDLITLISNESSQLDADVRATYANYNSA(defgabcdefgabcdefgabcdefgabcd)<br>EQSLRVQLVRLAKTAYVDVFINWFHIKALRVYVESVL(defgabcdefgabcdefgabcdefgabcdefgabcde) |
| 1uur | 251-265:A 288-313:A 326-351:A | IYKLLSEQEQTLVQM(defgabcdefgabcd)<br>LKSLSQKQITLSGQMNTEMSALDATK(abcdefgabcdefgabcdefgabcde)<br>LFALKQDLQIQFKQLSLLHNEIQSIL(abcdefgabcdefgabcdefgabcde) |
| 1wp7 | 147-171:A 459-481:A 144-165:C | LKSSIESTNEAVVKLQETAEKTVYV(abcdefgabcdefgabcdefgabcd)<br>QISSMNQSLQQSKDYIKEAQRLL(gabcdefgabcdefgabcdefga)<br>INKLKSSIESTNEAVVKLQETA(defgabcdefgabcdefgabcd) |
| 1y1u | 148-172:A 218-243:A 282-296:A | FEELRLITQDTENELKKLQQTQEYF(abcdefgabcdefgabcdefgabcd)<br>EAQTLQQYRVELAEKHQKTLQLLRKQ(gabcdefgabcdefgabcdefgabcd)<br>LAEIIWQNRQQIRRA(abcdefgabcdefga) |
| 1ztm | 147-171:A 151-172:B 460-481:C | LKEAIRDTNKAVQSVQSSIGNLIVA(abcdefgabcdefgabcdefgabcd)<br>IRDTNKAVQSVQSSIGNLIVAI(defgabcdefgabcdefgabcd)<br>LNKAKSDLEESKEWIRRSNQKL(abcdefgabcdefgabcdefga) |

| | | |
|---|---|---|
| 2bug | 62-73:A 78-91:A 98-109:A | AIYYGNRSLAYL(defgabcdefga) YGYALNDATRAIEL(abcdefgabcdefg) GYYRRAASNMAL(abcdefgabcde) |
| 2c0m | 519-530:F 535-548:F 566-577:F | IRSRYNLGISCI(defgabcdefga) HREAVEHFLEALNM(abcdefgabcdefg) IWSTLRLALSML(abcdefgabcde) |
| 2cwo | 13-24:A 39-50:A 78-89:A | ALSRSESLLRRV(defgabcdefga) CVDEFNELASFN(defgabcdefga) IGEMLKEIRAFL(defgabcdefga) |
| 2d4c | 29-43:D 151-165:D 197-211:D | LDDDFKEMERKVDVT(defgabcdefgabcd) LREIQSALQHHLKKL(abcdefgabcdefga) SKEIAESSMFNLLEM(defgabcdefgabcd) |
| 2e2a | 10-30:C 39-50:C 76-93:C | GFEIVAYAGDARSKLLEALKA(abcdefgabcdefgabcdefg) ADSLVVEAGSCI(defgabcdefga) MMHGQLHLMTTILLKDVI(abcdefgabcdefgabcd) |
| 2ein | 19-34:C 42-56:C 31-42:J | TGALSALLMTSGLTMW(defgabcdefgabcde) LLMIGLTTNMLTMYQ(defgabcdefgabcd) LYRVTMTLCLGG(gabcdefgabcd) |
| 2fyz | 130-172:A 126-168:C 452-474:D | AQTNARAIAAMKNSIQATNRAVFEVKEGTQRLAIAVQAIQDHI (defgabcdefgabcdefgabcdefgabcdefgabcdefgabcd) SLVQAQTNARAIAAMKNSIQATNRAVFEVKEGTQRLAIAVQAI (abcdefgabcdefgabcdefgabcdefgabcdefgabcdefga) ELSKVNASLQNTVKYIKESNHQL(gabcdefgabcdefgabcdefga) |
| 2gnx | 193-204:A 212-223:A 258-269:A | VLSHLLKAQAQI(defgabcdefga) SLVTLHNAHTKL(defgabcdefga) LXKLKTXLLAKF(defgabcdefga) |
| 2hr2 | 10-21:C 29-40:C 54-65:C | AYLALSDAQRQL(defgabcdefga) AAANCRRAXEIS(defgabcdefga) FDAFCHAGLAEA(defgabcdefga) |
| 2ieq | 7-45:A 77-102:A 6-48:C | VASFSSVNDAITQTAEAIHTVTIALNKIQDVVNQQGSAL(abcdefgabcdefgabcdefgabcdefgabcdefgabcd) ELKQLEAKTASLFQTTVELQGLIDQI(gabcdefgabcdefgabcdefgabcd) IVASFSSVNDAITQTAEAIHTVTIALNKIQDVVNQQGSALNHL (abcdefgabcdefgabcdefgabcdefgabcdefgabcdefga) |
| 2iub | 175-198:A 207-236:A 251-275:A | DALVDDYFVLLEKIDDEIDVLEEE(efgabcdefgabcdefgabcdefg) TVQRTHQLKRNLVELRKTIWPLREVLSSLY(defgabcdefgabcdefgabcdefgabcde) FRDVYDHTIQIADTVETFRDIVSGL(abcdefgabcdefgabcdefgabcd) |

| | | |
|---|---|---|
| 2j7a | 372-386:K 407-422:K 440-460:K | TFDLLLAAQEVSVKA(defgabcdefgabcd) LMIQAREMVRKGQFFW(gabcdefgabcdefga) LDTLAQSQQFSQKAIDLAMEA(efgabcdefgabcdefgabcd) |
| 2n8i | 28-39:A 44-58:A 67-81:A | GRALYNIGLEKN(defgabcdefga) VKEAIEYFLRAKKVF(abcdefgabcdefga) ARRAAKSLSEAYQKV(efgabcdefgabcde) |
| 2ntx | 113-124:A 179-200:A 330-352:A | NIPALRKLDAXL(defgabcdefga) LYFQKDSVTQVQKAAXAINAQV(abcdefgabcdefgabcdefga) LESYSRILESLAYTVXSRIEDVL(defgabcdefgabcdefgabcde) |
| 2odu | 424-449:A 462-486:A 501-519:A | IVTKLQMEAGLCEEQLNQADALLQSD(gabcdefgabcdefgabcdefgabcd) AGEVERDLDKADSMIRLLFNDVQTL(abcdefgabcdefgabcdefgabcd) VYRLHKRLVAIRTEYNLRL(abcdefgabcdefgabcde) |
| 2oex | 430-457:A 492-516:A 663-691:A | IQTVDQLIKELPELLQRNREILDESLRL(abcdefgabcdefgabcdefgabcdefg) GTNFRTVLDKAVQADGQVKECYQSH(abcdefgabcdefgabcdefgabcd) FVELVANLKEGTKFYNELTEILVRFQNKC(abcdefgabcdefgabcdefgabcdefga) |
| 2p22 | 254-286:A 89-126:C 43-61:D | MQESIARFHEIIAIDKNHLRAVEQAIEQTMHSL(defgabcdefgabcdefgabcdefgabcdefga) AGKIHAFRDQFKQLEENFEDLHEQKDKVQALLENARIL(efgabcdefgabcdefgabcdefgabcdefgabcdefg) TRDLLCPWYEECDNITKVC(defgabcdefgabcdefga) |
| 2qfc | 155-166:A 174-185:A 194-205:A | LYIENAIANIYA(defgabcdefga) GIDLFEQILKQL(defgabcdefga) FDVKVRYNHAKA(defgabcdefga) |
| 3a8y | 372-383:D 398-409:D 427-438:D | VLGNLSEIQGEV(defgabcdefga) LEELLTKQLLAL(defgabcdefga) AVRLAQNILSYL(defgabcdefga) |
| 3atp | 56-70:A 90-104:A 129-140:A | SWVALLQTRNTLNRA(defgabcdefgabcd) LMESASISLKQAEKN(gabcdefgabcdefg) YDIYHNALAELI(defgabcdefga) |
| 3c18 | 122-140:B 148-165:B 204-215:B | XSLSFAKLLRRFQDGRNLF(defgabcdefgabcdefga) AYTHVHHALHHLARLSVL(defgabcdefgabcdefg) IHLALIGLEHLL(defgabcdefga) |
| 3efz | 142-156:B 167-182:B 194-205:B | AFCIKLKGDLMRYKA(defgabcdefgabcd) CIKQAVEFYEDALQRE(gabcdefgabcdefga) LYLATILNYTIL(defgabcdefga) |
| 3etv | 73-84:A 92-103:A 120-131:A | FLNLIKEVKTNL(defgabcdefga) CYYSLQSLRKKM(defgabcdefga) ISTYVDTLHLEL(defgabcdefga) |

| | | |
|---|---|---|
| 3f4y | 4-35:A 3-31:B 10-31:E | VQQQNNLLRAIEAQQHLLQLTVWGIKQLQARI(abcdefgabcdefgabcdefgabcdefgabcd) IVQQQNNLLRAIEAQQHLLQLTVWGIKQL(abcdefgabcdefgabcdefgabcdefga) IAEYAARIEALIRAAQEQQEKN(defgabcdefgabcdefgabcd) |
| 3fp2 | 161-172:A 177-190:A 197-208:A | PVFYSNISACYI(defgabcdefga) LEKVIEFTTKALEI(abcdefgabcdefg) ALLRRASANESL(abcdefgabcde) |
| 3h90 | 80-94:A 145-156:A 190-205:A | SLAALAQSMFISGSA(abcdefgabcdefga) QAVRADMLHYQS(defgabcdefga) ILYSALRMGYEAVQSL(defgabcdefgabcde) |
| 3k1s | 8-29:H 38-59:H 74-85:H | IPFQLILNSGNARSFAXEALQF(gabcdefgabcdefgabcdefg) ADEAXVKAKEAINEAHHFQTEL(defgabcdefgabcdefgabcd) LLIHAQDHLXNA(gabcdefgabcd) |
| 3l8r | 17-31:C 39-53:C 83-97:C | SGNARSIVHEAFDAM(abcdefgabcdefga) AEQKLQEANDELLKA(defgabcdefgabcd) LMTTMTLREVAIEML(abcdefgabcdefga) |
| 3m71_ba1 | 15-27:B 41-52:B 85-96:B | GIPLGLAALSLAW(bcdefgabcdefg) SDVLGIVASAVW(defgabcdefga) LIPITTMLVGDI(gabcdefgabcd) |
| 3mx3 | 113-124:A 160-171:A 190-201:A | GLAHFIKAYKIF(defgabcdefga) ILSDLETAYELF(defgabcdefga) WKKATTAFQLKV(defgabcdefga) |
| 3n27 | 8-29:A 58-76:A 4-31:C | INKLKSSIESTNEAVVKLQETA(defgabcdefgabcdefgabcd) AKSDLEESKEWIRRSNQKL(defgabcdefgabcdefga) NADNINKLKSSIESTNEAVVKLQETAEK(abcdefgabcdefgabcdefgabcdefg) |
| 3nqp | 142-156:A 191-205:A 224-237:A | GEVYGLRAFYYFDLY(defgabcdefgabcd) XTQIKSDLNKSXEYF(abcdefgabcdefga) KAATECLXGEVYLW(efgabcdefgabcd) |
| 3okq_ba1 | 562-587:A 603-670:A 603-673:B | SQTELGDLSDTLLSKVDDLQDVIEIM(gabcdefgabcdefgabcdefgabcd) LETVSKDLENAQADVLKLQEFIDTEKPHWKKTWEAELDKVCEEQQFLTLQEELILDLKEDLGKALETF(abcdefgabcdefgabcdefgabcdefgabcdefgabcdefgabcdefgabcdefgabcdefgabcde) LETVSKDLENAQADVLKLQEFIDTEKPHWKKTWEAELDKVCEEQQFLTLQEELILDLKEDLGKALETFDLI(defgabcdefgabcdefgabcdefgabcdefgabcdefgabcdefgabcdefgabcdefgabcdefgabcd) |
| 3q15 | 139-150:A 158-169:A 178-189:A | AEFHFKVAEAYY(defgabcdefga) SMYHILQALDIY(defgabcdefga) RTIQSLFVIAGN(defgabcdefga) |

| | | |
|---|---|---|
| 3sxq | 434-455:A 468-479:A 497-518:A | THGKIMKAEFWLARMIDLFPVA(defgabcdefgabcdefgabcd) VRALHYDAHLHW(defgabcdefga) RESLMKSITKSKEGVGKLDAAI(efgabcdefgabcdefgabcde) |
| 3txn | 208-219:A 228-239:A 246-257:A | GALDLQSGILHA(defgabcdefga) AFSYFYEAFEGF(defgabcdefga) KALTSLKYMLLC(defgabcdefga) |
| 3ulq | 141-152:A 160-171:A 180-191:A | AEFFFKMSESYY(defgabcdefga) SMDYARQAYEIY(defgabcdefga) RLLQCHSLFATN(defgabcdefga) |
| 3um3 | 240-261:A 270-291:A 309-323:A | KYLHLKMCFYTAYAYCYHGETL(gabcdefgabcdefgabcdefg) AIRSLQEAEKLYAKAEALCKEY(defgabcdefgabcdefgabcd) FRKLGNLVKNTLEKC(abcdefgabcdefga) |
| 3uun | 352-365:A 372-389:A 427-452:A | LSWLLSAEDTLQAQ(efgabcdefgabcd) VEVVKDQFHTHEGYMMDL(efgabcdefgabcdefga) LNSRWECLRVASMEKQSNLHRVLMDL(defgabcdefgabcdefgabcdefga) |
| 3vtq | 38-69:E 37-65:B 128-145:D | VQQQNNLLRAIEAQQHLLQLTVWGIKQLQARI(abcdefgabcdefgabcdefgabcdefgabcd) IVQQQNNLLRAIEAQQHLLQLTVWGIKQL(abcdefgabcdefgabcdefgabcdefga) TKQIYKILEESQEQQDRN(abcdefgabcdefgabcd) |
| 3wmi_ba1 | 487-526:A 572-599:B 502-523:E | VQNHTFEVENNTINGLELVEEQVHILYAMVLQTHADVQLL(defgabcdefgabcdefgabcdefgabcdefgabcdefga) VDKMENLNHDILTTLHTARNNLEQSMIT(abcdefgabcdefgabcdefgabcdefg) LELVEEQVHILYAMVLQTHADV(defgabcdefgabcdefgabcd) |
| 3zc1 | 23-38:D 46-60:D 77-88:D | VLREMRIHSTKSIALI(gabcdefgabcdefga) AEQELKKAIELLEKV(defgabcdefgabcd) AMQELVEAIAFK(defgabcdefga) |
| 4a1s | 157-178:B 190-211:B 217-230:B | RLSEGRALYNLGNVYHAKGKHL(gabcdefgabcdefgabcdefg) VKEALTRAVEFYQENLKLMRDL(defgabcdefgabcdefgabcd) QGRACGNLGNTYYL(efgabcdefgabcd) |
| 4e68 | 151-179:A 199-228:A 270-287:A | VRKRVQDLEQKMKVVENLQDDFDFNYKTL(defgabcdefgabcdefgabcdefgabcd) KMQQLEQMLTALDQMRRSIVSELAGLLSAM(gabcdefgabcdefgabcdefgabcdefga) LAESQLQTRQQIKKLEEL(abcdefgabcdefgabcd) |
| 4egw | 144-161:A 177-198:A 221-241:A | TRSYSRILMNLEDELEEL(abcdefgabcdefgabcd) ILGLRKTLVYFHKSLIANRDVL(abcdefgabcdefgabcdefga) YYDTLQLIDMSATYREVLTSM(efgabcdefgabcdefgabcd) |

| | | |
|---|---|---|
| 4eqf | 201-212:A<br>217-230:A<br>237-248:A | YSLWNRLGATLA(defgabcdefga)<br>SEEAVEAYTRALEI(abcdefgabcdefg)<br>SRYNLGISCINL(abcdefgabcde) |
| 4ev6 | 144-161:A<br>174-202:A<br>217-245:A | TRSYSRILMNLEDELEEL(abcdefgabcdefgabcd)<br>MEKILGLRKTLVYFHKSLIANRDVLVLLK(efgabcdefgabc defgabcdefgabcde)<br>FEDLYYDTLQLIDMSATYREVLTSMMDIT(abcdefgabcdef gabcdefgabcdefga) |
| 4f53 | 134-145:A<br>179-190:A<br>213-224:A | SLAELYRIVVXL(defgabcdefga)<br>XLEDLDNIITVL(defgabcdefga)<br>WYKFANSLKLRX(defgabcdefga) |
| 4gpk | 189-200:A<br>208-219:A<br>227-238:A | TGLYYNIALAYT(defgabcdefga)<br>AIHFVNMALEGF(defgabcdefga)<br>NIINCQILIAVS(defgabcdefga) |
| 4gpk | 229-240:I<br>245-259:I<br>268-283:I | INCQILIAVSYT(defgabcdefga)<br>YEEALKMYESILREA(abcdefgabcdefga)<br>LLAITLSNMGSIYYKK(defgabcdefgabcde) |
| 4hwc | 164-175:C<br>196-207:C<br>223-234:C | ISLEVDRLGGRV(defgabcdefga)<br>VIELLMNELIKL(defgabcdefga)<br>QVKRVQNYVETL(defgabcdefga) |
| 4hwd | 144-158:D<br>170-187:D<br>203-221:D | ISFQVERLAGQLSAF(defgabcdefgabcd)<br>EKNLENLMEMLMNQLVKL(efgabcdefgabcdefga)<br>QEERLHKYVEALDLLKIKN(defgabcdefgabcdefga) |
| 4hwh | 146-159:C<br>174-185:C<br>204-218:C | TGEVDKLSDRVVAL(efgabcdefgabcd)<br>FDMAAELLMRQL(abcdefgabcde)<br>EVRRIQNLQEAVDKL(defgabcdefgabcd) |
| 4i0u | 167-198:C<br>207-232:C<br>247-275:C | DYLLYSLIDALVDDYFVLLEKIDDEIDVLEEE(defgabcdef gabcdefgabcdefgabcdefg)<br>TVQRTHQLKRNLVELRKTIWPLREVL(defgabcdefgabcde fgabcdefga)<br>TVPYFRDVYDHTIQIADTVETFRDIVSGL(defgabcdefgab cdefgabcdefgabcd) |
| 4jhr | 131-149:B<br>171-188:B<br>194-207:B | KVGEARALYNLGNVYHAKG(gabcdefgabcdefgabcd)<br>LQAAVDLYEENLSLVTAL(abcdefgabcdefgabcd)<br>QGRAFGNLGNTHYL(efgabcdefgabcd) |
| 4kik | 451-469:A<br>533-544:A<br>612-629:A | QRAAMMNLLRNNSCLSKMK(defgabcdefgabcdefga)<br>LVERMMALQTDI(defgabcdefga)<br>LSKTVVCKQKALELLPKV(efgabcdefgabcdefga) |
| 4nqj | 168-257:A<br>273-287:A<br>160-257:C | QGQLETTLKELQTLRNMQKEAIAAHKENKLHLQQHVSMEFLKL HQFLHSKEKDILTELREEGKALNEEMELNLSQLQEQCLLAKDM LVSI(cdefgabcdefgabcdefgabcdefgabcdefgabcde fgabcdefgabcdefgabcdefgabcdefgabcdefgabcdef gabcdefga) ITTLLHSLEQGMKVL(defgabcdefgabcd)<br>FMEELAIQQGQLETTLKELQTLRNMQKEAIAAHKENKLHLQQH |

| | | |
|---|---|---|
| | | VSMEFLKLHQFLHSKEKDILTELREEGKALNEEMELNLSQLQEQCLLAKDMLVSI(defgabcdefgabcdefgabcdefgabcdefgabcdefgabcdefgabcdefgabcdefgabcdefgabcdefgabcdefgabcdefgabcdefgabc) |
| 4p1n | 660-678:A 711-728:A 472-483:C | KKLCMESLLLYLKSLTILA(defgabcdefgabcdefga) FNECLDKAEFLRLKLHTL(defgabcdefgabcdefg) ISSSLIKFQSMK(defgabcdefga) |
| 4r61 | 16-48:A 64-96:A 127-159:A | LLRQIEAQQHLLQLTVSKIKQLQARILAVERYL(defgabcdefgabcdefgabcdefgabcdefga) LYREVALIRAQLQKIESETLQLLHQQAEIEREL(defgabcdefgabcdefgabcdefgabcdefga) LKRAIEAQKHLLQLTVWGIKQLQARILAVERYL(defgabcdefgabcdefgabcdefgabcdefga) |
| 4u6u_ba3 | 46-71:A 111-136:B 111-139:H | CSSLLSKMDYYSGHITKELESTIQVL(gabcdefgabcdefgabcdefgabcd) FERALKLQTVSSKIHQTTTLLRSSLI(abcdefgabcdefgabcdefgabcde) FERALKLQTVSSKIHQTTTLLRSSLIYVH(defgabcdefgabcdefgabcdefgabcd) |
| 4ui9 | 703-714:F 719-732:F 739-750:F | PLCKFHRASVLF(defgabcdefga) YKSALQELEELKQI(abcdefgabcdefg) VYFLIGKVYKKL(abcdefgabcde) |
| 4ui9 | 581-592:O 600-611:O 619-630:O | ISVLLSVAELYW(defgabcdefga) ALPMLLQALALS(defgabcdefga) LASETVLNLAFA(defgabcdefga) |
| 4ui9 | 661-675:O 686-701:O 710-724:O | GRAMFLVAKCQVASA(defgabcdefgabcd) ALEAAIENLNEAKNYF(gabcdefgabcdefga) IRDVVYFQARLYHTL(efgabcdefgabcde) |
| 4wpc | 45-80:A 113-148:A 237-248:A | INALLSRLKQSLLTCEEFMKFIRKKYAFEEEHVQEL(defgabcdefgabcdefgabcdefgabcdefgabcd) AQVKQSYITALQKMYSEISSLLLTMTKLRKSVKENS(abcdefgabcdefgabcdefgabcdefgabcdefga) IVQELKDLILEI(defgabcdefga) |
| 4xng | 89-102:B 114-127:B 181-199:B | LVGLQQLELEYVNL(efgabcdefgabcd) TELLNNLKELVDEH(efgabcdefgabcd) LRQRFKKLSQKIDSSLKQI(defgabcdefgabcdefga) |
| 4ynw | 254-265:A 270-283:A 290-301:A | FPSRVSLVRLLI(defgabcdefga) EEEALEVTERLIAE(abcdefgabcdefg) VWYLGGYARYRL(abcdefgabcde) |
| 4yo5 | 436-447:C 455-466:C 478-489:C | WLLRLLXARVAE(defgabcdefga) ALHLLAELDERA(defgabcdefga) LVFEVKARRLKL(defgabcdefga) |
| 4yv6 | 185-196:D 204-215:D 223-234:D | TQLSINCLIISI(defgabcdefga) SHYLIEKIEFLL(defgabcdefga) EKTVFLYVHGYY(defgabcdefga) |

| | | |
|---|---|---|
| 5a7d | 298-309:C 314-328:C 337-351:C | AQSCYSLGNTYT (defgabcdefga) FNTAIEYHNRHLAIA (abcdefgabcdefga) EARACWSLGNAHSAI (efgabcdefgabcde) |
| 5aqf | 159-179:B 207-221:B 233-254:B | LKKLKHLEKSVEKIADQLEEL (efgabcdefgabcdefgabcd ) VKATIEQFMKILEEI (abcdefgabcdefga) SRLKRKGLVKKVQAFLAECDTV (defgabcdefgabcdefgab cd) |
| 5aqg | 159-176:B 207-221:B 233-255:B | LKKLKHLEKSVEKIADQL (efgabcdefgabcdefga) VKATIEQFMKILEEI (abcdefgabcdefga) SRLKRKGLVKKVQAFLAECDTVE (defgabcdefgabcdefga bcde) |
| 5cwi | 160-171:A 188-204:A 214-228:A | LASQAAEAVKLA (defgabcdefga) IKAASEAAEEASKAAEE (efgabcdefgabcdefg) ARDEIKEASQKAEEV (defgabcdefgabcd) |
| 5cwm | 3-24:A 33-54:A 62-83:A | PEDELKRVEKLVKEAEELLRQA (defgabcdefgabcdefgab cd) LEKALRTAEEAAREAKKVLEQA (abcdefgabcdefgabcdef ga) VALRAVELVVRVAELLLRIAKE (defgabcdefgabcdefgab cd) |
| 5cwo | 67-81:A 89-100:A 120-134:A | AAKVAAEVIKVAIQA (abcdefgabcdefga) LFRAALELVRAV (defgabcdefga) ARAAKVAAEVIKVAI (abcdefgabcdefga) |
| 5cwq | 33-51:A 70-84:A 92-109:A | LKELAEALIEEARAVQELA (defgabcdefgabcdefga) AQRVLEEARKVSEEA (abcdefgabcdefga) VLALALIAIALAVLALAE (defgabcdefgabcdefg) |
| 5ehb_ba1 | 4-21:A 8-25:B 8-26:C | IXDALEKLAEIQKEIAEF (abcdefgabcdefgabcd) LEKLAEIQKEIAEFLREL (efgabcdefgabcdefga) LEKLAEIQKEIAEFLRELI (defgabcdefgabcdefga) |
| 5gjq | 167-178:Q 186-197:Q 206-217:Q | VEVQLLESKTYH (defgabcdefga) ARAALTSARTTA (defgabcdefga) LQATLDMQSGII (defgabcdefga) |
| 5h0n | 18-39:A 61-75:B 14-42:E | QNNLLRAIEAQQHLLQLTVWGI (defgabcdefgabcdefgab cd) IEEYTKKIEELIKKS (defgabcdefgabcd) IVQQQNNLLRAIEAQQHLLQLTVWGIKQL (abcdefgabcdef gabcdefgabcdefga) |
| 5h6i | 50-61:A 80-91:A 104-115:A | MAQKLDQDSIQL (defgabcdefga) AITSVEKLKTSL (defgabcdefga) IGSRVEALTDVI (defgabcdefga) |
| 5iig | 70-81:A 102-113:A 163-174:A | VFRRVKEVQEQV (defgabcdefga) LEEELSDIIADV (defgabcdefga) LVVKISQLYDIA (defgabcdefga) |

| | | |
|---|---|---|
| 5ir4 | 103-121:A 132-150:A 174-192:A | VDWLLKTAGVIVELIVNFV(defgabcdefgabcdefga) FERIAAGLSGDLEAARQVH(defgabcdefgabcdefga) LQTRVIALLTRVGLLVDDI(defgabcdefgabcdefga) |
| 5j0j | 5-30:A 42-70:A 46-67:C | IKETLKRLEDSLRELRRILEELKEML(defgabcdefgabcdefgabcdefga) IVEVLKVIVKAIEASVENQRISAENQKAL(abcdefgabcdefgabcdefgabcdefga) LKVIVKAIEASVENQRISAENQ(defgabcdefgabcdefgabcd) |
| 5j1h | 758-769:B 787-811:B 896-921:B | VREAEGQLQKLQ(abcdefgabcde) LEDLLQDAQDEKEQLNEYKGHLSGL(abcdefgabcdefgabcdefgabcd) VTRLEAQHQALVTLWHQLHVDMKSLL(abcdefgabcdefgabcdefgabcde) |
| 5j1i | 1013-1034:A 1046-1067:A 1090-1119:A | ISELKDIRLQLEACETRTVHRL(efgabcdefgabcdefgabcde) CAQRIAEQQKAQAEVEGLGKGV(abcdefgabcdefgabcdefga) LRSELELTLGKLEQVRSLSAIYLEKLKTIS(defgabcdefgabcdefgabcdefgabcde) |
| 5j2l | 10-34:B 41-69:B 48-73:A | QRELDKQRKDTEEIRKRLKEIQRLT(efgabcdefgabcdefgabcdefga) ADELIKELREIIRRLQEQSEKLREIIEEL(abcdefgabcdefgabcdefgabcdefga) LREIIRRLQEQSEKLREIIEELEKII(defgabcdefgabcdefgabcdefga) |
| 5j2l | 41-69:B 6-34:A 44-72:A | ADELIKELREIIRRLQEQSEKLREIIEEL(defgabcdefgabcdefgabcdefgabcd) LREAQRELDKQRKDTEEIRKRLKEIQRLT(abcdefgabcdefgabcdefgabcdefga) LIKELREIIRRLQEQSEKLREIIEELEKI(defgabcdefgabcdefgabcdefgabcd) |
| 5l1x | 456-481:H 119-165:J 120-162:L | FNVALDQVFESIENSQALVDQSNRIL(defgabcdefgabcdefgabcdefga) TAGVAIAKTIRLESEVTAIKNALKKTNEAVSTLGNGVRVLATAVREL(defgabcdefgabcdefgabcdefgabcdefgabcdefgabcdefga) AGVAIAKTIRLESEVTAIKNALKKTNEAVSTLGNGVRVLATAV(defgabcdefgabcdefgabcdefgabcdefgabcdefgabcd) |
| 5lki | 1857-1868:C 1879-1890:C 1959-1970:C | LDLLIARGDHAY(defgabcdefga) AKMWYMQALHLL(defgabcdefga) YWQTLAQRVYNL(defgabcdefga) |
| 5mj3 | 7-25:C 51-69:C 95-113:C | IQKEIAQIQAVIAGIQKYI(defgabcdefgabcdefga) IQKQIAAIQXQIAAIQKQI(defgabcdefgabcdefga) IQKQIAAIQEQILAIYKQI(defgabcdefgabcdefga) |

| | | |
|---|---|---|
| 5mjy | 243-254:B 273-286:B 298-309:B | MKIYYFAAVAHL(defgabcdefga) FQSALDKLNEAIKL(abcdefgabcdefg) LRFTMDVIGGKY(abcdefgabcde) |
| 5n77 | 131-162:A 174-203:A 212-244:A | ELLLDLFETKIEQLADEIENIYSDLEQLSRVI(efgabcdefgabcdefgabcdefgabcdefga) ALSTLAELEDIGWKVRLCLMDTQRALNFLV(defgabcdefgabcdefgabcdefgabcde) QLEQAREILRDIESLLPHNESLFQKVNFLMQAA(defgabcdefgabcdefgabcdefgabcdefga) |
| 5t5s | 14-25:A 35-46:A 59-70:A | LKKCLSVMEAKV(defgabcdefga) VQREIADLGEAL(defgabcdefga) LRETLKSLKKVM(defgabcdefga) |
| 5vjx | 15-43:U 17-46:V 21-42:W | ITLKTKELIRQNQATQAELDQLKEQTQXF(defgabcdefgabcdefgabcdefgabcd) LKDQLEQRTRXIEANIHRQQEELRKIQEQL(gabcdefgabcdefgabcdefgabcdefga) LEQRTRXIEANIHRQQEELRKI(defgabcdefgabcdefgabcd) |
| 5xdb | 249-260:A 309-320:A 337-348:A | PITQLIFTHLYL(defgabcdefga) AEAGAREVALRV(defgabcdefga) LMVQVASAKIVA(defgabcdefga) |
| 5xdb | 249-267:C 302-320:C 337-355:C | PITQLIFTHLYLGIARGAL(defgabcdefgabcdefga) FAAQLQVAEAGAREVALRV(defgabcdefgabcdefga) LMVQVASAKIVATRLVIEL(defgabcdefgabcdefga) |
| 5xsj | 141-156:L 171-186:L 197-211:L | ITNEIKQHVDSSLDNF(gabcdefgabcdefga) YNNEVILAKQKIGNLK(defgabcdefgabcde) LRDLDNTLDSYIESS(abcdefgabcdefga) |
| 5z2c | 294-311:A 325-336:A 350-367:A | LFVLTAVNIRGTCLLSYS(abcdefgabcdefgabcd) LCEAKEAFEIGL(abcdefgabcde) ELHSFVKAAFGLTTVHRR(abcdefgabcdefgabcd) |
| 5z2c | 194-205:B 213-224:B 234-245:B | SVCIQIRGQILQ(defgabcdefga) AAELIWASIVGY(defgabcdefga) GLSTSLGILADI(defgabcdefga) |
| 5zhy | 10-49:A 108-132:A 6-52:C | FNKAMTNIVDAFTGVNDAITQTSQALQTVATALNKIQDVV(gabcdefgabcdefgabcdefgabcdefgabcdefgabcd) ISTLENKSAELNYTVQKLQTLIDNI(abcdefgabcdefgabcdefgabcd) LAASFNKAMTNIVDAFTGVNDAITQTSQALQTVATALNKIQDVVNQQ(defgabcdefgabcdefgabcdefgabcdefgabcdefgabcdefga) |
| 6c12 | 459-471:B 483-494:B 516-527:B | IRKALQECMQHNF(defgabcdefgab) GLEQLKVIRERL(defgabcdefga) LDNLMETAYATA(defgabcdefga) |

| | | |
|---|---|---|
| 6dkm | 45-69:C 87-111:D 121-141:D | ERLLEELLRILDENAELLKRNLELL(abcdefgabcdefgabcdefgabcd)<br>LREYHRVLREYEKLLEELRRLYEEY(abcdefgabcdefgabcdefgabcd)<br>SDRILREIKEILDKSERLWDL(abcdefgabcdefgabcdefg) |
| 6ds9 | 7-25:A 35-56:A 70-92:A | FKQRLAAIKTRLAAIKTRL(defgabcdefgabcdefga)<br>LAAFEKEIAAFESEIAAFESEL(abcdefgabcdefgabcdefga)<br>LRKEAAAIRDEAAAIRDELQAYR(defgabcdefgabcdefgabcde) |
| 6efk | 94-105:B 110-123:B 135-146:B | VKAHFFLGQCQL(defgabcdefga)<br>YDEAIANLQRAYSL(abcdefgabcdefg)<br>IPSALRIAKKKR(abcdefgabcde) |
| 6gy7 | 298-316:A 119-133:D 302-319:D | LTEAVKIEQVWISFAEQLH(gabcdefgabcdefgabcd)<br>DNVIVSFTGRTSKLT(abcdefgabcdefga)<br>VKIEQVWISFAEQLHKLS(abcdefgabcdefgabcd) |
| 6gyp | 344-355:B 373-384:B 403-414:B | LYWKIISLDRDL(defgabcdefga)<br>IRRELDIFQYKV(defgabcdefga)<br>ALFQISTVSWKL(defgabcdefga) |
| 6mbw | 151-162:A 229-240:A 285-296:A | LRLVTQDTENEL(defgabcdefga)<br>LAEKHQKTLQLL(defgabcdefga)<br>IIWQNRQQIRRA(defgabcdefga) |
| 6mbz | 230-244:B 282-296:B 313-327:B | AEKHQKTLQLLRKQQ(abcdefgabcdefga)<br>LAEIIWQNRQQIRRA(defgabcdefgabcd)<br>LAEVNATITDIISAL(abcdefgabcdefga) |
| 6n9h_ba1 | 2-31:A 43-74:A 40-65:C | AQDKLKYLVKQLERALRELKKSLDELERSL(gabcdefgabcdefgabcdefgabcdefga)<br>LVENNRLNVENNKIIVEVLRIILELAKASAKL(abcdefgabcdefgabcdefgabcdefgabcd)<br>EDALVENNRLNVENNKIIVEVLRIIL(defgabcdefgabcdefgabcdefga) |
| 6nqx | 103-114:C 164-175:C 192-203:C | ALSNYQSGVNTI(defgabcdefga)<br>ALNYFSKAINHL(defgabcdefga)<br>LLSDVYRSWIMA(defgabcdefga) |
| 6nt9 | 450-471:A 505-526:A 626-647:A | IKDDYNETVHKKTEVVITLDFC(defgabcdefgabcdefgabcd)<br>LLRLSSSQGTIETSLQDIDSRL(abcdefgabcdefgabcdefga)<br>LRKQLLSLTNQCFDIEEEVSKY(defgabcdefgabcdefgabcd) |
| 6nz3 | 27-86:A 4-21:B 30-89:B | EKQIVEAIRAIVENNAQIVEAIRAIVENNAQIVENNRAIIEALEAIGGHNKILEEMKKQL(defgabcdefgabcdefgabcdefgabcdefgabcdefgabcdefgabcdefgabcdefg)<br>MGDLKYSLERLREILERL(abcdefgabcdefgabcd)<br>IVEAIRAIVENNAQIVEAIRAIVENNAQIVENNRAIIEALEAI |

| | | |
|---|---|---|
| | | GGHNKILEEMKKQLKDL(abcdefgabcdefgabcdefgabcdefgabcdefgabcdefgabcdefgabcdefgabcd) |
| 6poo | 307-329:A 342-353:A 367-388:A | YAKKIERISSKGLALSKKAKEIY(gabcdefgabcdefgabcdefga) YADSVGTYLNRF(defgabcdefga) GQSGLDEAKKXLDEVKKLLKEL(abcdefgabcdefgabcdefga) |
| 6q6g | 445-456:Q 461-474:Q 481-492:Q | EPLLNNLGHVCR(defgabcdefga) YAEALDYHRQALVL(abcdefgabcdefg) TYSAIGYIHSLM(abcdefgabcde) |
| 6qdl | 81-92:A 97-110:A 117-128:A | VKALFRRSLARE(defgabcdefga) VGPAFQDAKEALRL(abcdefgabcdefg) IVEVLQRLVKAN(abcdefgabcde) |
| 6qti | 657-668:A 709-720:A 788-799:A | PQLVAAFHSLVG(defgabcdefga) TFSGSLVAYGKL(defgabcdefga) MPVVITVLNSYS(defgabcdefga) |
| 6r17 | 63-143:A 67-143:B 78-103:D | LDSSIDILQKRAQELIENINKSRQKDHALMTNFRNSLKTKVSDLTEKLEERIYQIYNDHNKIIQEKLQEFTQKMAKISHLE(defgabcdefgabcdefgabcdefgabcdefgabcdefgabcdefgabcdefgabcdefgabcdefgabcdefgabcdefg) IDILQKRAQELIENINKSRQKDHALMTNFRNSLKTKVSDLTEKLEERIYQIYNDHNKIIQEKLQEFTQKMAKISHLE(bcdefgabcdefgabcdefgabcdefgabcdefgabcdefgabcdefgabcdefgabcdefgabcdefgabcdefga) RAAVDASYIDEIDELFKEANAIENFL(defgabcdefgabcdefgabcdefga) |
| 6r80 | 936-951:A 994-1005:A 1145-1159:A | FEKAVYYLDAVVSFIE(defgabcdefgabcde) ADKRLTVLCLRC(defgabcdefga) VRYTRQGLHWLRQDA(abcdefgabcdefga) |
| 6re5 | 241-252:2 262-273:2 284-295:2 | PAQAVEAAYGLA(defgabcdefga) FKALFGVVAPAI(defgabcdefga) SLAQLHVASTIS(defgabcdefga) |
| 6rw6 | 2019-2043:C 2255-2279:C 2307-2318:C | FPHMLENARGMVSQLTQFGSTLQNI(defgabcdefgabcdefgabcdefg) RGRLAAIYFQFYDLAVARCLMAEQA(defgabcdefgabcdefgabcdefg) GETLMLSLAQME(abcdefgabcde) |
| 6rw9 | 1988-2005:A 2223-2238:A 2272-2283:A | LDSARSLTGQLMQFGSTL(abcdefgabcdefgabcd) LASIYYRFYDLTAARC(gabcdefgabcdefga) GESLLLNLAEME(abcdefgabcde) |

| | | |
|---|---|---|
| 6sz9 | 59-80:D 107-128:D 240-261:D | NVLNGIVIQQASDYHVYAQKLL(abcdefgabcdefgabcdefga) ERQRLVQLGDEFHKLELEQDNL(defgabcdefgabcdefgabcd) TNHLSQQAKSFRTQFYEVILRI(abcdefgabcdefgabcdefga) |
| 6t46_ba1 | 133-144:G 149-163:G 173-187:G | AEFHYKIGVAYY(defgabcdefga) HLVSVNKVTKARDIY(abcdefgabcdefga) AIQCSLVVGINLYDM(efgabcdefgabcde) |
| 6t46_ba1 | 170-184:G 190-207:G 212-227:G | NLEAIQCSLVVGINL(gabcdefgabcdefg) LDDADAYFRDALTEALDH(abcdefgabcdefgabcd) PITKIYHNLGLVHWQK(defgabcdefgabcde) |
| 6tms | 8-29:E 41-66:E 40-64:F | LRKLLEEAEKKLYKLEDKTRRS(defgabcdefgabcdefgabcd) AQSLQLIAESLMLIAESLLIIAISLL(defgabcdefgabcdefgabcdefga) KAQSLQLIAESLMLIAESLLIIAIS(efgabcdefgabcdefgabcdefga) |
| 6vbu | 135-146:4 151-164:4 170-181:4 | WEICHNLGVCYI(defgabcdefga) FDKAQDQLHNALHL(abcdefgabcdefg) TYIMLGKIFLLK(abcdefgabcde) |
| 6vbu | 451-462:8 467-480:8 487-498:8 | YEPHFNFATISD(defgabcdefga) LQRSYAAAKKSEAA(abcdefgabcdefg) TQHLIKQLEQHF(abcdefgabcde) |
| 6w2q | 68-82:A 91-105:A 113-127:A | PEAIIAAARALLKIA(defgabcdefgabcd) AKQAIEAASKAAQLA(abcdefgabcdefga) LVCEALALLIAAQVL(defgabcdefgabcd) |
| 6wa0 | 6-26:A 5-26:B 9-26:C | LLVAFVAYYTALIALIFAILA(efgabcdefgabcdefgabcd) ALLVAFVAYYTALIALIFAILA(defgabcdefgabcdefgabcd) AFVAYYTALIALIFAILA(abcdefgabcdefgabcd) |
| 6wc3 | 4-17:B 31-49:B 80-91:B | ATQLALLQDELLDM(abcdefgabcdefg) IIDKTLRFRELLGCYRLQV(defgabcdefgabcdefga) ALAQLLLWERFL(defgabcdefga) |
| 6xf1 | 1432-1450:A 1460-1490:A 1512-1537:A | CIFKNNELLKNIQDVQSQI(defgabcdefgabcdefga) VPAVKHRKKSLIRLDKVLDEYEEEKRHLQEM(efgabcdefgabcdefgabcdefgabcdefg) TVVLWENTKALVTECLEQCGRVLELL(defgabcdefgabcdefgabcdefga) |
| 6xns | 192-206:B 221-235:B 251-265:B | LKAVETVVKVARALN(abcdefgabcdefga) AARVASEAARLAERV(defgabcdefgabcd) ARELQEKVLDILLDI(abcdefgabcdefga) |

| | | |
|---|---|---|
| 6xr1 | 3-24:A 32-54:A 62-80:A | LELALKALQILVNAAYVLAEIA(defgabcdefgabcdefgabcd) LLEKAARLAEEAARQAEEIARQA(gabcdefgabcdefgabcdefga) LALKALQILVNAAYVLAEI(defgabcdefgabcdefga) |
| 6xt4_ba1 | 191-208:A 217-245:A 218-242:C | LLILILILLDLKEMLERL(abcdefgabcdefgabcd) IVKVLKVIVKAIEASVLNQAISAINQILL(abcdefgabcdefgabcdefgabcdefga) VKVLKVIVKAIEASVLNQAISAINQ(abcdefgabcdefgabcdefgabcd) |
| 6zl1 | 16-27:C 50-61:C 96-107:C | IQKRIAAIQKRI(defgabcdefga) ITKQIAAIQLRI(defgabcdefga) IETQICKIEAAI(defgabcdefga) |
| 7ag9 | 170-186:B 231-242:B 282-296:B | DDLINKNVEKCFEIQEL(efgabcdefgabcdefg) LSRKFLILKRNI(defgabcdefga) IMKNFRFMMNEIKDL(defgabcdefgabcd) |
| 7ag9 | 309-330:B 344-362:B 375-389:B | FINLNNELEYIIEEVRLLLKKI(defgabcdefgabcdefgabcd) FNSQLAKKSKIITKTFNII(gabcdefgabcdefgabcd) IALKTNELAKVWVDL(defgabcdefgabcd) |
| 7bjs | 857-910:A 857-906:B 282-314:C | LENNLDQLTKVHKQLVRDNADLRCELPKLEKRLRCTMERVKALETALKEAKEGA(defgabcdefgabcdefgabcdefgabcdefgabcdefgabcdefgabcdefga) LENNLDQLTKVHKQLVRDNADLRCELPKLEKRLRCTMERVKALETALKEA(defgabcdefgabcdefgabcdefgabcdefgabcdefgabcdefgabcd) AESEVAALNRRIQLLEEDLERSEERLGSATAKL(defgabcdefgabcdefgabcdefgabcdefga) |
| 7d3u | 332-346:D 369-380:D 531-544:D | HHMIVKAALFLAIGA(defgabcdefgabcd) VAVAFFASAMSL(abcdefgabcde) AAPALALSVVTLAL(abcdefgabcdefg) |
| 7dhg | 187-198:C 203-216:C 224-235:C | VKALFRRAKAHE(defgabcdefga) KKECLEDVTAVCIL(abcdefgabcdefg) SMLLADKVLKLL(abcdefgabcde) |
| 7drw | 104-125:M 147-167:M 202-216:M | LALVKPEVWTLKEKCILVITWI(abcdefgabcdefgabcdefga) LERVNAVKTKVEAFQTTISKY(efgabcdefgabcdefgabcd) LRAMVLDLRAFYAEL(abcdefgabcdefga) |
| 7kam | 122-133:F 138-151:F 159-170:F | SGLYANRAQAHM(defgabcdefga) WPEGWVDAKCSVES(abcdefgabcdefg) GWWRGGKCLVEM(abcdefgabcde) |
| 7n7p | 12-65:A 83-94:A 12-61:B | CLRDWEDLQQDFQNIQETHRLYRLKLEELTKLQNNCTSSITRQKKRLQELALAL(defgabcdefgabcdefgabcdefgabcdefgabcdefgabcdefgabcdefga) LENQMKERQGLF(defgabcdefga) CLRDWEDLQQDFQNIQETHRLYRLKLEELTKLQNNCTSSITRQ |

| | | |
|---|---|---|
| | | KKRLQEL(defgabcdefgabcdefgabcdefgabcdefgabc defgabcdefgabcd) |
| 7nj0 | 813-824:A 832-843:A 856-867:A | AGSSCHITQLLL(defgabcdefga) AQLHLEEAASSL(defgabcdefga) LSLTCDLLRSQL(defgabcdefga) |
| 7o3v | 122-158:B 168-213:B 169-205:C | MRRSIEERSRTAAATDKAVGLRAYEGAQQRLAQIEGL(cdefg abcdefgabcdefgabcdefgabcdefgabcd) QKAIEELQARIAGEQAAIQNETTKLQMIAQLRQAEQALISEQR RER(efgabcdefgabcdefgabcdefgabcdefgabcdefga bcdefga) KAIEELQARIAGEQAAIQNETTKLQMIAQLRQAEQAL(defga bcdefgabcdefgabcdefgabcdefgabcde) |
| 7opb | 2-13:D 21-32:D 42-53:D | VIEKLRKLEKQA(defgabcdefga) LVMLARMVLEYL(defgabcdefga) ADESADRIEEVL(defgabcdefga) |
| 7qpg | 20-38:W 59-77:W 95-113:W | LGTRISRLTRRVEEIKGEV(defgabcdefgabcdefga) LITQVDKLSEDIDLLKSRI(defgabcdefgabcdefga) LKQQLERDSVVLSLLKQLQ(defgabcdefgabcdefga) |
| 7sil | 620-631:A 655-668:A 838-849:A | AVLGIFLTAFVL(abcdefgabcde) LFSLLCCFSSSLFF(abcdefgabcdefg) VIAILAASFGLL(defgabcdefga) |
| 7v1m | 85-99:G 176-190:G 201-215:G | GEAFFFYGKSLLELA(defgabcdefgabcd) LELAWDMLDLAKIIF(abcdefgabcdefga) YAAQAHLKLGEVSVE(defgabcdefgabcd) |
| 7vrb | 7-18:A 63-80:A 89-107:A | QLHQLRAQIMAY(defgabcdefga) IQKMLDDNNHLIQCIMDS(defgabcdefgabcdefg) CSQYQQMLHTNLVYLATIA(defgabcdefgabcdefga) |
| 7w5m | 282-293:A 298-323:A 358-383:A | AELNFRICICLE(defgabcdefga) PKEAIPYCQKALLICKARMERLSNEI(abcdefgabcdefgab cdefgabcde) KEVEIGDLAGLAEDLEKKLEDLKQQA(abcdefgabcdefgab cdefgabcde) |

Table S3. Sequences of *de novo* peptides and proteins in this study

| Systematic name | Amino acid sequence<br>*gabdcef gabdcef gabdcef gabdcef* | Mass (kDa) | ε |
|---|---|---|---|
| apCCTri-A | Ac-G -LAALEK ELAA**T**EK ELAALEW ELAALE- G-$NH_2$ | 2979.58 | 5690 ($ε_{280}$) |
| apCCTri-B | Ac-G -LAALKE KLAALKE K**N**AALKY KLAALK- G-$NH_2$ | 2964.82 | 1280 ($ε_{275}$) |
| apCCTri-N | Ac-G KLAALEK KLAALEK K**T**AALEY KLAAL-- G-$NH_2$ | 2952.77 | 1280 ($ε_{275}$) |
| apCCTri-N' | Ac-G ELAALKE ELAALKE E**T**AALKY ELAAL-- G-$NH_2$ | 2955.62 | 1280 ($ε_{275}$) |
| apCCTri-A-c4CF | Ac-G -LAALEK ELAA**T**EK ELAALEQ ELAAL**Z**- G-$NH_2$ | 2964.58 | 32393 ($ε_{214}$) |
| apCCTri-A-n4CF | Ac-G **Z**LAALEK ELAA**T**EK ELAALEQ ELAALE- G-$NH_2$ | 3093.62 | 33394 ($ε_{214}$) |
| apCCTri-B-nMSE | Ac-G -**X**AALKE KLAALKE K**N**AALKQ KLAALK- G-$NH_2$ | 2995.72 | 28116 ($ε_{214}$) |
| apCCTri-N-nMSE | Ac-G K**X**AALEK KLAALEK K**T**AALEQ KLAAL-- G-$NH_2$ | 2982.63 | 28058 ($ε_{214}$) |
| X = Selenomethionine (MSE), Z = 4-cyanophenyalanine (4CF) | | | |
| sc-apCC3-ABN / CW1 | MGSSHHHHHHSSGLVPRGSHM<br>MLAALKE KLAALKE K**N**AALKY KLAALK EKLGLTPE<br>LAALEK ELAA**T**EK ELAALEW ELAALE ADPNPDPA<br>KLAALEK KLAALEK K**T**AALEY KLAAL | 12472.7 | 8250 ($ε_{280}$) |
| sc-apCC3-ABN / CW2 | MGSSHHHHHHSSGLVPRGSHM<br>MLAALKE KLAALKE K**N**AALKY KLAALK KEKGETPE<br>LAALEK ELAA**T**EK ELAALEW ELAALE ADPNPDPA<br>KLAALEK KLAALEK K**T**AALEY KLAAL | 12503.6 | 8250 ($ε_{280}$) |
| sc-apCC3-ABN' / ACW | MGSSHHHHHHSSGLVPRGSHM<br>MLAALKE KLAALKE K**N**AALKY KLAALK KHKATPAE<br>LAALEK ELAA**T**EK ELAALEW ELAALE KKEPLTP<br>ELAALKE ELAALKE E**T**AALKY ELAAL | 12486.6 | 8250 ($ε_{280}$) |
| | MGSSHHHHHHSSGLVPRGSHM | | |

| | | | |
|---|---|---|---|
| sc-apCC3-CW-flex | MLAALKE KLAALKE KNAALKY KLAALK GSGSGS | 11964.9 | 8250 (ε280) |
| | LAALEK ELAATEK ELAALEW ELAALE GSGSGSGS | | |
| | KLAALEK KLAALEK KTAALEY KLAAL | | |
| sc-apCC3-ACW-flex | MGSSHHHHHHSSGLVPRGSHM | 11967.7 | 8250 (ε280) |
| | MLAALKE KLAALKE KNAALKY KLAALK GSGSGS | | |
| | LAALEK ELAATEK ELAALEW ELAALE GSGSGSGS | | |
| | ELAALKE ELAALKE ETAALKY ELAAL | | |
| sc-apCC3-CW-mismatch | MGSSHHHHHHSSGLVPRGSHM | 12483.8 | 8250 (ε280) |
| | MLAALKE KLAALKE KNAALKY KLAALK AEKATPEE | | |
| | LAALEK ELAATEK ELAALEW ELAALE KKEPLTP | | |
| | KLAALEK KLAALEK KTAALEY KLAAL | | |
| sc-apCC3-ACW-mismatch | MGSSHHHHHHSSGLVPRGSHM | 12506.5 | 8250 (ε280) |
| | MLAALKE KLAALKE KNAALKY KLAALK EKLGLTPE | | |
| | LAALEK ELAATEK ELAALEW ELAALE ADPNPDPA | | |
| | ELAALKE ELAALKE ETAALKY ELAAL | | |
| sc-apCC3-CW-0NTT | MGSSHHHHHHSSGLVPRGSHM | 13087 | 8250 (ε280) |
| | MLAALKE KLAALKE KLAALKY KLAALK EKLGLTPE | | |
| | LAALEK ELAALEK ELAALEW ELAALE ADPNPDPA | | |
| | ELAALKE ELAALKE ELAALKY ELAAL | | |
| sc-apCC3-CW-2NTT | MGSSHHHHHHSSGLVPRGSHM | 12449.5 | 8250 (ε280) |
| | MLAALKE K**N**AALKE K**N**AALKY KLAALK EKLGLTPE | | |
| | LAALEK ELAA**T**EK ELAA**T**EW ELAALE ADPNPDPA | | |
| | KLAALEK K**T**AALEK K**T**AALEY KLAAL | | |
| sc-apCC3-CW-3NTT | MGSSHHHHHHSSGLVPRGSHM | 12426.3 | 8250 (ε280) |
| | M**N**AALKE K**N**AALKE K**N**AALKY KLAALK EKLGLTPE | | |
| | LAALEK ELAA**T**EK ELAA**T**EW ELAA**T**E ADPNPDPA | | |
| | E**T**AALEK E**T**AALEK E**T**AALKY ELAAL | | |
| sc-apCC3-CW-4NTT | MGSSHHHHHHSSGLVPRGSHM | 12403.2 | 8250 (ε280) |
| | M**N**AALKE K**N**AALKE K**N**AALKY K**N**AALK EKLGLTPE | | |
| | LAA**T**EK ELAA**T**EK ELAA**T**EW ELAA**T**E ADPNPDPA | | |
| | E**T**AALEK E**T**AALEK E**T**AALKY E**T**AAL | | |
| | MLAALKE KLAALKE KNAALKY KLAALK EKLGLTPE | | |
| | LAALEK ELAATEK ELAALEW ELAALE ADPNPDPA | | |

| | |
|---|---|
| | EAAAAKE EAAAAEK ETAAAKY EAAAA |

| Systematic name | synthesis / host organism | gene synthesis | Insertion site | Vector | Insert length | Construct length |
|---|---|---|---|---|---|---|
| sc-apCC3-CW1 | Escherichia coli | clonal | NdeI_XhoI | pET-28a(+) | 308 | 5600 |
| sc-apCC3-CW2 | Escherichia coli | clonal | NdeI_XhoI | pET-28a(+) | 308 | 5600 |
| sc-apCC3-ACW | Escherichia coli | clonal | NdeI_XhoI | pET-28a(+) | 309 | 5601 |
| sc-apCC3-CW-flex | Escherichia coli | clonal | NdeI_XhoI | pET-28a(+) | 309 | 5601 |
| sc-apCC3-ACW-flex | Escherichia coli | clonal | NdeI_XhoI | pET-28a(+) | 309 | 5601 |
| sc-apCC3-CW-mismatch | Escherichia coli | clonal | NdeI_XhoI | pET-28a(+) | 309 | 5601 |
| sc-apCC3-ACW-mismatch | Escherichia coli | clonal | NdeI_XhoI | pET-28a(+) | 309 | 5601 |
| sc-apCC3-CW-0NTT | Escherichia coli | clonal | NdeI_XhoI | pET-28a(+) | 312 | 5604 |
| sc-apCC3-CW-2NTT | Escherichia coli | clonal | NdeI_XhoI | pET-28a(+) | 312 | 5604 |
| sc-apCC3-CW-3NTT | Escherichia coli | clonal | NdeI_XhoI | pET-28a(+) | 312 | 5604 |
| sc-apCC3-CW-4NTT | Escherichia coli | clonal | NdeI_XhoI | pET-28a(+) | 312 | 5604 |

| Systematic name | DNA sequence |
|---|---|
| sc-apCC3-CW-1 | ATGCTTGCAGCATTAAAGGAGAAACTCGCAGCGCTGAAAGAGAAGAATGCGGCTCTGAAATACAAACTCGCCGCCCTGAAAGAGAAGTTGGGGCTGACTCCTGAGTTGGCAGCCCTTGAGAAAGAGTTAGCGGCGACTGAGAAGGAACTGGCGGCCCTGGAATGGGAGTTGGCCGCTTTAGAGGCAGACCCAAATCCTGATCCAGCGAAGTTAGCTGCCCTTGAGAAGAAGTTGGCAGCACTGGAGAAGAAGACGGCAGCCCTGGAGTACAAGCTTGCAGCGCTCTGA |
| sc-apCC3-CW-2 | ATGCTCGCAGCACTCAAGGAGAAACTTGCAGCCTTAAAGGAAAAGAACGCGGCCCTCAAGTACAAGTTAGCAGCTTTGAAGAAAGAAAAGGGTGAGACACCTGAACTGGCTGCGCTGGAGAAGGAATTAGCCGCGACCGAGAAAGAGTTGGCAGCACTGGAATGGGAACTCGCAGCTTTAGAGGCCGATCCTAACCCAGACCCTGCAAAGCTCGCGGCCTTAGAAAAGAAGCTCGCTGCCTTAGAAAAGAAGACAGCCGCGCTTGAGTACAAGCTTGCGGCGCTGTGA |
| sc-apCC3-ACW | ATGCTGGCAGCTCTTAAAGAAAAACTGGCAGCACTCAAAGAAAAGAACGCAGCTCTGAAGTACAAACTGGCTGCTCTTAAAAAACATAAAGCCACGCCAGCGGAACTGGCAGCATTGGAAAAAGAGCTGGCCGCCACAGAAAAGGAGCTGGCGGCATTAGAGTGGGAATTAGCGGCCCTTGAAAAAAAAGAACCGCTTACTCCCGAATTGGCAGCCTTGAAAGAAGAATTGGCAGCCTTAAAAGAAGAGACAGCGGCCCTGAAGTATGAACTTGCGGCACTGTGA |
| sc-apCC3-CW-flex | ATGCTTGCGGCACTCAAAGAGAAATTAGCAGCATTAAAGGAAAAGAACGCCGCTCTGAAGTACAAATTAGCTGCCCTGAAAGGCTCTGGTAGCGGTAGTGAGCTGGCTGCACTCGAAAAAGAACTGGCGGCGACGGAAAAGGAACTGGCGGCTCTCGAATGGGAACTGGCTGCACTTGAAGGGTCAGGCTCGGGTTCCGGAAGCAAGCTGGCCGCTCTGGAAAAGAAACTGGCGGCGCTGGAGAAGAAAACCGCTGCGTTAGAATATAAACTCGCCGCGCTGTGA |

| | |
|---|---|
| sc-apCC3-ACW-flex | ATGCTGGCTGCCCTGAAGGAGAAGCTGGCGGCTCTGAAAGAAAAGAACGCCGCGCTTAAATACAAGCTCGCTGCGTTAAAAGGTAGCGGATCCGGTAGTGAACTCGCCGCTCTTGAAAAGGAACTTGCAGCGACGGAAAAAGAATTGGCGGCATTAGAGTGGGAACTCGCCGCACTGGAAGGCAGCGGCTCTGGTAGCGGCTCCGAACTGGCGGCACTTAAAGAAGAATTAGCTGCACTTAAGGAAGAAACAGCCGCGTTGAAGTATGAACTGGCGGCTCTTTAA |
| sc-apCC3-CW-mismatch | ATGTTGGCCGCGTTAAAAGAAAAACTTGCTGCGCTGAAAGAAAAAAACGCGGCTTTAAAATATAAGTTGGCGGCATTAAAAAAGCATAAGGCGACTCCTGCCGAATTAGCAGCCCTTGAAAAAGAATTAGCGGCCACCGAGAAAGAACTGGCGGCACTGGAATGGGAATTGGCGGCGCTGGAGAAGAAGGAGCCTCTGACACCGAAATTAGCGGCATTGGAGAAAAAATTGGCCGCGCTCGAAAAAAAAACTGCGGCGTTAGAATATAAACTGGCCGCGCTTTAG |
| sc-apCC3-ACW-mismatch | ATGCTGGCCGCCCTCAAAGAGAAACTTGCGGCACTCAAAGAGAAGAACGCGGCGCTGAAATATAAATTAGCCGCACTGAAAAAAGAAAAAGGGGAGACTCCTGAGCTGGCGGCACTGGAAAAAGAACTGGCGGCCACGGAGAAAGAACTGGCGGCGTTAGAGTGGGAGCTGGCCGCCCTGGAAGCCGATCCAAATCCAGATCCCGCTGAGCTGGCAGCCCTTAAAGAGGAATTGGCTGCGTTAAAGGAGGAAACGGCGGCGCTGAAATATGAACTGGCAGCTTTGTAA |
| sc-apCC3-CW-0NTT | ATGCTCGCGGCGCTGAAAGAGAAGCTGGCCGCGCTCAAAGAGAAGCTGGCCGCACTTAAATACAAGCTGGCAGCCCTGAAAGAGAAATTAGGCTTAACACCAGAGCTCGCTGCACTCGAAAAGGAACTGGCCGCTCTTGAAAAAGAACTGGCAGCGCTCGAATGGGAGTTAGCTGCACTGGAAGCTGATCCGAACCCAGATCCGGCGAAGCTGGCGGCCCTCGAAAAAAAGCTTGCGGCGTTAGAAAAGAAGCTGGCCGCGCTGGAATATAAACTCGCAGCTCTTTAA |

| | |
|---|---|
| sc-apCC3-CW-2NTT | ATGCTTGCAGCGCTGAAGGAAAAAAATGCGGCACTTAAAGAAAAAA<br>ATGCAGCTCTGAAATATAAACTCGCCGCACTGAAGGAAAAACTGGG<br>TTTGACTCCGGAACTGGCGGCTCTGGAGAAGGAATTGGCAGCCAC<br>AGAGAAGGAGCTGGCCGCTACAGAGTGGGAACTTGCTGCACTGGA<br>GGCAGATCCGAATCCTGATCCTGCCAAATTAGCCGCACTTGAGAAG<br>AAAACGGCGGCTCTTGAAAAAAAAACGGCAGCCCTTGAGTACAAAC<br>TGGCAGCTCTTTGA |
| sc-apCC3-CW-3NTT | ATGAATGCGGCCTTGAAAGAGAAGAATGCCGCACTGAAGGAAAAAA<br>ACGCGGCACTGAAGTACAAACTTGCCGCCTTAAAGGAGAAGCTGG<br>GTTTGACCCCTGAACTCGCAGCTCTTGAGAAAGAACTGGCAGCAAC<br>CGAAAAAGAATTAGCCGCGACCGAGTGGGAGCTGGCGGCAACAGA<br>GGCGGACCCTAACCCCGATCCAGCCAAAACCGCGGCTTTAGAAAA<br>AAAAACGGCTGCACTGGAAAAAAAAACTGCCGCGCTGGAATATAAA<br>CTGGCCGCCCTGTAA |
| sc-apCC3-CW-4NTT | ATGAATGCAGCACTTAAAGAAAAAAACGCCGCATTAAAGGAAAAAAA<br>TGCCGCCTTGAAATATAAAAACGCTGCGCTGAAAGAAAAACTCGGT<br>CTTACACCGGAATTAGCCGCCACAGAGAAAGAATTAGCCGCAACTG<br>AGAAAGAACTGGCCGCAACAGAATGGGAATTAGCGGCTACCGAAG<br>CCGACCCCAATCCGGATCCGGCTAAGACAGCAGCTCTCGAAAAGA<br>AAACGGCGGCTCTGGAAAAGAAAACCGCAGCCCTTGAATATAAAAC<br>TGCGGCCTTGTAA |

Table S4. Evaluated Alphafold2 metrics for peptide assemblies, the unique 3-helix off-target combinations and proteins.

| Model | average pLDDT | pTM | max pAE | Average pAE | *i*pTM |
|---|---|---|---|---|---|
| peptide mix AAA | 96.55 | 0.88 | 16.42 | 2.21 | 0.87 |
| peptide mix AAB | 95.95 | 0.85 | 21.28 | 2.54 | 0.83 |
| peptide mix NNN | 95.50 | 0.85 | 22.13 | 2.54 | 0.83 |
| peptide mix ABB | 95.20 | 0.85 | 22.33 | 2.81 | 0.83 |
| peptide mix ABN | 95.18 | 0.85 | 23.39 | 2.69 | 0.83 |
| peptide mix AAN | 95.09 | 0.85 | 20.36 | 2.71 | 0.83 |
| peptide mix ANN | 94.81 | 0.84 | 23.64 | 2.71 | 0.82 |
| peptide mix BBB | 94.71 | 0.85 | 23.94 | 2.84 | 0.83 |
| peptide mix BNN | 94.60 | 0.85 | 25.69 | 2.80 | 0.83 |
| peptide mix BBN | 94.14 | 0.85 | 22.42 | 2.90 | 0.83 |
| sc-apCC3-ACW | 85.20 | 0.65 | 30.98 | 10.06 | |
| sc-apCC3-ACW-flex | 84.37 | 0.67 | 31.30 | 10.33 | |
| sc-apCC3-ACW-mismatch | 83.57 | 0.65 | 31.31 | 10.17 | |
| sc-apCC3-CW1 | 85.31 | 0.65 | 31.25 | 9.35 | |
| sc-apCC3-CW2 | 85.65 | 0.65 | 31.30 | 9.36 | |
| sc-apCC3-CW-flex | 82.48 | 0.65 | 31.31 | 10.67 | |
| sc-apCC3-CW-mismatch | 79.39 | 0.63 | 31.30 | 9.97 | |
| sc-apCC3-CW-0NTT | 84.84 | 0.65 | 31.22 | 9.44 | |
| sc-apCC3-CW-2NTT | 85.51 | 0.65 | 31.25 | 9.30 | |
| sc-apCC3-CW-3NTT | 84.07 | 0.62 | 31.31 | 10.01 | |
| sc-apCC3-CW-4NTT | 83.23 | 0.68 | 31.06 | 8.78 | |

Table S5. MASTER hits for each of the designed proteins in this study

| Systematic name | MASTER hit PDB id | Hit residues | RMSD (Å) |
|---|---|---|---|
| sc-apCC3-CW | Loop1: 2fu2 | 29-35, 44-50 | 0.298 |
| | Loop2: 3zn3 | 202-208, 217-223 | 0.350 |
| sc-apCC3-ACW | Loop1: 3gae | 165-171, 179-185 | 0.528 |
| | Loop2: 2fu2 | 29-35, 43-49 | 0.495 |

Table S6. Summary of biophysical characterization

| systematic name | CD % helicity$_{222}$ | $T_M$ (°C) | SV weight (Observed / expected) | SE weight (Observed / expected) | Expected mass (Da) |
|---|---|---|---|---|---|
| Individual peptides | | | | | |
| peptide A | 20.1 | <10.0 | n.d. | n.d. | 2979.58 |
| peptide B | 42.7 | <15.0 | 1.2 | n.d. | 2964.82 |
| peptide N | 78.7 | 79.0 | 2.5 | 2.3 | 2952.77 |
| peptide N' | 71.1 | 65.0 | 2.1 | 2.9 | 2955.62 |
| Peptide assemblies (complex) | | | | | |
| apCCTri-ABN-CW | 90.9 | 78.0 | 1.2 | 1.0 | 8897.17 |
| apCCTri-ABN'-ACW | 61.9 | 65.0 | 0.8 | 1.2 | 8900.02 |
| Proteins | | | | | |
| sc-apCC3-CW1 | 78.5 ± 2.4 | > 95.0 | 0.9 | 0.8 | 12472.7 |
| sc-apCC3-CW2 | 70.8 | > 95.0 | 1.0 | n.d. | 12503.6 |
| sc-apCC3-CW-flex | 68.7 ± 2.6 | > 95.0 | 0.8 | 0.8 | 11964.9 |
| sc-apCC3-CW-mismatch | 57.1 ± 2.2 | > 95.0 | 1.1 | 0.7 | 12483.8 |
| sc-apCC3-ACW | 82.2 ± 1.8 | > 95.0 | 0.9 | 0.8 | 12486.6 |
| sc-apCC3-ACW-flex | 70.2 ± 2.0 | > 95.0 | 0.8 | 0.8 | 11967.7 |
| sc-apCC3-ACW-mismatch | 53.5 ± 1.5 | > 95.0 | 0.8 | 0.9 | 12506.5 |
| sc-apCC3-CW-0NTT | 75.1 | > 95.0 | 2.2 | 0.8 | 13087.0 |
| sc-apCC3-CW-2NTT | 66.6 | > 95.0 | 0.9 | 0.8 | 12449.5 |
| sc-apCC3-CW-3NTT | 62.6 | 86.5 ± 0.8 | 0.8 | 0.8 | 12426.3 |
| sc-apCC3-CW-4NTT | 36.3 | 25.3 ± 0.7 | 1.3 | 0.8 | 12403.2 |
| Alternate states (peptides) | | | | | |
| peptide mix AAB | 50.7 | 53.0 | n.d. | n.d. | |
| peptide mix ABB | 54.3 | 56.0 | n.d. | n.d. | |
| peptide mix AAN | 29.7 | 57.0 | n.d. | n.d. | |
| peptide mix ANN | 48.6 | 65.0 | n.d. | n.d. | |
| peptide mix BNN | 28.8 | 58.0 | n.d. | n.d. | |
| peptide mix BBN | 46.7 | 63.0 | n.d. | n.d. | |

Table S7. Crystallization conditions used to obtain the X-ray structures discussed in this study

| Systematic name | Molecular dimensions screen | Crystallization conditions of the reservoir | Dilution sample / reservoir (/ seed) within the drop |
|---|---|---|---|
| sc-apCC3-CW-1 | JCSG, E1 | 1.0 M sodium citrate tribasic dihydrate, 0.1 M sodium cacodylate, pH 6.5 | 2:1 |
| sc-apCC3-CW-2 | Proplex, G4 | 2.0 M ammonium sulfate, 0.1 M Tris, pH 8.0 | 2:1 |
| sc-apCC3-CW-0NTT | Structure screen 1 and 2, G4 | 0.2 M Ammonium sulfate 0.1 M MES 30 % w/v PEG 5000 MME, pH 6.5 | 2:1 |
| sc-apCC3-CW-2NTT | ProPlex, G12 | 1.0 M Sodium citrate tribasic dihydrate 0.1 M Sodium HEPES, pH 7.0 | 2:1 |
| sc-apCC3-ACW-mismatch | JCSG, H3 | 0.1 M BIS-Tris, 25 % *w/v* PEG 3350, pH 5.5 | 2:1:1 |

Table S8. Merging and refinement statistics for X-ray structures

| | sc-apCC3-CW1 | sc-apCC3-CW2 | sc-apCCC3-ACW | sc-apCC3-CW-0NTT | sc-apCC3-CW-2NTT |
|---|---|---|---|---|---|
| PDB IDs | 9rgv | 9rgw | 9rgx | 9rgy | 9rgz |
| Wavelength (Å) | 0.9537 (13,000 eV) | 0.9537 (13,000 eV) | 0.6199 (20,001 eV) | 0.6199 (20,001 eV) | 0.9762 (12,701 eV) |
| Resolution range | 36.82 - 2.1 (2.175 - 2.1) | 50.18 - 2.15 (2.227 - 2.15) | 36.97 - 2.2 (2.279 - 2.2) | 32.22 - 1.9 (1.968 - 1.9) | 40.12 - 2.05 (2.123 - 2.05) |
| Space group | C 1 2 1 | P 41 21 2 | P 21 21 21 | C 1 2 1 | I 1 2 1 |
| Unit cell | 95.832 37.866 77.432 90 122.206 90 | 63.358 63.358 246.633 90 90 90 | 37.9161 37.9161 147.87 90 90 90 | 107.409 37.1196 63.4334 90 112.617 90 | 72.7187 37.2862 83.821 90 106.83 90 |
| Total reflections | 26286 (2674) | 56597 (5544) | 144843 (14753) | 125686 (12758) | 89967 (8606) |
| Unique reflections | 13624 (1380) | 28306 (2772) | 11473 (1126) | 18511 (1829) | 13719 (1345) |
| Multiplicity | 1.9 (1.9) | 2.0 (2.0) | 12.6 (13.1) | 6.8 (7.0) | 6.6 (6.4) |
| Completeness (%) | 97.09 (99.78) | 99.38 (99.96) | 99.98 (100.00) | 99.92 (99.95) | 99.59 (99.63) |
| Mean I/sigma(I) | 10.89 (1.96) | 32.56 (2.23) | 8.87 (3.08) | 10.34 (1.61) | 13.56 (3.56) |
| Wilson B-factor ($Å^2$) | 42.23 | 53.18 | 25.90 | 23.15 | 33.94 |
| R-merge | 0.0255 (0.306) | 0.007184 (0.2151) | 0.2102 (0.4033) | 0.1007 (0.3515) | 0.06717 (0.4526) |
| R-meas | 0.03607 (0.4327) | 0.01016 (0.3042) | 0.2193 (0.4196) | 0.1091 (0.3802) | 0.07284 (0.4927) |
| R-pim | 0.0255 (0.306) | 0.007184 (0.2151) | 0.06169 (0.1149) | 0.04144 (0.144) | 0.02786 (0.1923) |
| CC1/2 | 0.999 (0.845) | 1 (0.886) | 0.99 (0.885) | 0.997 (0.938) | 0.999 (0.941) |
| CC* | 1 (0.957) | 1 (0.969) | 0.997 (0.969) | 0.999 (0.984) | 1 (0.985) |
| Reflections used in refinement | 13600 (1378) | 28302 (2771) | 11473 (1126) | 18502 (1828) | 13708 (1344) |
| Reflections used for R-free | 705 (61) | 1403 (143) | 621 (70) | 923 (89) | 660 (65) |
| R-work | 0.2101 (0.2789) | 0.2233 (0.2700) | 0.2196 (0.2675) | 0.1814 (0.2358) | 0.2164 (0.2827) |
| R-free | 0.2506 (0.3372) | 0.2387 (0.3123) | 0.2419 (0.2430) | 0.2099 (0.2859) | 0.2487 (0.3633) |

| | | | | | |
|---|---|---|---|---|---|
| CC(work) | 0.971 (0.880) | 0.965 (0.834) | 0.876 (0.460) | 0.960 (0.909) | 0.966 (0.895) |
| CC(free) | 0.977 (0.662) | 0.970 (0.774) | 0.790 (0.564) | 0.946 (0.852) | 0.981 (0.717) |
| Number of non-hydrogen atoms | 1534 | 2264 | 1371 | 1789 | 1495 |
| macromolecules | 1384 | 2164 | 1289 | 1414 | 1353 |
| ligands | 0 | 11 | 0 | 0 | 0 |
| solvent | 150 | 89 | 82 | 375 | 142 |
| Protein residues | 196 | 300 | 192 | 193 | 195 |
| RMS(bonds) | 0.003 | 0.002 | 0.002 | 0.008 | 0.003 |
| RMS(angles) | 0.61 | 0.41 | 0.45 | 0.76 | 0.58 |
| Ramachandran favored (%) | 100 | 100 | 97.31 | 100 | 100 |
| Ramachandran allowed (%) | 0 | 0 | 2.69 | 0 | 0 |
| Ramachandran outliers (%) | 0 | 0 | 0 | 0 | 0 |
| Rotamer outliers (%) | 0 | 1.12 | 4.65 | 0 | 0 |
| Clashscore | 3.58 | 0.91 | 4.75 | 5.14 | 4.84 |
| Average B-factor | 52.67 | 66.04 | 26.77 | 29.79 | 43.6 |
| macromolecules | 52.62 | 66.09 | 26.83 | 27.97 | 43.34 |
| Ligand | - | 96.6 | - | - | - |
| solvent | 53.19 | 61.1 | 25.5 | 36.68 | 46.09 |
| Number of TLS groups | 2 | 10 | 2 | 2 | 2 |

Table S9. Comparison of helicity predicted by DSSP analysis of AlphaFold2 models and experimentally measured helicity by CD spectroscopy across 3 repeats

| protein name | DSSP calculated % helicity | experimentally measured % helicity ($N$ = 3) |
|---|---|---|
| sc-apCC3-CW1 | 78.45 | 78.45 ± 2.41 |
| sc-apCC3-CW-flex | 76.52 | 68.72 ± 2.55 |
| sc-apCC3-CW-mismatch | 78.26 | 57.07 ± 2.24 |
| sc-apCC3-ACW | 75.65 | 82.21 ± 1.79 |
| sc-apCC3-ACW-flex | 73.91 | 70.20 ± 1.96 |
| sc-apCC3-ACW-mismatch | 75.00 | 53.50 ± 1.51 |

Table S10. Summary of thermodynamic analyses

| [GdmHCl] (M) | % helicity | Fitted $T_M$ (°C) |
|---|---|---|
| 0.0 | 75.1 | 87.0 |
| 0.2 | 58.0 | 88.9 |
| 0.3 | 54.4 | 83.5 |
| 0.5 | 45.7 | 71.7 |
| 0.8 | 46.3 | 63.5 |
| 1.0 | 44.1 | 56.9 |
| 1.4 | 32.2 | 43.4 |
| 1.5 | 34.9 | 44.2 |
| 1.8 | 22.1 | 39.1 |

Globally fitted parameters:

| | |
|---|---|
| $\Delta G$ (kcal mol$^{-1}$) | 6.16 ± 0.07 |
| $\Delta H$ (kcal mol$^{-1}$) | 24.08 ± 0.53 |
| $\Delta S$ (kcal mol$^{-1}$ K$^{-1}$) | 0.06 ± 0.00 |
| $\Delta C_p$ (kcal mol$^{-1}$ K$^{-1}$) | 0.19 ± 0.01 |
| $\Delta\Delta C_p$ (kcal mol$^{-1}$ K$^{-1}$ M$^{-1}$) | 0.46 ± 0.01 |
| m (kcal mol$^{-1}$ M$^{-1}$) | -3.22 ± 0.03 |

### 1.8 Supplementary figures

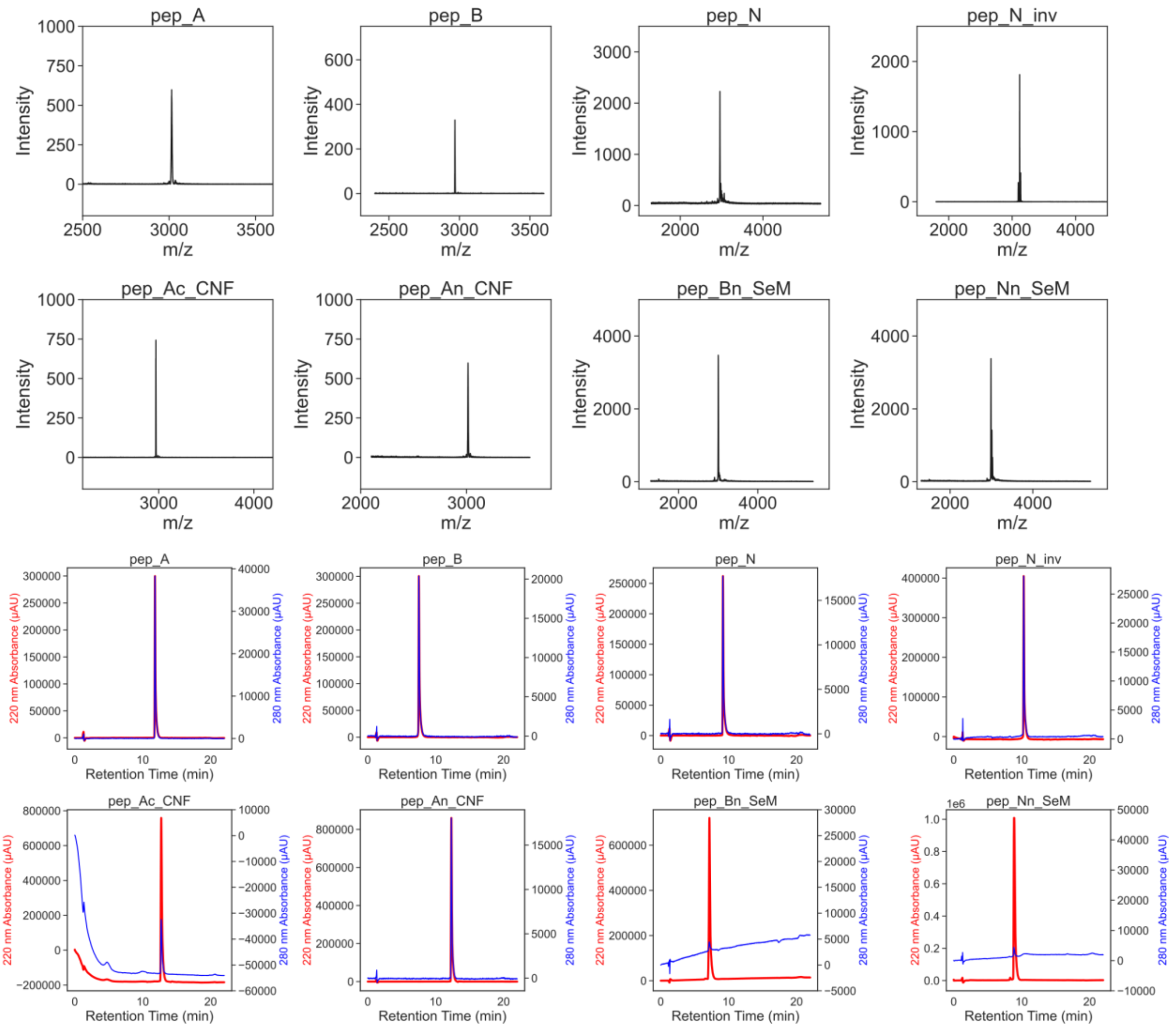

Fig. S1 MALDI-TOF spectra (top) and analytical HPLC traces (bottom) of *de novo* peptides (blue – 280 nm, red – 220 nm)

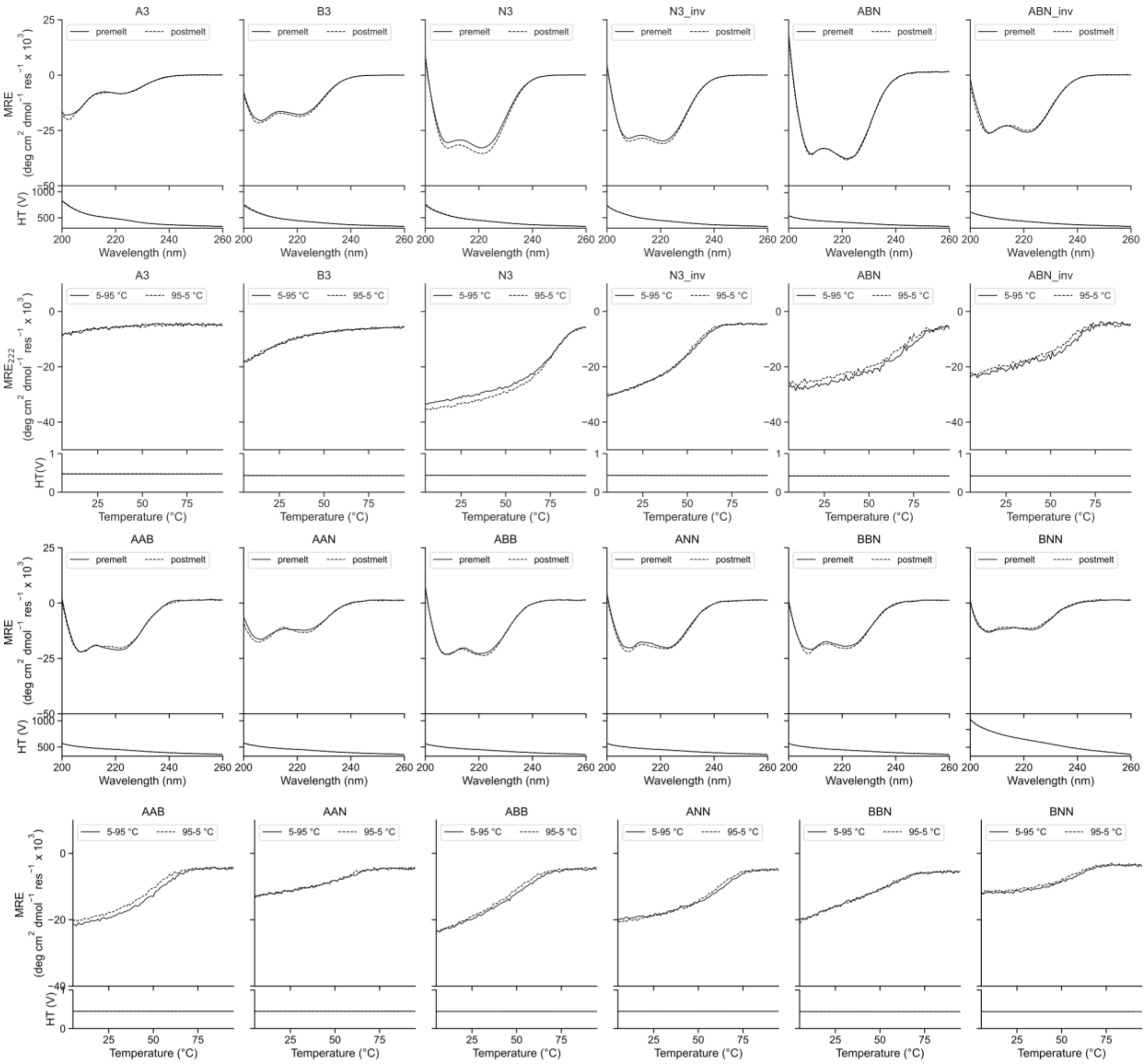

Fig. S2 CD spectra of the *de novo* peptides and the off-target mixtures

Sequences for individual peptides can be found in Supplementary Table 3. Summary of biophysical characterization can be found in Supplementary Table 6. Conditions: 100 μM overall peptide assembly, PBS, pH 7.4, 5 to 95 °C. CD scans before the thermal ramps up (solid line) and after thermal ramps down (dashed line).

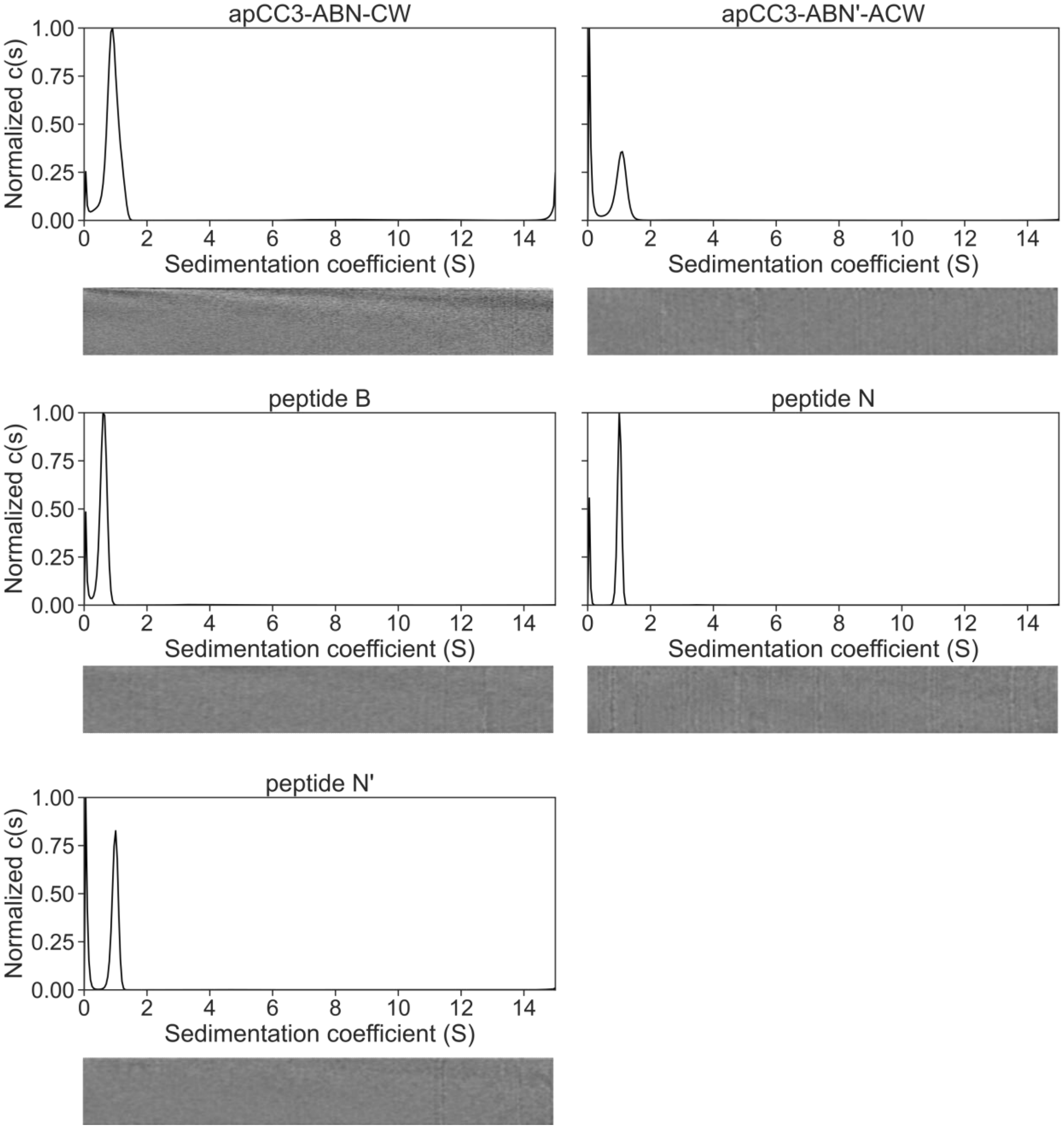

Fig. S3 AUC-SV traces of the *de novo* peptides

Sequences for individual peptides can be found in Supplementary Table 3. Summary of biophysical characterization can be found in Supplementary Table 6. Residuals are shown as bitmaps below the fitted data. Conditions: 250 μM overall peptide assembly, PBS, pH 7.4, 20 °C, 60 krpm.

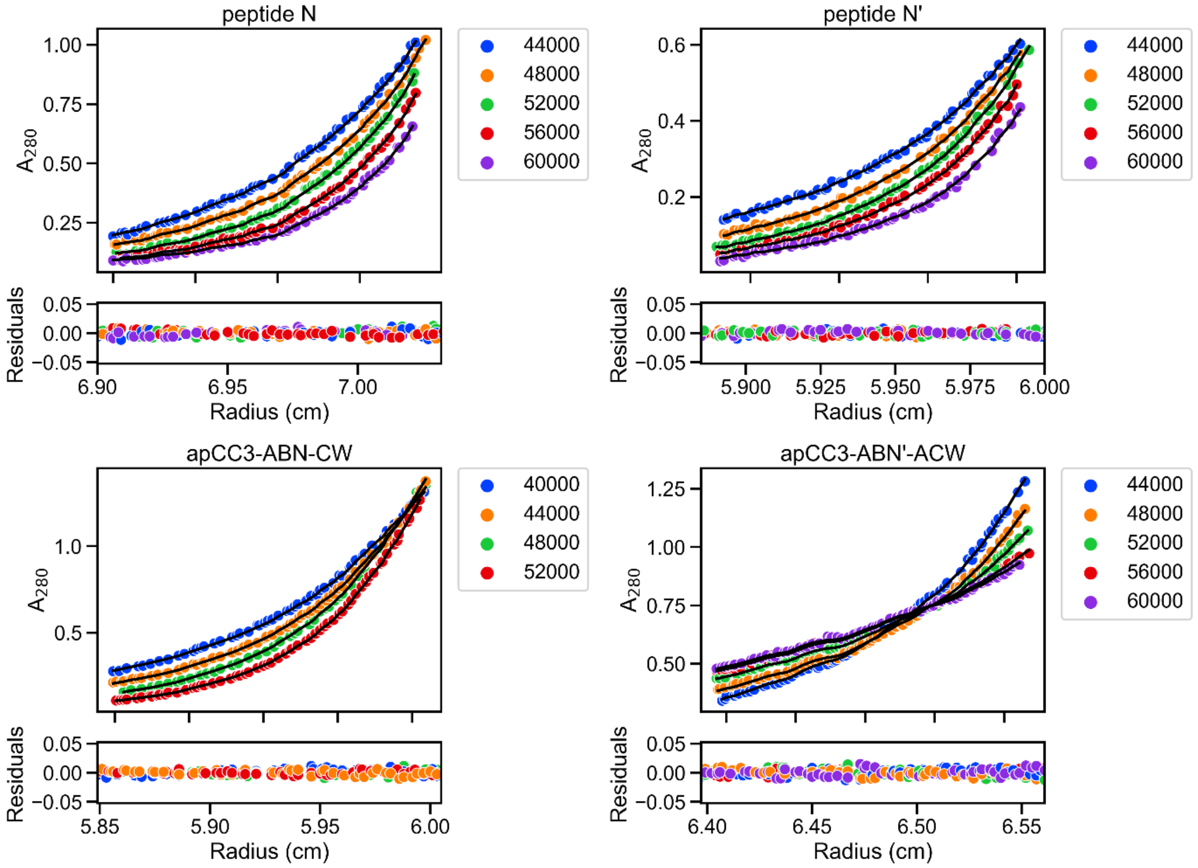

Fig. S4 AUC-SE traces of the *de novo* peptides

Sequences for individual peptides can be found in Supplementary Table 3. Summary of biophysical characterization can be found in Supplementary Table 6. Residuals are shown as residual maps below the fitted data. Conditions: 250 µM overall peptide assembly, PBS, pH 7.4, 20 °C, 60 krpm.

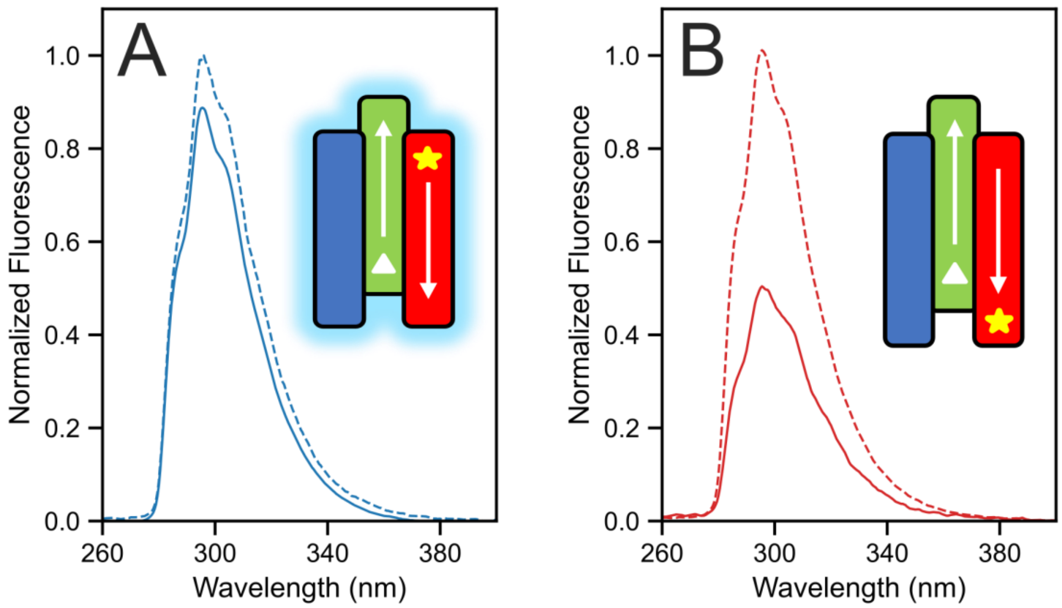

Fig. S5 Fluorescence-quenching assays for mixtures of apCC3-N-nMSE and apCC3-B plus apCC3-A-n4CF (A) or apCC3-A-c4CF (B). Dotted lines are for the fluorescently-labelled A peptides alone, and solid lines are for the 1:1:1 mixture. In the cartoons, arrows indicate the peptide direction from *N* to *C* termini; and 4CF is represented by the yellow star, and MSE by the white triangle. Conditions: 33 µM of each peptide in phosphate buffer (8.2 mM potassium phosphate dibasic, 1.8 mM potassium phosphate monobasic), pH 7.4.

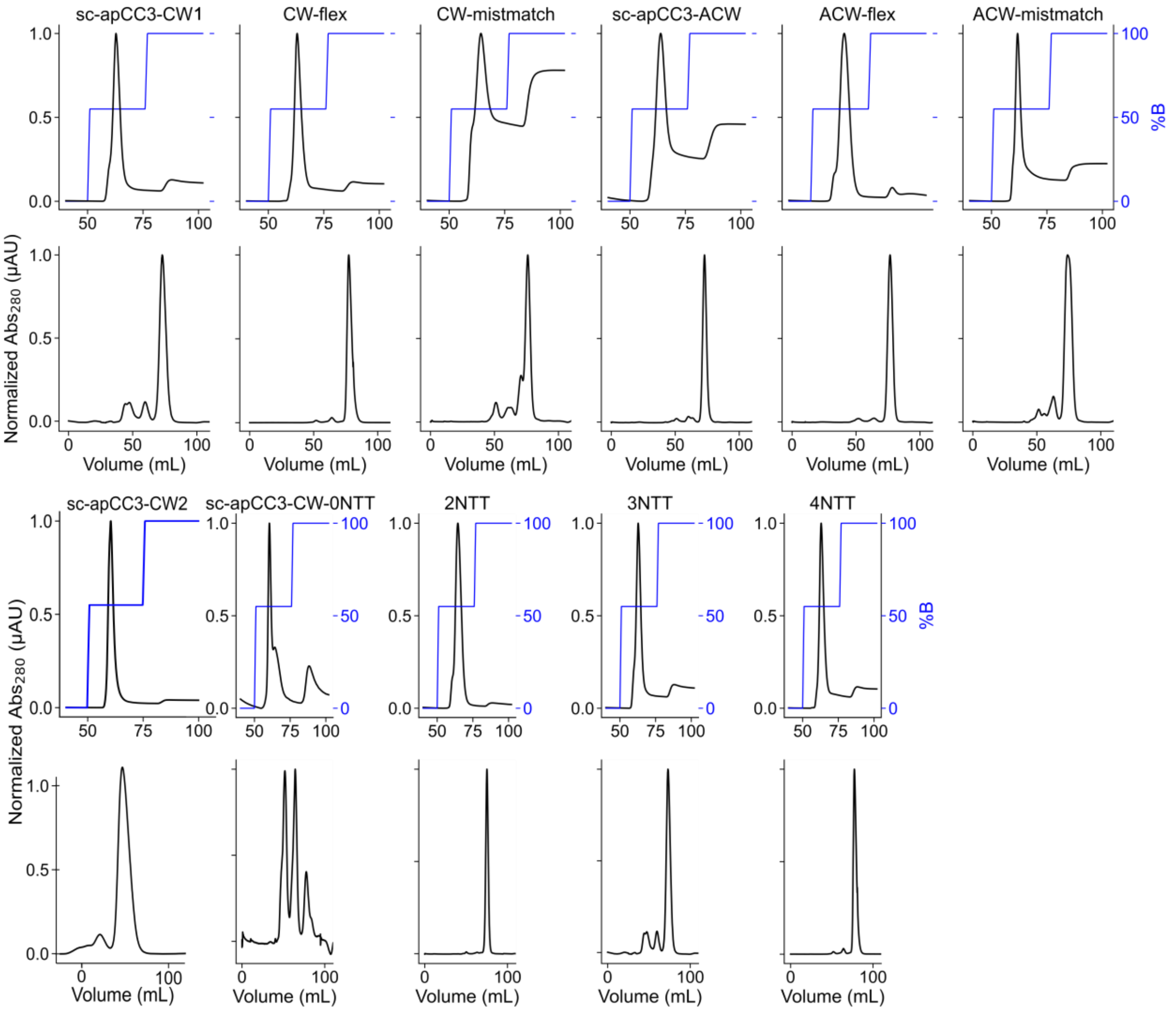

**Fig. S6 FPLC purification of sc-apCC3 and its variants**
Sequences for individual proteins can be found in Supplementary Table 3. Top: Ni-affinity chromatography. Fractions eluting at 55% B were pooled and concentrated then further purified by size-exclusion chromatography. Conditions: Buffer A: 50 mM Tris, pH 7.4, 500 mM NaCl, 30 mM imidazole, B: 300 mM imidazole. Bottom: Size-exclusion chromatography. Fractions eluting between 75-90 mL were pooled and concentrated for further biophysical analysis. Conditions: 50 mM sodium phosphate, pH 7.4, 150 mM NaCl.

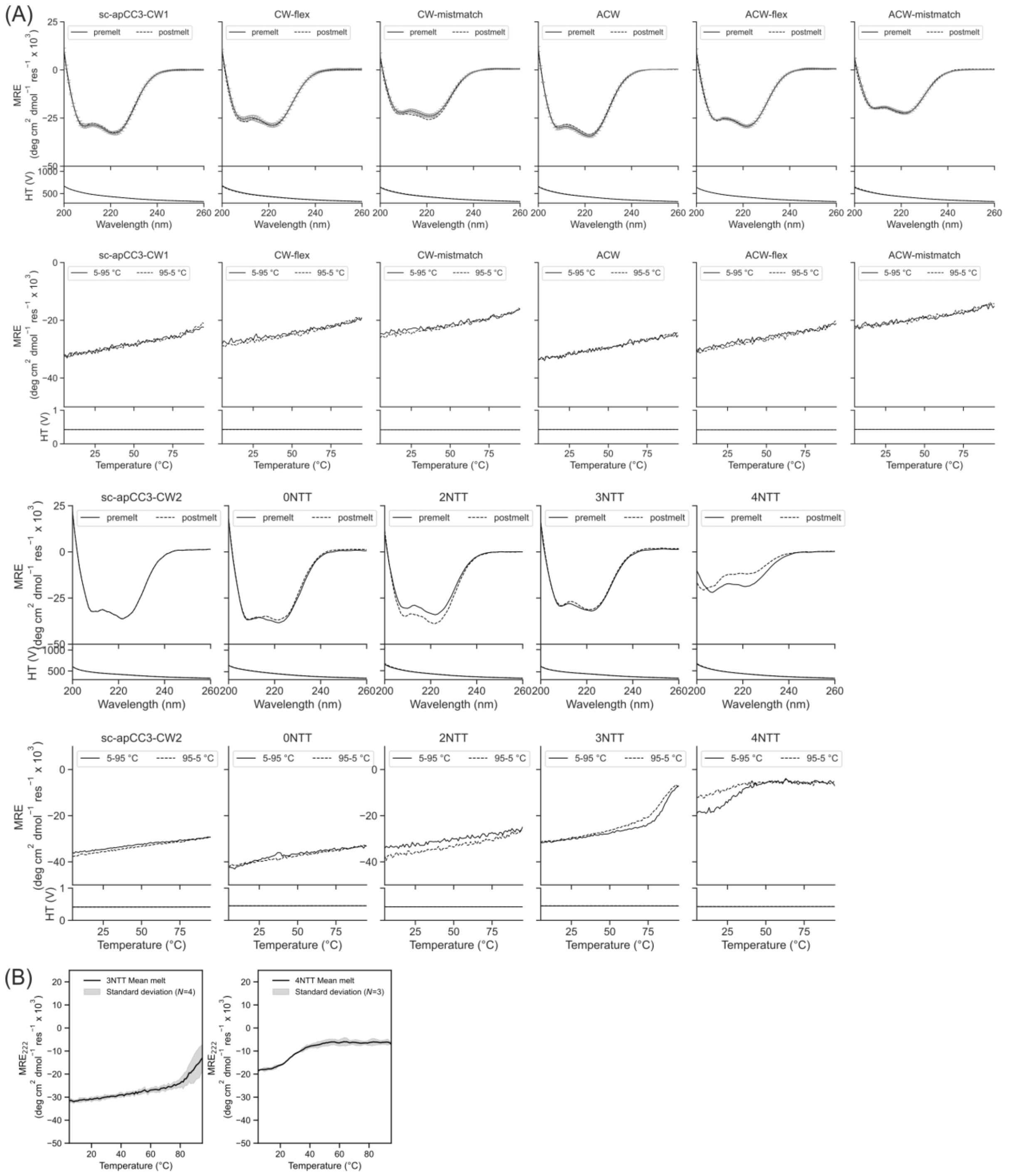

Fig. S7 CD spectra of the *de novo* proteins

Sequences for individual proteins can be found in Supplementary Table 3. Summary of biophysical characterization can be found in Supplementary Table 6. (A) Conditions: 10 µM protein, 50 µM sodium phosphate, pH 7.4, 150 mM NaCl, top – 5 °C, error bars represent one standard deviation of the mean, *N*=3 for all loop variants; bottom – thermal response 5 to 95 °C. CD scans before the thermal ramps up (solid line) and after thermal ramps down (dashed line). (B) The filled regions represent one standard deviation of the mean; *N*=4 for 3NTT variant, *N*=3 for 4NTT variant**.**

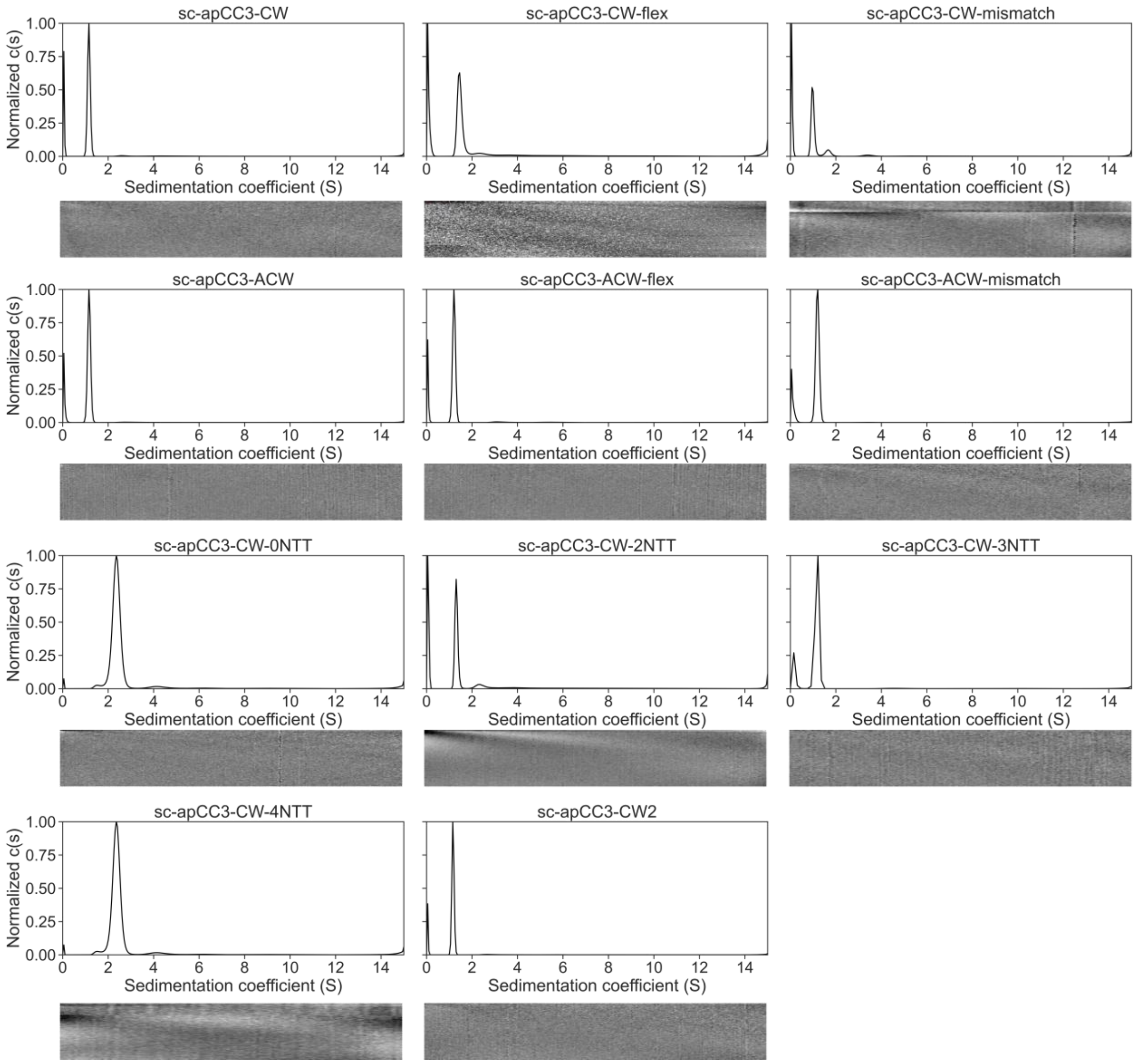

Fig. S8 AUC-SV traces of the *de novo* proteins

Sequences for individual proteins can be found in Supplementary Table 3. Summary of biophysical characterization can be found in Supplementary Table 6. Residuals are shown as bitmaps below the fitted data. Conditions: 50 μM protein, 50 μM sodium phosphate, pH 7.4, 150 mM NaCl, 20 °C, 60 krpm.

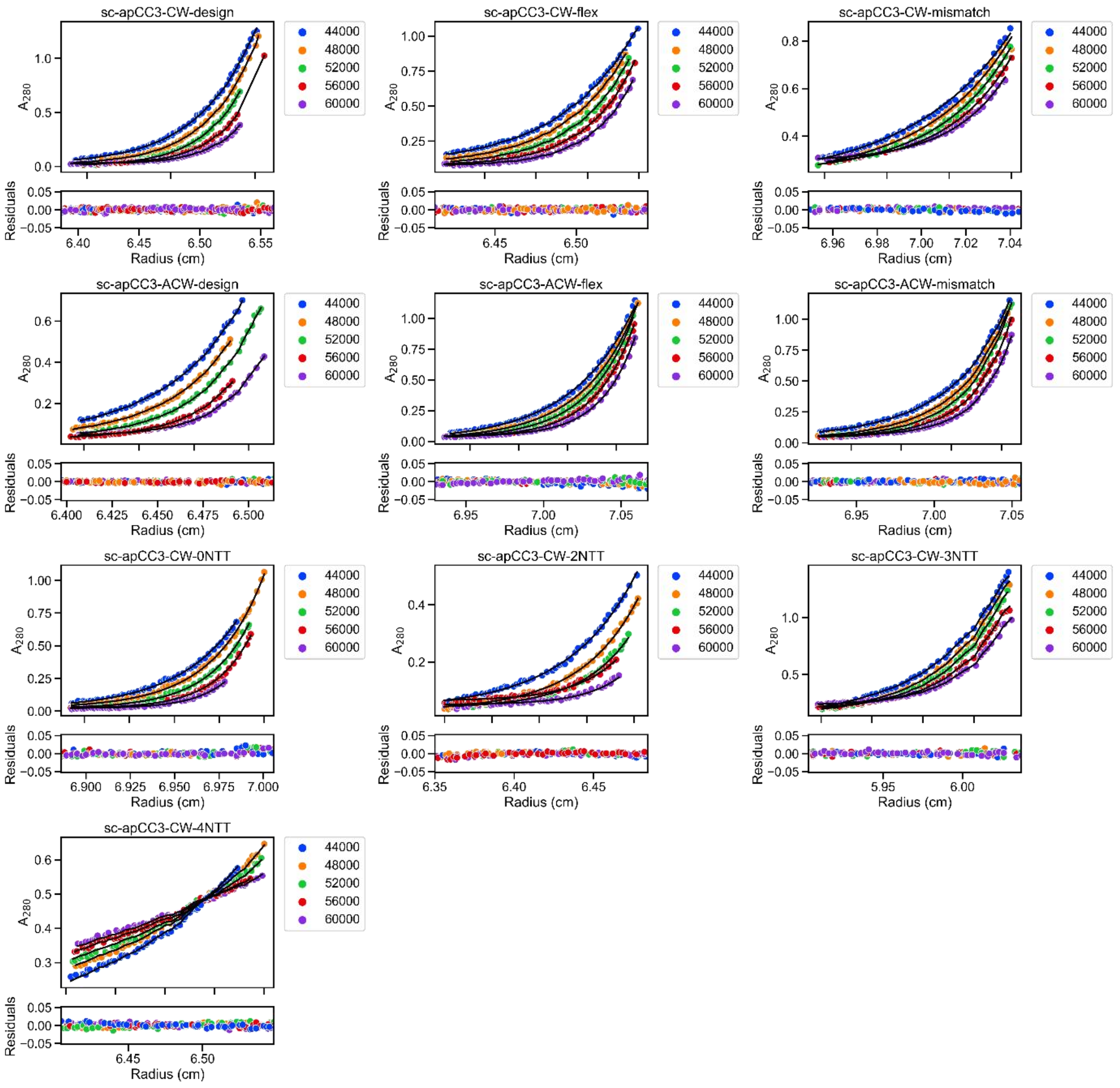

Fig. S9 AUC-SE traces of the *de novo* proteins

Sequences for individual proteins can be found in Supplementary Table 3. Summary of biophysical characterization can be found in Supplementary Table 6. Residuals are shown as residual maps below the fitted data. Conditions: 50 μM protein, 50 μM sodium phosphate, pH 7.4, 150 mM NaCl, 20 °C, 44-60 krpm with 4 krpm increment at 8 h intervals.

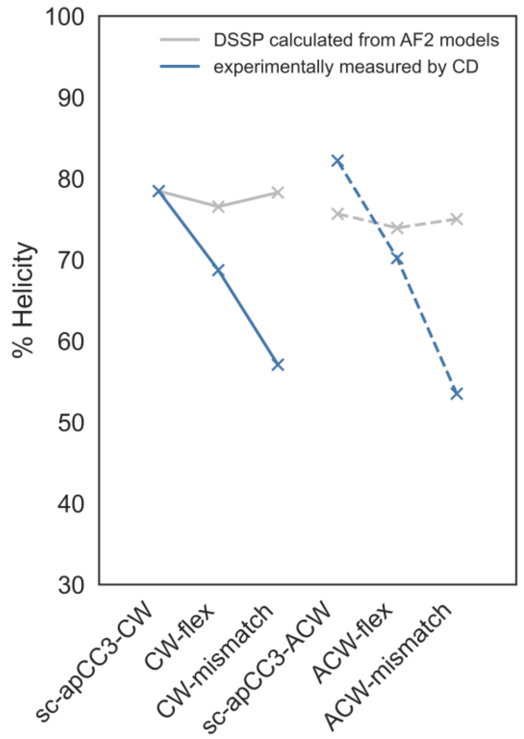

Fig. S10 Comparison of helicity predicted by DSSP analysis of AlphaFold2 models (gray) and experimentally measured helicity by CD spectroscopy (blue) across flexible and mismatched loop constructs. Key: variants with CW core sequences – solid lines, variants with ACW core sequences – dotted lines. Results for calculated helicities can be found in Supplementary Table S9.

sc-apCC3-CW1

Rank 1

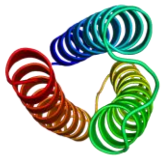

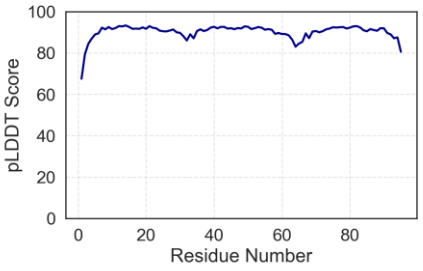

Rank 2

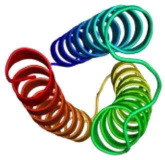

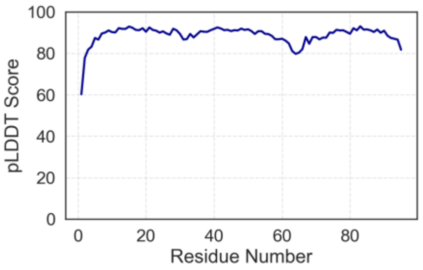

Rank 3

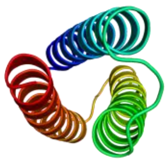

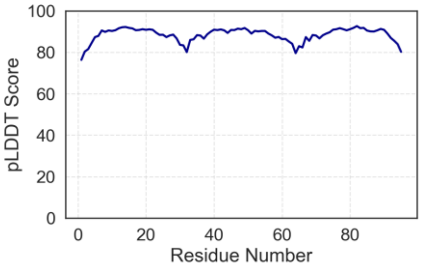

Rank 4

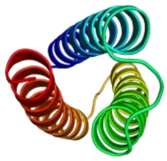

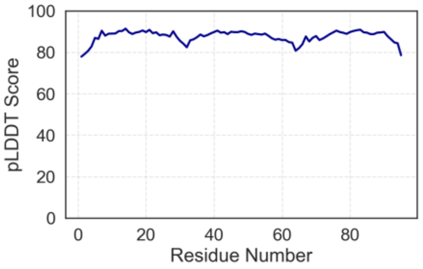

Rank 5

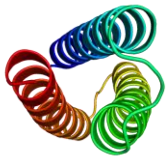

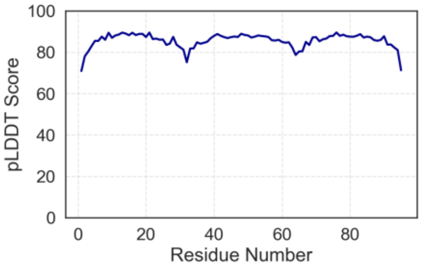

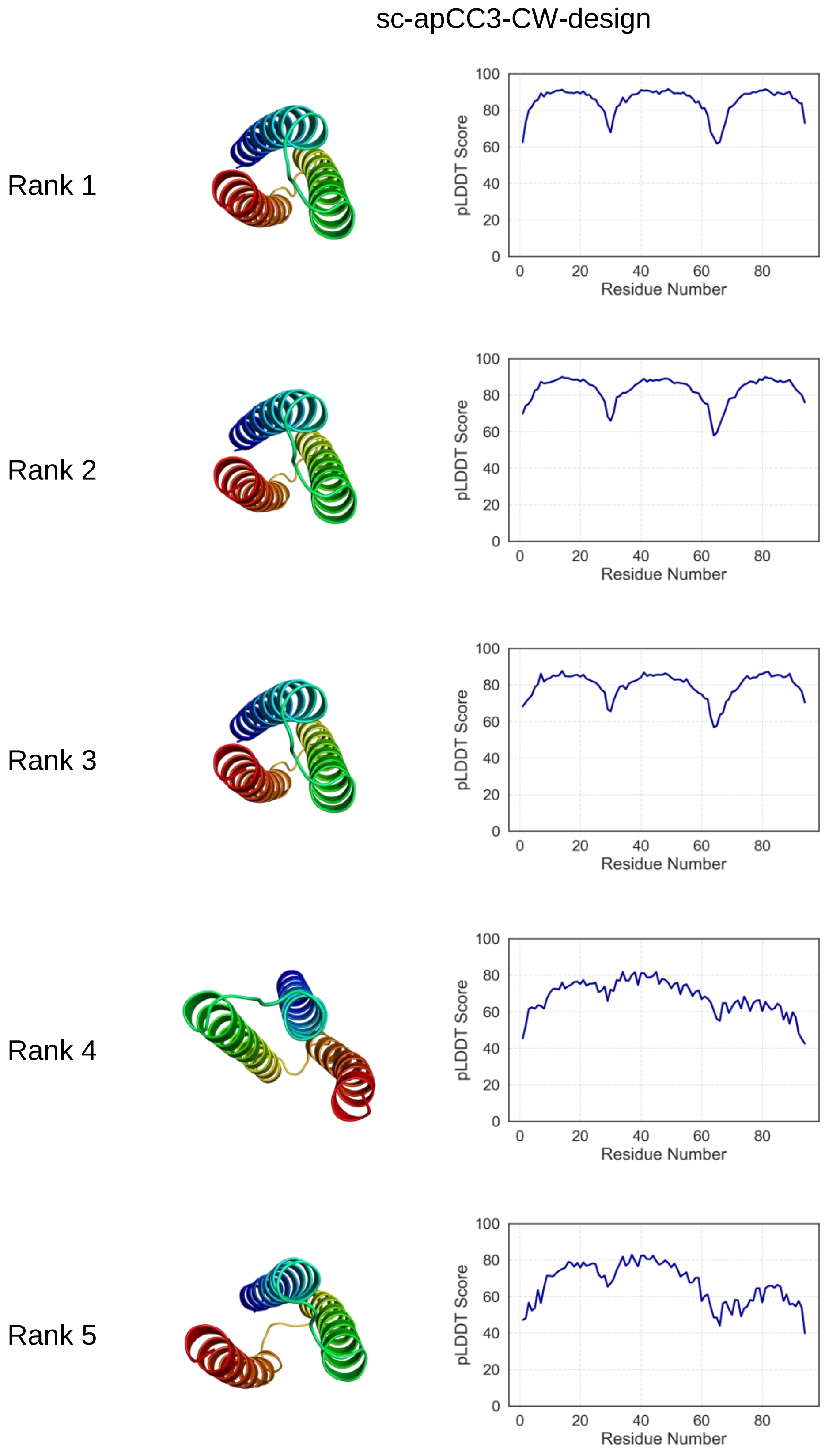

Fig. S11 AF2 predictions reveal potential access to alternative folds in mismatched constructs. Top-down views and pLDDT scores of AF2-predicted models for sc-apCC3-CW1 (top) all show consistent adoption of CW topology with high confidence score, whereas rank 4 and 5 predictions of sc-apCC3-CW-mismatch (bottom) show access to ACW topology and unfolding of the 3HB bundle, respectively. Mismatched sequences may challenge accurate global fold prediction.

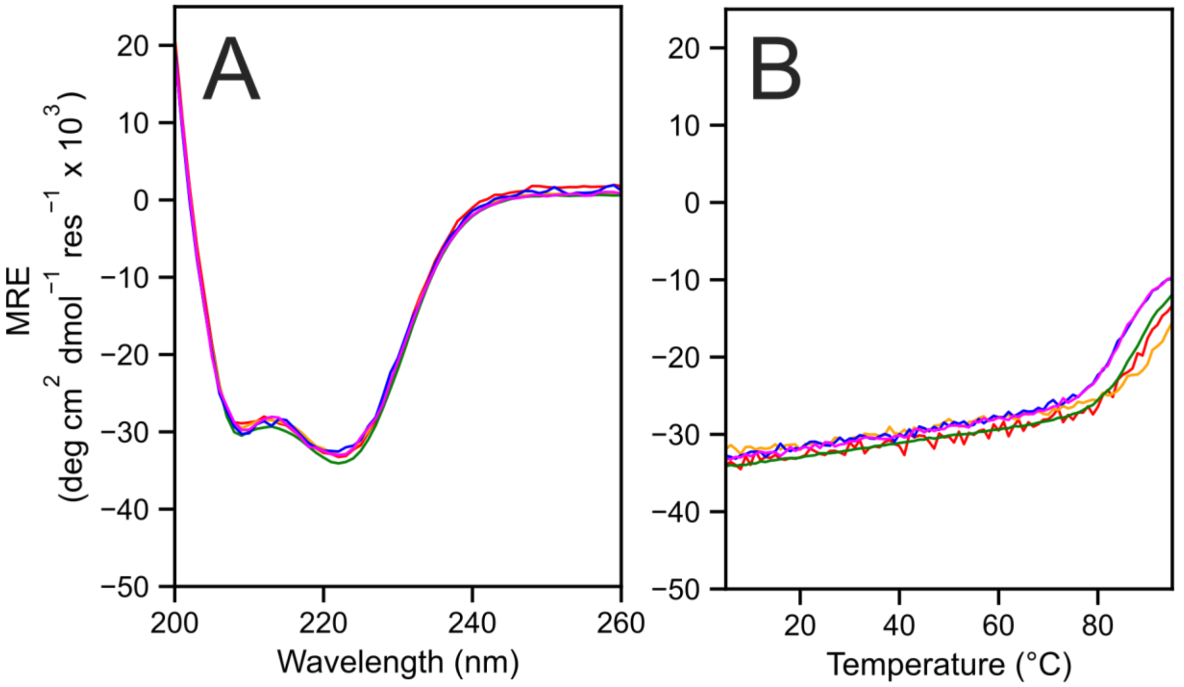

Fig. S12 Concentration-independent CD measurements of sc-apCC3-CW-3NTT. Conditions: 50 µM sodium phosphate, pH 7.4, 150 mM NaCl, (A) 5 °C, (B) thermal response 5 to 95 °C. Key: 5 µM – red, 10 µM – yellow, 20 µM – green, 30 µM – blue, 45 µM – magenta.

Thermal denaturation modelling and thermodynamic analysis using van't Hoff equations

Each thermal denaturation data was fitted to a two-state folding model, assuming there are only folded and unfolded states due to a sharp sigmoidal transition. The two-state folding thermodynamic parameters were extracted as described by Gellman[11] using Equations 1-9 and the following parameters.

| | |
|---|---|
| T | absolute temperature |
| $\theta_f$ | ellipticity contribution from folded conformation |
| $\theta_u$ | ellipticity contribution from unfolded conformation |
| $\Delta H$ | enthalpy change at a given temperature |
| $\Delta S$ | entropy change at a given temperature |
| K | equilibrium constant |
| $b_f$ | y-intercept of folded baseline |
| α | fraction folded conformation ($0 \leq \alpha \leq 1$) |
| $\Delta G$ | Gibbs free energy change |
| $\Delta C_p$ | heat capacity change of folding |
| $\theta_T$ | measured ellipticity |
| M | melting temperature |
| $T_r$ | reference temperature |
| $m_f$ | slope of folded baseline |
| x | temperature |
| $\Delta\Delta C_p$ | Denaturant concentration dependence of heat capacity of folding |
| R | universal gas constant |
| $b_u$ | y-intercept of unfolded baseline |

Equation 1 shows how the observed ellipticity $\theta_T$ is composed of a combination of signals from both the folded and unfolded states. The melting curve function models the temperature dependent transition between the two states by calculating the fraction folded conformation at a given temperature x using a sigmoidal function, Equations 2 and 3. The sigmoidal function describes how the folding transition occurs as the temperature changes, driven by enthalpy-like steepness term *h* and melting temperature M. The function uses a weighted sum of signals from both states, $b_f$ and $b_u$, and $m_f$ describes a linear slope at the folded state (Figure S13). The fraction of folded conformation, α, is defined by rearranging the equilibrium constant of folding (Equation 4, Figure S14) in terms of the Gibbs free energy (Equations 5 and 6). This relationship forms the basis for the subsequent thermodynamic models.

$$\theta_T = \alpha\theta_f + (1-\alpha)\theta_u \quad (1)$$

$$y(x) = f(x) \cdot (b_f - b_u + m_f x) + b_u \quad (2),$$

where

$$f(x) = \frac{1}{1 + \exp\left(-\frac{h\left(1-\frac{x}{M}\right)}{x}\right)} \quad (3)$$

$$\alpha = \frac{1}{1+K} \quad (4)$$

$$K = e^{-\frac{\Delta G}{RT}} \quad (5)$$

$$\ln K = -\Delta G(T) \quad (6)$$

To characterize the thermal stability of the system, we fitted all unfolding transitions assuming the following dependences. The ellipticities of folded and unfolded states linearly depend on temperature and concentration of denaturant concentration (Equations 7 and 8). The Gibbs free energy of the native state is determined by equation 9 (Fig S15, Table S10).[12, 13]

$$\theta_f = b_f + m_f T + a[\mathrm{GdmHCl}] \quad (7)$$

$$\theta_u = b_u + m_u T + b[\mathrm{GdmHCl}] \quad (8)$$

$$\Delta G(T, [GdmHCl]) = \Delta H_0 - T\Delta S_0 + (\Delta C_{p,0} + \Delta\Delta C_p \cdot [\mathrm{GdmHCl}])\left(T - T_0 + T\ln\left(\frac{T_0}{T}\right)\right) + m \cdot [\mathrm{GdmHCl}] \quad (9)$$

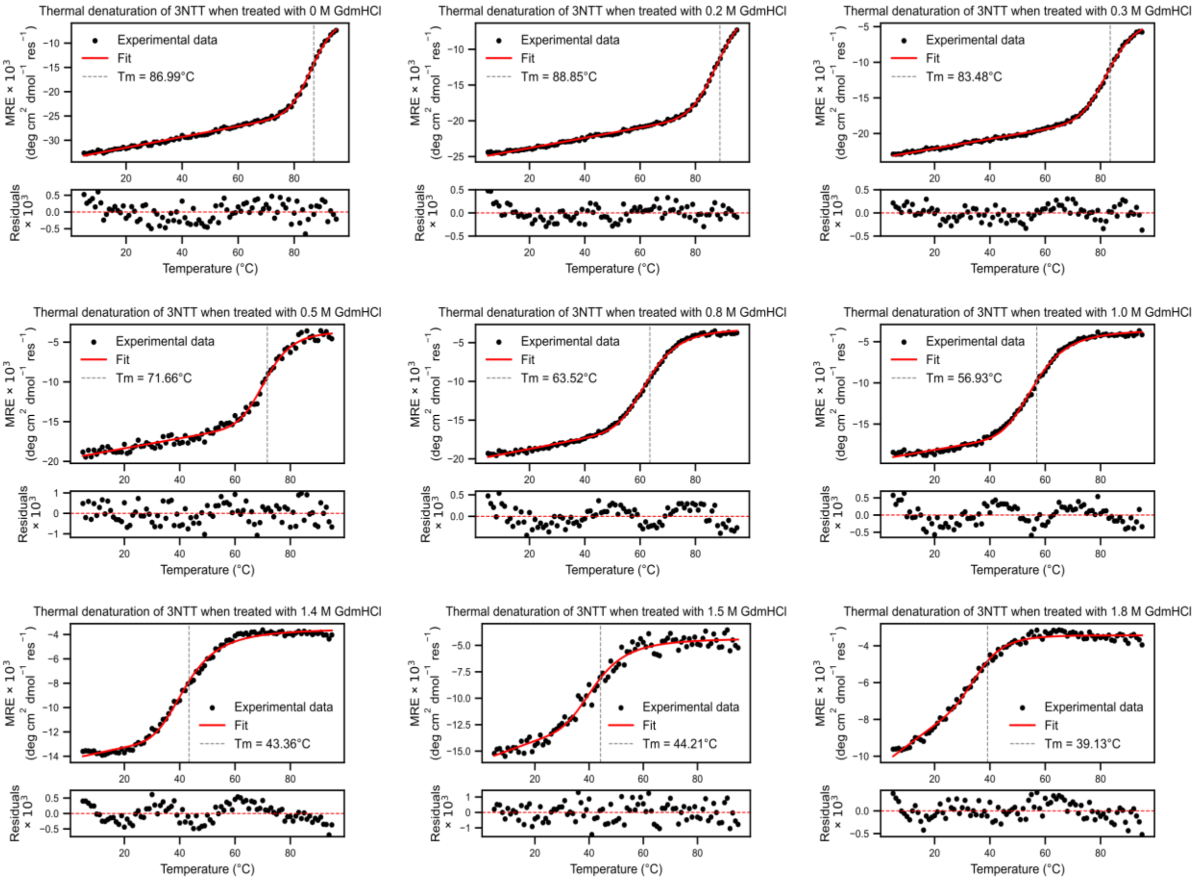

Fig. S13 Variable temperature CD signal of sc-apCC3-CW-3NTT when treated with GdmHCl concentrations of 0, 0.2, 0.3, 0.5, 0.8, 1.0, 1.4, 1.5, 1.8 M, monitored at 222 nm, 5 to 95 °C. Data was fitted to a two-state folding model (top) for melting temperature calculation and fit residual (bottom) using Equations 2 and 3. Results for individually fitted $T_M$ can be found in Supplementary Table S10.

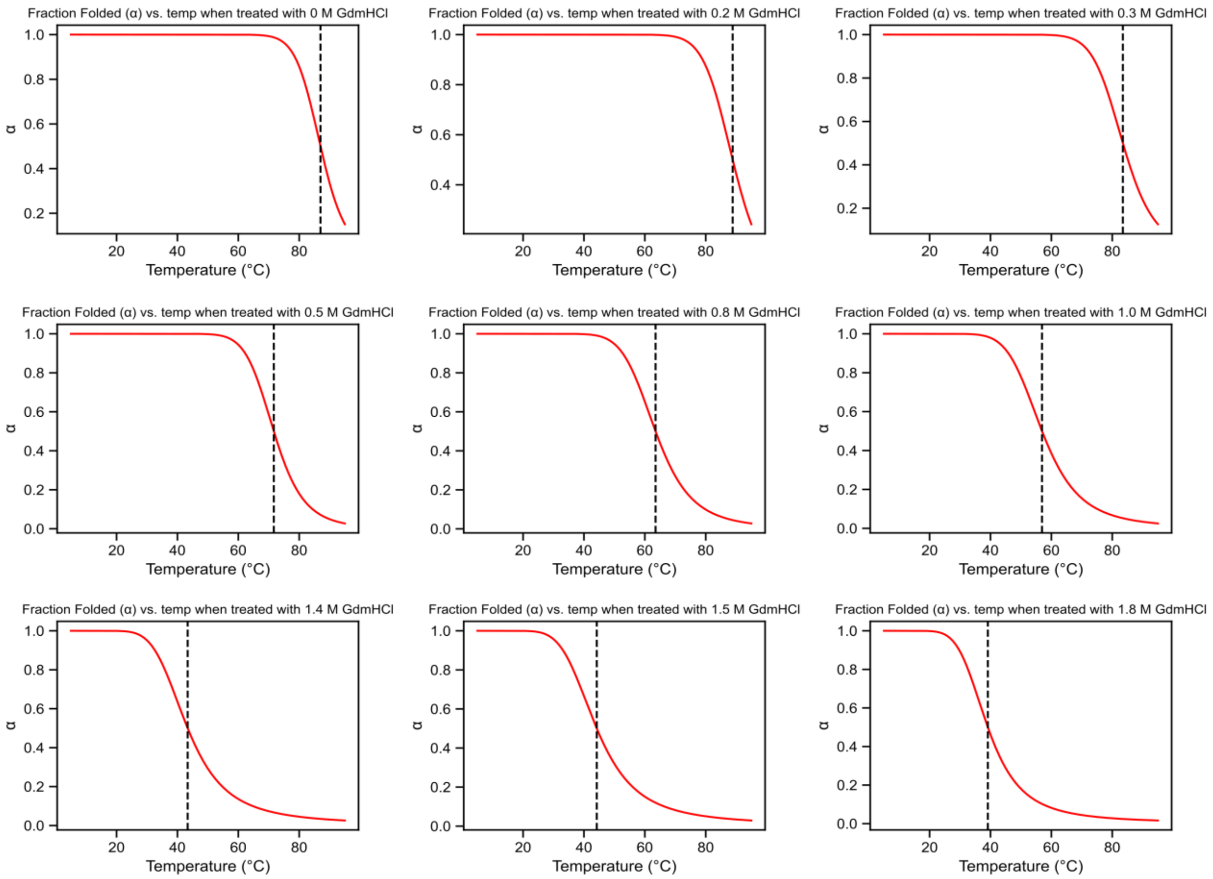

Fig. S14 Fraction folded (α) of sc-apCC3-3NTT unfolding as a function of temperature calculated using Equation 4. The dotted line represents the transition midpoint temperature when α = 0.5.

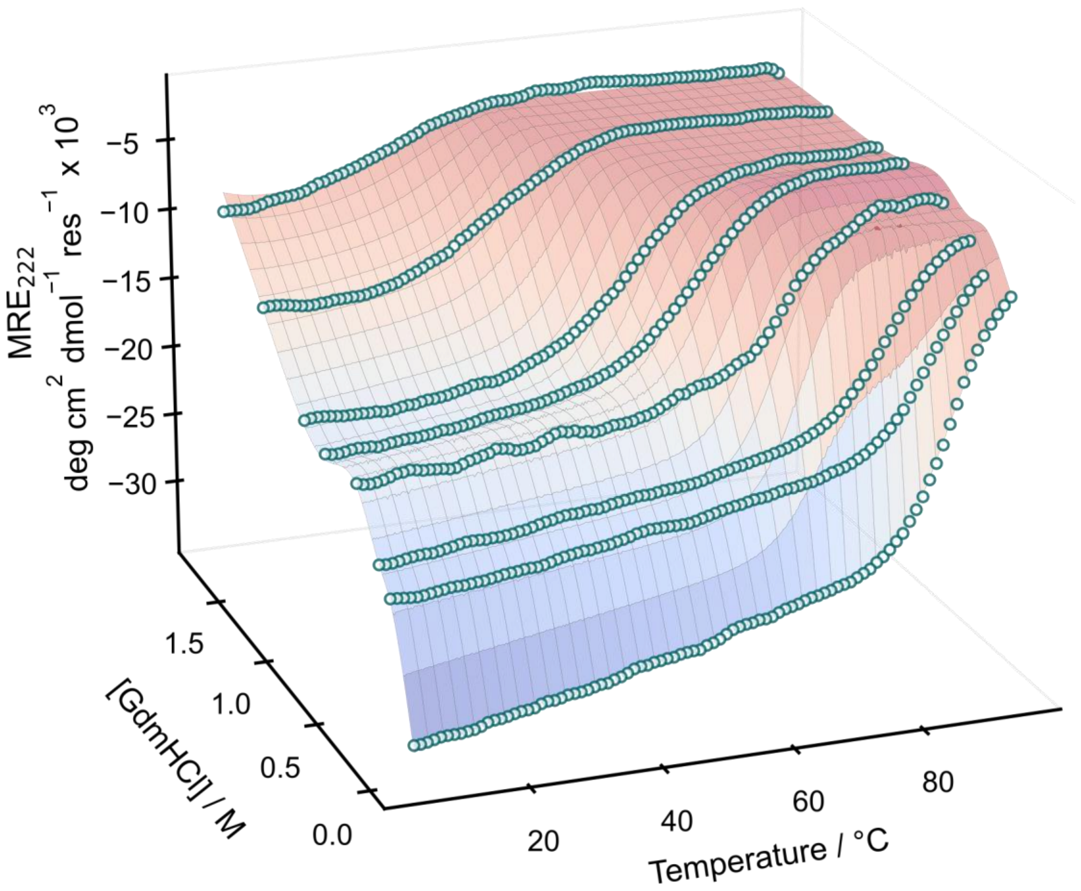

Fig. S15 MRE at 222 nm of sc-apCC3-3NTT unfolding as a function of temperature and GdmHCl concentration. Conditions: 10 μM protein, 50 μM sodium phosphate, 150 mM NaCl, pH 7.4, 5 to 95 °C; treated with GdmHCl concentrations of 0, 0.2, 0.3, 0.5, 0.8, 1.0, 1.4, 1.5, 1.8 M. Raw data are shown as blue scatters, and the globally-fitted thermodynamic parameters are represented by the surface. Errors are reported as fitting errors. Thermodynamic parameters were extracted using Equation 9 and results can be found in Supplementary Table S10.

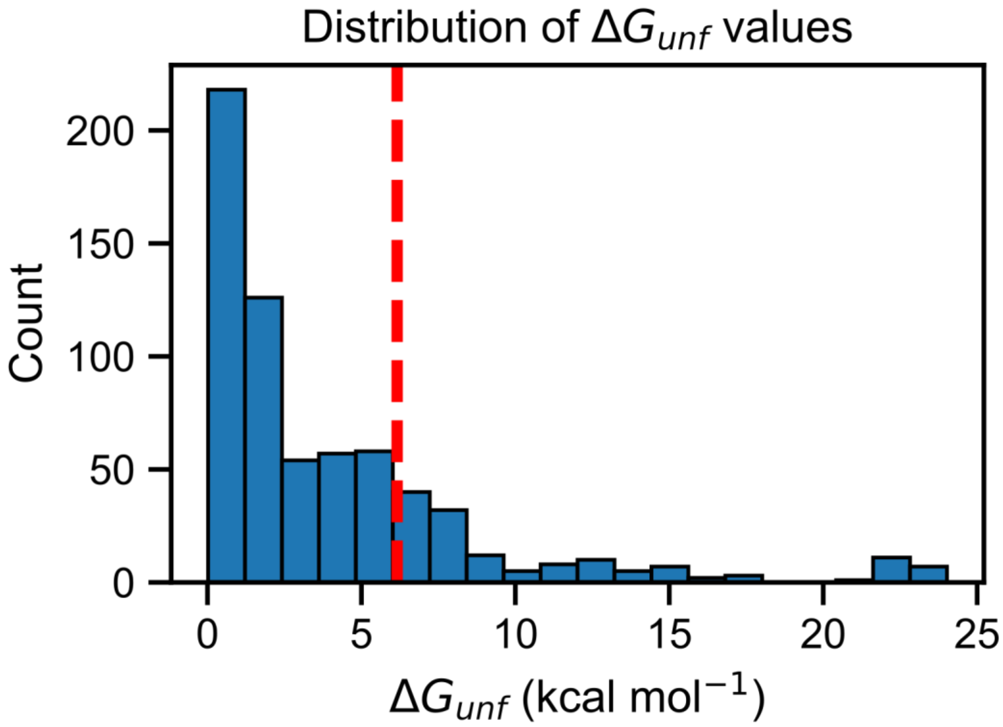

Fig. S16 Histogram of $\Delta G_{unf}$ in the ProTherm protein stability database[14] (search limited to helix, GdmHCl, yielded 656 examples). The estimated $\Delta G_{unf}$ of sc-apCC3-CW-3NTT ($\Delta G_{unf}$ = 6.16 ± 0.07 kcal mol$^{-1}$) reported at reference temperature of 25 °C is indicated by the red dashed line.
